# Loss of ESE3/EHF is sufficient to promote cell plasticity, transformation and androgen-independent status in the early stage of prostate carcinogenesis

**DOI:** 10.1101/2025.11.26.690649

**Authors:** Domenico Albino, Giada Sandrini, Carola Musumeci, Elisa Storelli, Giovanni Papa, Daniela Impellizzieri, Alessia Cacciatore, Giada Andrea Cassanmagnago, Atik Balla, Simone Mosole, Gianluca Civenni, Andrea Rinaldi, Carmen Jeronimo, Rui M. Henrique, Marco Bolis, Carlo V. Catapano, Giuseppina M. Carbone

## Abstract

Phenotypic plasticity enables tumor progression and treatment resistance. However, its timing and underlying mechanisms are poorly understood. Here, we demonstrate that cell plasticity can emerge early during prostate cancer development, resulting from the knockout of the epithelial-specific ETS transcription factor EHF in prostate epithelial cells. Inspecting the transcriptome of human prostate cancers, we identified a correlation between low EHF expression, loss of luminal epithelial identity, and attenuated androgen signaling in both primary tumors and castration-resistant prostate cancers (CRPC). In EHF knockout mouse models and human epithelial cells, EHF ablation was sufficient to disrupt epithelial cell lineage integrity and promote a progenitor/stem cell-like state with both basal and luminal features, enabling high plasticity and multi-lineage phenotypic transitions. Mechanistically, EHF acted as a central node controlling a hierarchy of transcriptional regulatory factors and downstream signaling pathways (e.g., COL1A1/DDR1, JAK/STAT3), thereby regulating epithelial lineage integrity and restricting stemness and phenotypic transitions. Activation of these downstream pathways, consequent to EHF loss, promoted non-luminal cell features, attenuated androgenic response, and resistance to AR antagonists. Collectively, these data provide novel insights into the causes of phenotypic plasticity and androgen indifference already at the early stages of prostate tumorigenesis and a new perspective on the paths to cancer progression directly relevant to the development of more efficient treatment strategies.

## Introduction

Phenotypic plasticity is emerging as a critical hallmark of cancer^1^. Cancer cell plasticity enables malignant cells to acquire new properties and multiple oncogenic capabilities, promoting adaptation to the microenvironment, invasion, angiogenesis, immune evasion, metastasis, and treatment resistance^1^. Lineage plasticity may occur via direct trans-differentiation from one cell lineage to another or through dedifferentiation, achieving a progenitor/stem cell-like state, followed by differentiation to a distinct cell lineage^2^. However, the factors that enhance cell plasticity and phenotypic transitions, as well as the timing and mechanisms that enable the maintenance or disruption of cell lineage integrity, are largely unknown.

Advanced prostate cancers exhibit enhanced phenotypic plasticity, leading to altered androgen receptor (AR) signaling, AR indifference, neuroendocrine differentiation, and unresponsiveness to androgen deprivation therapy (ADT) and AR antagonists^3,4^. Significantly, metastatic castration-resistant prostate cancer (CRPC) has limited treatment options and remains a leading cause of cancer-related death in men^4,5^. Phenotypic plasticity is considered a late event in the evolution of prostate cancer toward castration resistance^3^. Cancer cell plasticity, accordingly, was shown to arise from cumulative genetic hits in the murine prostate gland and was sustained by the activation of pro-tumorigenic and inflammatory pathways, like JAK/STAT3^6^.

Transcriptional and epigenetic mechanisms are fundamental for lineage determination and plasticity in normal and tumor cells^2^. In this context, ETS family transcription factors (TFs) have essential roles in many cell types, acting as nodal points in cell fate-determining pathways^7^. Despite the structural similarities, individual ETS factors have distinct expression patterns and drive different biological responses^7,8^. The epithelial-specific ETS transcription factor EHF/ESE3 (hereafter EHF) is one of the most expressed TFs in normal prostate epithelium^9^. Although genetic deletions and mutations are rare^10^, EHF expression is frequently reduced in prostate cancers due to epigenetic silencing^9^. We were the first to show that the EHF mRNA level, when compared to the normal prostate, is significantly reduced in primary prostate cancers, ranging from mild to severe reduction^9^. Reduced level of EHF in primary prostate cancer datasets was associated with epithelial-to-mesenchymal transition (EMT) signatures and poor clinical outcome^11^. Consistently, in prostate epithelial cells, EHF knockdown enhanced stemness, tumor-initiating properties and induced EMT^11^. Reduced expression of EHF is associated with tumorigenesis in other cancers, like pancreatic^12^, kidney^13^, and colon cancer^14^. Taken together, these findings support the notion that EHF constitutes a barrier against dedifferentiation and malignant transformation in epithelial cancers.

Conversely, the TMPRSS2:ERG gene fusion, the most common genetic rearrangement in prostate tumors, leads to increased expression of the ETS factor ERG^15^. The TMPRSS2:ERG gene fusion is associated with indolent disease in cellular and mouse models, but cooperating events, like PTEN deletion or EZH2 overexpression, can unleash the transforming potential of ERG^16,17^. Intriguingly, we found that ERG overexpression and EHF loss frequently co-occur in prostate cancers^8,9,11^, raising the possibility that ectopic expression of ERG and loss of EHF could synergize and favor malignant transformation and tumor progression.

In this study, we found that reduced expression of EHF was sufficient to induce epithelial cell plasticity starting at early stages of prostate tumorigenesis. Exploring the transcriptomic landscape of prostate cancers at the bulk and single-cell levels, we found that low expression of EHF correlated in both primary and metastatic cancers with the loss of luminal epithelial identity and impaired AR signaling, both prequels to androgen indifference and anti-androgen therapy resistance. Consistently, EHF depletion in human and murine experimental models promoted a progenitor/stem cell-like state, featuring basal/luminal characteristics, high phenotypic plasticity, and the ability to transdifferentiate and acquire properties of other cell lineages. Prostate-specific EHF knockout alone (Pb-Cre4; EHF^flox/flox^) and in combination with TMPRSS2:ERG gene fusion (Pb-Cre4;EHF^flo*x/flox*^;R26*^ERG^*) in mice led to the loss of epithelial lineage integrity and enabled multi-lineage phenotypic transitions, concomitant with malignant transformation in the prostate gland. Moreover, EHF loss in both human and murine models was associated with attenuated androgenic signaling, androgen indifference, and impaired response to anti-androgen therapy, in line with our findings in human EHF^low^ primary tumors and CRPC. Mechanistically, we found that EHF directly controlled key pathways defining the epithelial cell phenotype and actively repressed multiple transcriptional regulators and downstream genes involved in cell plasticity, stemness, extracellular matrix (ECM) interactions, and the epithelial-to-mesenchymal transition (EMT). In this context, the loss of EHF in human and murine epithelial cells led to the activation of multiple downstream pathways, including the COL1A1/DDR1 and JAK/STAT3 signaling, which had a critical role in promoting non-luminal cell lineage features, attenuated androgenic response, and resistance to anti-androgen therapy. These data provide novel insights into the causes and consequences of cancer cell plasticity at the early stages of prostate tumorigenesis and a new perspective on the timing and paths to progression to androgen indifference and therapy resistance, which may prove to be highly relevant to the development of more efficient management and treatment strategies for prostate cancer patients.

## Results

### EHF maintains cell lineage integrity in human prostate epithelial cells

We have reported previously that the level of EHF is significantly reduced in a substantial portion of primary prostate tumors compared to normal healthy prostate^8,9^. Here, we examined an extensive compendium of RNA-seq datasets from multiple patient cohorts comprising normal prostate, primary tumors, CRPC, and NEPC^18^. EHF expression level was significantly reduced in CRPC and NEPC compared to normal prostate and primary tumors in line with an association of EHF loss with a more aggressive phenotype (Figure 1A, Figure S1A). Similarly, by examining single-cell (sc) RNA-seq datasets, the expression of EHF was also significantly reduced in CRPC and NE tumors (Figure 1B). Specifically, at the single-cell level, we found that epithelial cells (e.g., basal, club, luminal AR+ and AR-) in the normal prostate have high expression of EHF (Figure 1C, upper panels). In primary tumors, few cell types (e.g., epithelial luminal AR+, epithelial ERG+) retain relatively high expression of EHF, which may account for the apparent increase in primary tumors versus normal prostate in the cumulative box plots (Figure 1C, middle panels). Epithelial cells in CRPC samples exhibit drastically reduced levels of EHF, with some residual expression only in the epithelial luminal, ERG+, and basal cells (Figure 1C, lower panel and Figure S1A). Notably, the main epithelial clusters in CRPC samples were EHF-depleted and exhibited non-luminal phenotypes (e.g., epithelial proliferating AR- and neuroendocrine) (Fig. 1C, lower panels). These data supported a link between loss of EHF expression in prostate epithelial cells and lineage infidelity. Corroborating this hypothesis, gene set enrichment analysis (GSEA) comparing the transcriptome of prostate tumors with low (EHF^low^) and high (EHF^high^) EHF expression revealed activation in both primary and metastatic EHF^low^ tumors of pathways associated with loss of luminal epithelial identity, such as increased EMT and inflammatory signaling, as well as attenuated androgen response (Figure 1D). Notably, attenuated androgenic signaling and other relevant pathways were also affected in treatment-naïve EHF^low^ primary tumors, supporting a role for EHF already at early disease stages and independent of any AR-targeting therapy.

**Figure 1.**
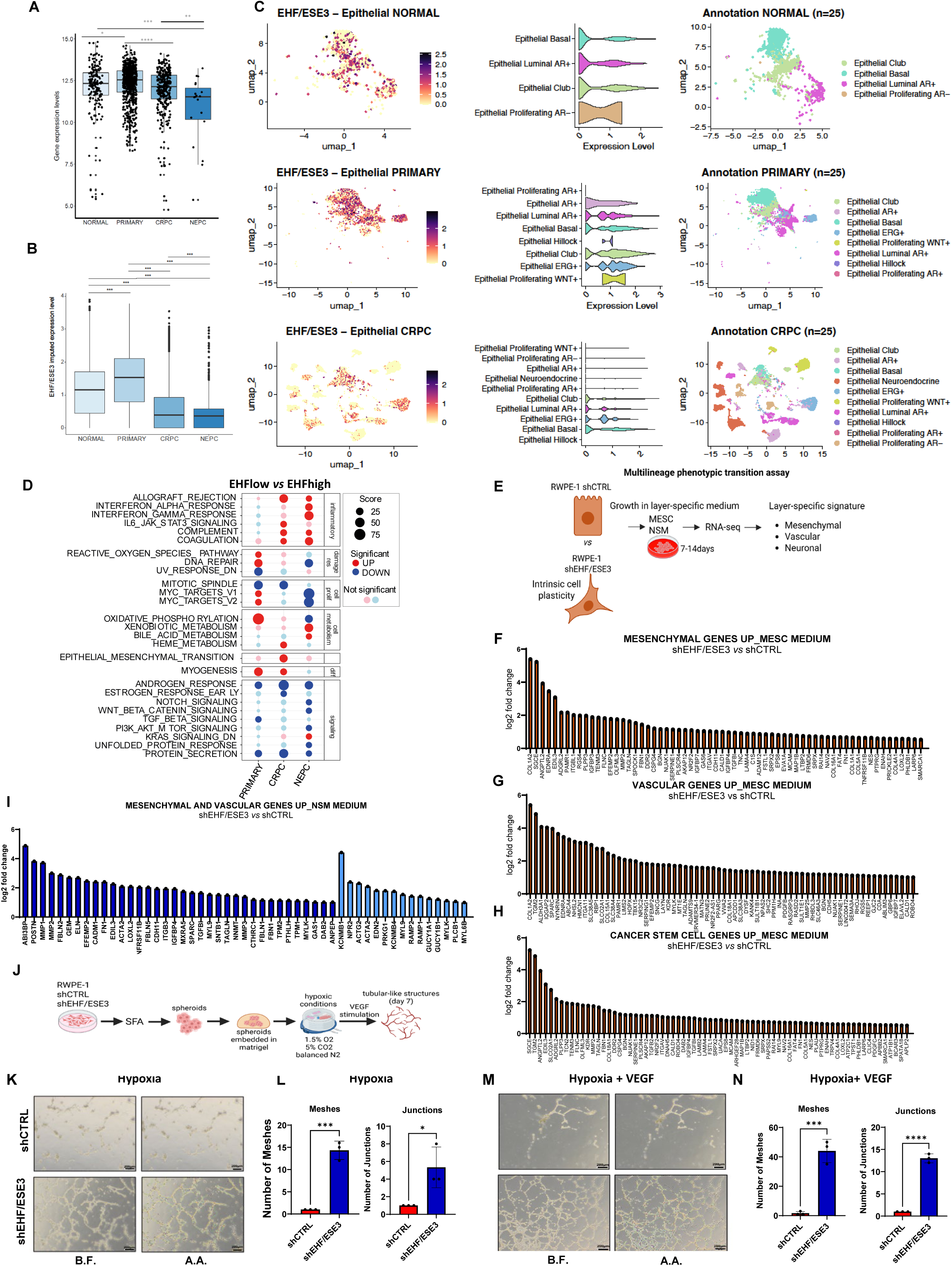
EHF loss is associated with cell plasticity in human cells and prostate tumors. **A-B** Box-plots of EHF RNA from bulk (**A**) and scRNAseq (**B**) in normal prostates and indicated prostate tumor subgroups. **C** Left panels, UMAP plots illustrate the level of EHF in scRNA-seq data from prostate samples: normal (n=25), primary tumor (n=25), and castration-resistant (CRPC) tumor (n=25). ESE3 level is indicated by a gradient scale, with high levels shown in darker colors. Central columns, violin plots of ESE3 expression levels across the indicated cell populations. Right columns, UMAP of indicated cell populations color-coded. **D** Hallmark pathways significantly enriched in EHF^low^ vs EHF^high^ gene signature extracted in primary, CRPC, and NEPC patients. **E** Workflow showing the experimental plan for alternative reprogramming of RWPE-1_ESE3Kd cells compared to RWPE-1 control upon layer-specific stimulation and RNA seq analysis. **F-H** Fold change of selected mesenchymal (**F**), vascular (**G**) and CSC-related (**H**) genes significantly upregulated in RWPE-1_ESE3Kd versus RWPE-1 shCTRL, stimulated with mesenchymal medium (MESC). **I** Fold change of selected mesenchymal and vascular gene sets was significantly up-regulated in RWPE-1_ESE3Kd versus control RWPE-1 shCTRL cells stimulated with neural medium (NSM). **J** Schematic of the experimental plan for functional vasculogenic mimicry assay in RWPE-1 shEHF/ESE3 and RWPE1 shCTRL in hypoxic and VEGF stimulation conditions. **K-L** Representative images of tubular-like structures in prostate-spheres of RWPE-1_ESE3Kd and shCTRL cells cultivated in hypoxic stimulation (**K**) and quantification of average number of meshes and junctions (**L**). Scale bar: 200µm (4X). **M-N** Representative images of tubular-like structures in prostate-spheres of RWPE-1_ESE3Kd and shCTRL cells cultivated in hypoxic + VEGF stimulation (**M**) and quantification of average number of meshes and junctions (**N**). Scale bar: 200µm (4X). Data represent the mean ±SEM. Meshes and junctions were calculated with the Angiogenesis Analyzer from ImageJ (n=3 biological replicates). B.F. stands for Bright Field; A.A. stands for Angiogenesis Analyzer. Statistical significance was determined by a two-tailed unpaired t-test. * P<0.05, *** P<0.001, ****P<0.0001. Scale bar: 200µm. See also Figure S1 and S2.

To understand the role of EHF in enforcing epithelial cell identity, we established normal human prostate epithelial RWPE-1 cells with stable EHF knockdown (RWPE-1 shEHF/ESE3) (Figure S2A). RWPE-1 cells are immortalized human prostate epithelial cells that are non-tumorigenic and express EHF. We have previously shown that EHF knockdown in two models of immortalized normal prostate epithelial cells (PrEc/LHS and RWPE1) induces malignant transformation and tumorigenic phenotypes^11^. EHF-depleted RWPE-1 cells acquired progenitor/stem cell-like and mesenchymal-like features, exhibiting increased prostate-sphere forming ability and cell migration compared to RWPE-1 shCTRL control cells (Figure S2B-C). Upon EHF depletion, RWPE-1 shEHF/ESE3 cells also formed an increased number of organoids with larger size, irregular shape, and higher Ki-67 positivity than RWPE-1 control cells (Figure S2D-E). Bulk RNA-seq showed robust differences between RWPE-1 shEHF/ESE3 and RWPE-1 shCTRL cells (Figure S2F-G-H). Notably, EMT and inflammatory response were among the top pathways enriched in the RWPE-1 shEHF/ESE3 cells (Figure S2I), indicative of enhanced phenotypic plasticity and loss of epithelial cell differentiation.

To test this aspect further, we cultured RWPE-1shEHF/ESE3 and RWPE-1 shCTRL cells in distinct lineage-specific growth conditions, i.e., in the presence of mesenchymal (MESC) and neuronal (NSM) cell culture medium and assessed the transcriptomic changes by RNA-seq (Figure 1E). Notably, RWPE-1 shEHF/ESE3 cells, compared to control RWPE-1 cells, exhibited a broad reprogramming of their transcriptome, with several genes modulated under both growth conditions (Figure S2G-H). EMT was among the pathways most affected by both stimulations (Figure S2I). Furthermore, growth in the MESC and NSM medium led to a consistent downregulation of several epithelial-specific genes, which was in line with the enhanced dedifferentiation and loss of epithelial identity (Figure S2J-L). RNA-seq revealed increased expression of mesenchymal and vasculogenesis-related genes in RWPE-1 shEHF/ESE3 incubated in MESC and NSM medium compared to RWPE-1 shCTRL cells (Figure 1F-I). Conversely, we observed the induction of neuronal genes specifically upon incubation in an NSM medium (Figure S2L). Intriguingly, growth in the MESC medium induced a robust response in terms of dedifferentiation and phenotypic plasticity with the induction of several cancer stem cell genes (Figure 1H). Thus, EHF-depleted epithelial cells exhibited a remarkable ability to adapt to different growth conditions and undertake massive transcriptomic reprogramming. Overall, these findings supported the increased phenotypic plasticity of the EHF-depleted prostate epithelial cells, which acquire the ability to dedifferentiate, bypass the lineage-specific programs, and adopt alternative cell phenotypes in response to external stimuli. Consistent with this enhanced adaptability, dissociated sphere-forming cells from RWPE-1 shEHF/ESE3 were capable of surviving and expanding in the alternative layer-specific media, while sphere-forming cells from RWPE-1 shCTRL were unable to grow (Figure S2M).

To investigate the functional relevance of this phenomenon, we tested the ability of RWPE-1 shEHF/ESE3 and RWPE-1 shCTRL cells to undergo phenotypic transition when grown in vasculogenic conditions. The epithelial-to-endothelial transition (EET) is a process by which cancer cells acquire characteristics of vascular endothelial cells and, through a process defined as vasculogenic mimicry (VM), sustain early stages of tumor vascularization^19,20^. Various factors in the tumor microenvironment favor the EET in cancer cells^19,20^. Hypoxia, which is frequent in aggressive prostate cancers, is a primary stimulus for the induction of EET and VM ^21^. To reproduce the dedifferentiation/transdifferentiation program leading to EET, we generated 3D prostate-sphere cultures of RWPE-1 shEHF/ESE3 and RWPE-1 shCTRL cells to enrich progenitor/stem-like cells. We then embedded equal numbers of prostate-sphere-forming progenitor/stem-like cells from the two cell lines in Matrigel supplemented with vasculogenic medium and incubated them in hypoxic conditions (Figure 1J). Cancer cells undergoing VM assume a vascular-like morphology with highly branched and tubular structures forming highly organized pseudo-capillary networks^19,20^. Control RWPE-1 cells had limited ability to form such vascular-like structures and tubular networks (Figure 1K). RWPE-1 shEHF/ESE3 cells, instead, expanded, adopted a vascular-like shape, and formed an extended network of tubular-like structures. A morphometric analysis of the vascular-like structures (i.e., junctions and meshes) showed a significant increase of such elements in RWPE-1 shEHF/ESE3 cells compared to control cells (Figure 1L). Adding VEGF to the medium further increased the vasculogenic phenotype of RWPE-1 shEHF/ESE3 (Figure 1M, N). Still, it did not affect control RWPE-1 cells. We observed the increased formation of vascular-like networks by prostate-spheres of RWPE-1 shEHF/ESE3 cells plated in Matrigel supplemented with VM medium, also in the absence of hypoxia (Figure S2N-Q). Furthermore, RWPE-1 shEHF/ESE3 prostate-spheres maintained in VM conditions exhibited progressively increased branching with massive outgrowth of highly irregular tubular and branched structures (Figure S2Q). In contrast, control RWPE-1 prostate-sphere cells had limited ability to expand and form branching structures in VM medium even after prolonged incubation (Figure S2Q). Together, these results indicated that EHF loss in human prostate epithelial cells induced an undifferentiated, progenitor/stem-like, high plasticity state, which favored the ability to bypass the lineage-specific programs and acquire molecular and functional features of alternative cell types.

### Prostate-specific EHF knockout in mice promotes malignant transformation

To provide in vivo evidence of the association between loss of EHF and phenotypic plasticity, we generated mice with prostate-specific conditional deletion of EHF by crossing *Pb-Cre4* and *EHF^flox/flox^* mice, in which the exon 3 is flanked by *loxP* sites for the Cre recombinase (Figure 2A and Figure S3A). The *Pb-Cre4; EHF^flox/flox^* (cEHF-KO) newborn mice were viable and showed no gross abnormalities or somatic defects (Figure S3B). The prostates of adult cEHF-KO and WT mice (age 8 to 36 weeks) were similar in size (Figure S3C). All prostate lobes at the various age groups of the cEHF-KO mice had suppressed EHF expression (Figure S3D). Histopathological alterations, like hyperplasia and mouse prostate intraepithelial neoplasia (mPIN), were visible as early as 8-12 weeks of age in homozygous cEHF-KO mice (Figure 2B-C and Figure S3E). Conversely, prostates from WT mice did not reveal any histopathological lesions (Figure 2C and Figure S3 F-G). Notably, mPIN prevailed in the younger cEHF-KO mice (8-12 weeks), and adenocarcinoma foci were more frequent in the older mice (≥13 weeks). This pattern was associated with a significant increase in the cell proliferation marker Ki67 in cEHF-KO mice compared to WT mice (Figure 2D-E). Partial loss of EHF in heterozygous cEHF-KO (cEHF-HET) mice was also associated with signs of malignant transformation. We found mainly hyperplasia at an early age (8-12 weeks) and mPIN and a few adenocarcinomas in older animals, in line with a deleterious impact of even partially attenuated EHF expression (Figure S3H-I). Together, these results demonstrate that the loss of EHF drives malignant transformation in the mouse prostate.

**Figure 2.**
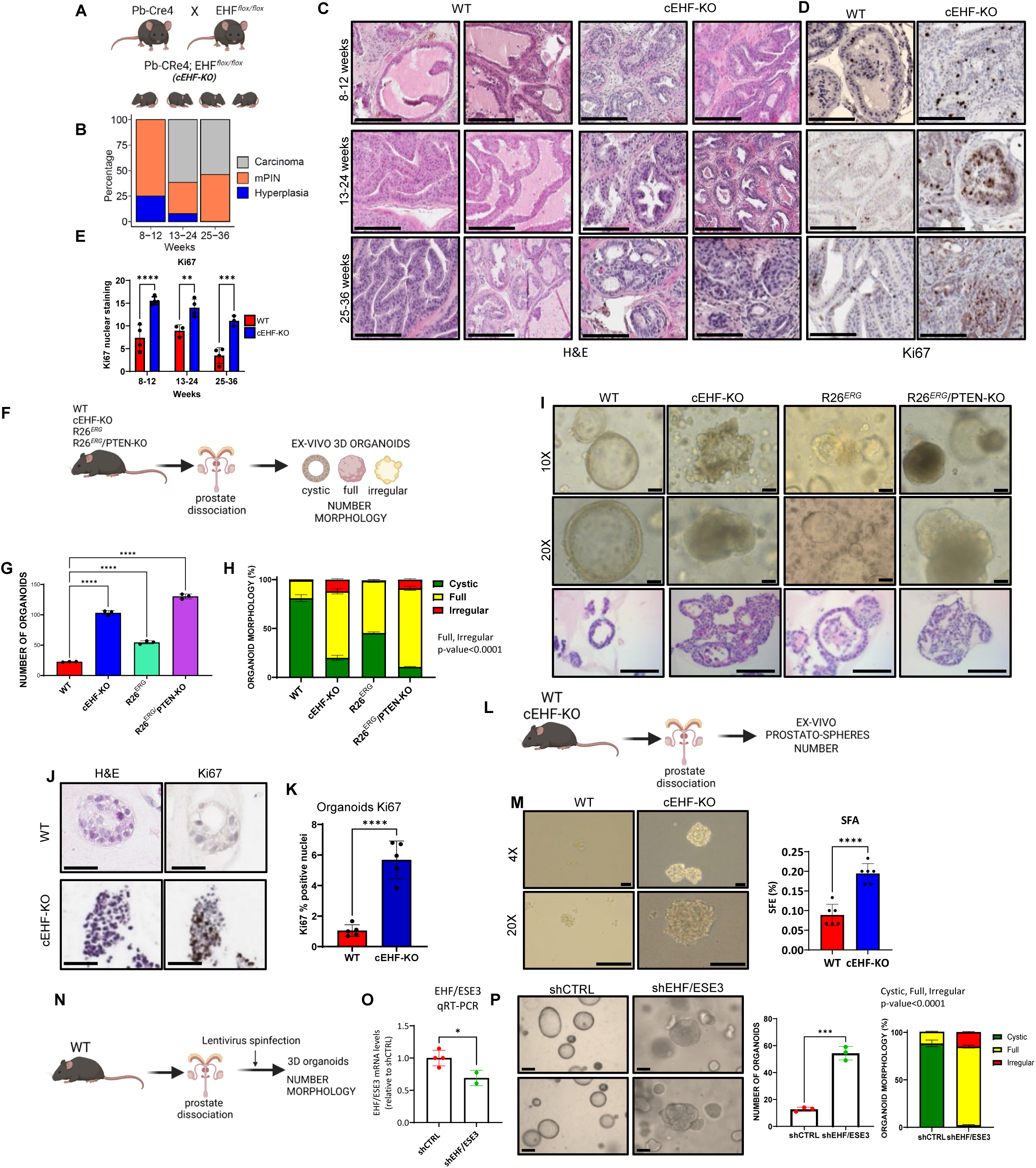
Prostate-specific EHF knock-out promotes prostate tumorigenesis. **A** Strategy for generating conditional prostate-specific cEHF-KO mice. **B** Most advanced histologic alterations in cEHF-KO mice. **C** H&E staining of prostate lobes in WT and cEHF-KO mice. 8-12 weeks (n=8 mice); 13-24 weeks (n=13 mice); 25-36 weeks (n=13 mice); Scale bar: 200µm (20X). **D** Representative images of Ki67 staining of WT and cEHF-KO mice prostate sections. Scale bar: 200µm (20X). **E** Quantitative evaluation of Ki67 staining (n=4 biological replicates). **F** Schematic of prostate dissociation for e*x vivo* evaluation of mouse-derived 3D organoids. **G-H** Number (**G**) and morphology (**H**) of organoids from indicated mouse models (n=3 biological replicates). **I** Representative brightfield images of organoids with different phenotypes from indicated mouse models. Scale bar: 100µm (10X) and 200µm (20X). Bottom, H&E staining of organoids. Scale bar: 200µm. **J-K** H&E and Ki67 immunostaining (**J**) and scoring of Ki67 positive nuclei (**K**) in organoids from WT and cEHF-KO mice (n=5 biological replicates). Scale bar: 200µm (20X). **L** Schematic of prostate dissociation for e*x vivo* evaluation of mouse-derived prostate-sphere formation. **M** Representative images (left) and percentage (right) of prostate-spheres from WT and cEHF-KO mice (n=6 biological replicates). Scale bars: 50µm (4X) and 200µm (20X). **N** Schematic of prostate dissociation for e*x vivo* evaluation of mouse-derived WT 3D organoids upon lentiviral transduction (n=4 and 2 biological replicates). **O** EHF mRNA expression levels in WT mice after lentivirus transduction evaluated by qRT-PCR. **P** Representative images (left) and number and percentage (right) of mouse-derived 3D organoids of WT mice after lentivirus transduction (n=3 biological replicates). Scale bars: 50µm (4X). Data represent the mean ±SEM. Statistical significance was determined by a two tailed unpaired t-test. *P<0.05, **P<0.01, *** P<0.001, ****P<0.0001. See also Figures S3 and S4.

To further explore the consequences of EHF deletion, we generated ex vivo 3D organoid cultures of prostate epithelial cells dissociated from WT, cEHF-HET, and cEHF-KO prostate (Figure 2F). Organoids can reflect the characteristics of the originating tissue and recapitulate essential aspects of the malignant transformation and tumor evolution ^22,23^. cEHF-KO mouse-derived cells produced more organoids than cells from WT mice (Figure 2G). We quantified the different organoid morphologies reflecting the normal-like (i.e., cystic) or more aberrant (i.e., irregular and full) structures^23^. Notably, cEHF-KO mouse-derived prostatic cells formed fewer cystic organoids and prevalently irregular and full structures with pronounced slithering and protrusions from the organoid borders (Figure 2H-I), a feature associated with dedifferentiation and increased invasive and migratory capacity of transformed cells^6^. The cEHF-KO mouse organoids also had higher Ki67 positivity compared to WT organoids, indicative of increased proliferative potential (Figure 2J-K). Cells from cEHF-HET mouse prostates also formed organoids with a higher ratio of irregular and full structures compared to cells of WT mice, confirming that even partial reduction of the EHF level is sufficient to alter the epithelial cell phenotype and cause malignant transformation (Figure S3J-M).

Interestingly, the number of organoids and the percentage of irregular and full structures were comparable between cEHF-KO and *Pb-Cre4;Pten^flox/flox^;R26^ERG^* (R26*^ERG^*/PTEN-KO) mice, a highly tumorigenic mouse model^16^ (Figure 2G-H). Conversely, prostatic cells derived from R26*^ERG^* mice, which also have TMPRSS2:ERG fusion and ERG overexpression but a more indolent phenotype^16^, formed organoids at a lower rate and with a prevalently cystic structure compared to cEHF-KO and R26*^ERG^*/PTEN-KO mice. Notably, the characteristics of the cEHF-KO mouse-derived organoids resembled those seen in patient-derived organoids from clinically advanced cancers and in more aggressive tumor models^6^.

Malignant transformation also implies the acquisition of stem-like tumor-initiating properties by the transformed epithelial cells ^24^. Hence, starting from dissociated prostate epithelial cells from WT and cEHF-KO mouse prostate, we performed *ex vivo* prostate-sphere assays to probe their ability to self-renew and expand in stem cell-selective conditions ^25^. Prostatic cells from cEHF-KO mice formed more prostate-spheres than WT mice (Figure 2L-M), supporting the acquisition of stem cell-like properties upon EHF deletion. To test this aspect further, we transduced lentiviral EHF-targeting shRNA constructs in prostatic cells from WT mice and plated them in organoid-forming conditions (Figure 2N). We found that EHF-depleted WT mouse cells formed more organoids, with a concomitant increase in full and irregular structures, as opposed to the mainly cystic organoids formed by control WT prostatic cells (Figure 2O-P). Collectively, these data showed that loss of EHF in prostate epithelial cells drives malignant transformation in mice through the induction of an undifferentiated, progenitor/stem-like, and tumorigenic state.

### EHF depletion drives lineage infidelity in murine prostate epithelial cells

To determine the molecular consequences of EHF knockout, we performed bulk RNA-seq comparing the transcriptome of WT and cEHF-KO mouse prostate (Figure 3A). We found a clear divergence between the two groups with several differentially expressed (DE) genes (Figure 3B-C). Notably, EMT and inflammatory response genes were among the top deregulated pathways in cEHF-KO mice, along with other signaling pathways, like Notch, Hedgehog, and Wnt, implicated in stemness and cell lineage determination (Figure 3D). We also found several mesenchymal- and stem cell-related genes significantly upregulated in cEHF-KO mouse prostate. Upregulation of various EMT and stem cell genes in cEHF-KO mice (e.g., FN1, FBN, PCOLCE2, COL1A1) was confirmed by qRT-PCR (Figure S4A). Thus, transcriptomic and functional data in human and murine epithelial cells converged in supporting the loss of EHF as a driver of impaired epithelial identity and induction of a progenitor/stem-like state.

**Figure 3.**
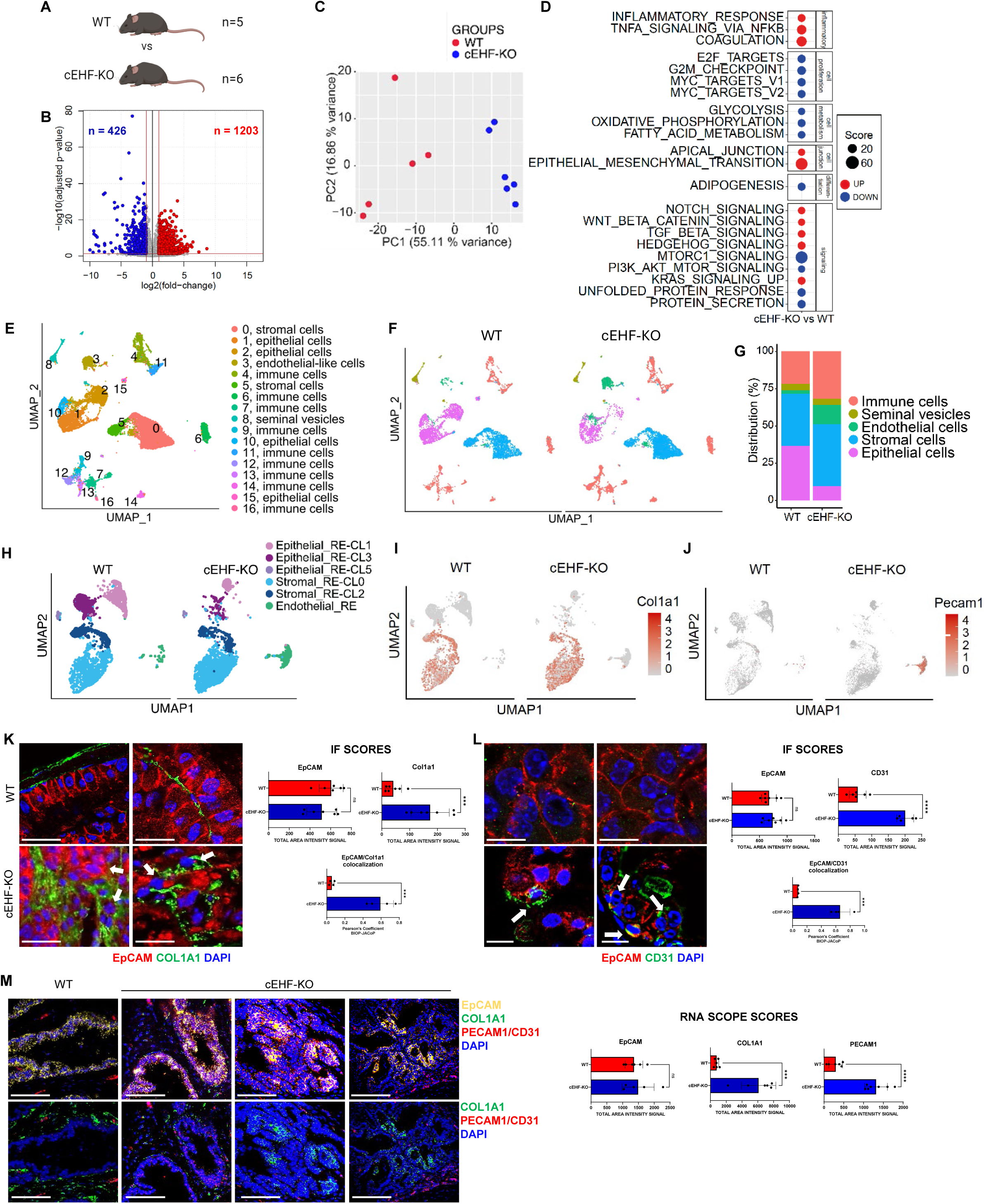
Comprehensive transcriptomic analysis reveals phenotypic plasticity in cEHF-KO mice. A Schematic of bulk RNA-Sequencing experiment. B Volcano plot showing genes significantly up- (red) and down- (blue) regulated (|log2FC| > 1) in cEHF-KO (n=5 biological replicates) vs WT (n=6 biological replicates) prostates. C PCA plot of WT (n=5 mice) and cEHF-KO (n=6 mice) samples based on bulk RNA-Seq data. D Hallmark pathways significantly enriched in cEHF-KO vs WT prostates. E UMAP analysis and cell clusters identified by unsupervised clustering by scRNA-Seq. F-G Grouping (F) and distribution (G) of main cell populations in WT (n=2 biological replicates) and cEHF-KO (n=2 biological replicates) samples. H Re-clustering of epithelial, endothelial, and stromal clusters in WT and cEHF-KO samples. I and J Distribution of Col1a1 (I) and Pecam1/CD31 (J) positive cells in the identified clusters. K-L Detection of EpCAM and Col1a1 (K) and EpCAM and CD31 (L) positive cells by immunofluorescence and confocal microscopy in WT and cEHF-KO mice (24-36 weeks). Quantitative assessment of the IF is shown in right panels. Scale bar: 30µm (K); 100µm (L top); 300µm (L bottom). M Detection of EpCAM, Col1a1 and Pecam1/CD31 positive cells by RNAScope in mouse prostate of WT (n=3 biological replicates) and cEHF-KO (n=3 biological replicates) mice. Scale bar: 300µm. Quantitative assessment of the RNAScope is shown in the right panels. See also Figure S4.

To examine further the impact of EHF knockout in the mouse prostate, we performed single-cell RNA-sequencing (scRNA-seq). Unsupervised K-nearest neighbor clustering identified 17 distinct clusters annotated through unsupervised and supervised methods, as well as expression of multiple cell type-specific markers (Figure 3E and Figure S4B). To facilitate comparisons between mouse models, we initially gathered the 17 clusters in 5 macro-clusters corresponding to the epithelial, stromal, seminal vesicle, endothelial, and immune cell populations, each characterized by cell-specific marker expression (Figure 3F and Figure S4C-D). Strikingly, the epithelial cell cluster appeared drastically reduced in cEHF-KO mice (from 36.8% to 9.8%) in favor of both the stromal and endothelial components (Figure 3G). This imbalance in cell type representation could be due to the loss of epithelial cells or, based on the bulk RNA-seq data, to the disruption of the epithelial cell identity and the consequent acquisition of alternative phenotypes by the transformed epithelial cells. Hence, this latter likely reflected a cell-intrinsic phenomenon due to the increased propensity of EHF-deleted epithelial cells to acquire progenitor/stem cell-like features and transdifferentiate, as seen with human RWPE-1 cells in vitro. In keeping with this hypothesis, the integration of bulk and scRNA-seq data showed that the most significant changes in both upregulated and downregulated genes detected in cEHF-KO mouse prostate occurred in the epithelial cell clusters, consistent with these cells being the main target of the EHF deletion (Figure S4E-F). To zoom into the relationship between epithelial, stromal, and endothelial cells, we restricted the cluster analysis to these three cell types (Figure 3H). This restricted clustering confirmed the reduction of the epithelial cell clusters with the increase of stromal- and endothelial-like cells in cEHF-KO mouse prostate. Furthermore, based on the expression of canonical luminal and basal epithelial cell type-specific markers ^26,27^, the main epithelial clusters affected by EHF deletion in terms of increased representation, differentially expressed genes, and pathway activation had both luminal-like (RE_CL3, RE_CL1) and basal-like (RE-CL5) features (Figure S4G), likely representing the collective expansion of a luminal progenitor/stem-like cell population in an undifferentiated and high plasticity state.

Consistent with the occurrence of aberrant phenotypic transitions, we found an increase in cells positive for Col1a1 (Col1a1+), a stromal cell marker, within the cEHF-KO epithelial cell clusters, marking the acquisition of mesenchymal features (Figure 3I). Conversely, Col1a1+ cells were only present in the stromal cell compartment of WT mice. We also observed an increase in cells expressing Pecam1/CD31, a typical endothelial cell marker, in the cEHF-KO mouse prostate (Figure 3J). Supporting the ability of EHF-depleted epithelial cells to transit to a hybrid, mesenchymal, and stem-like/progenitor state, confocal microscopy showed the presence of epithelial cells EpCAM^+^ concomitantly expressing Col1a1^+^ in the prostate of cEHF-KO mice (Figure 3K left and right panels). Moreover, confocal microscopy (Figure 3L left and right panels) and FACS (Figure S4H) also revealed the presence of cells co-expressing EpCAM and Pecam1/CD31 in cEHF-KO mice and absent in WT mice, consistent with cells having a hybrid epithelial-endothelial phenotype, reminiscent of the EET observed in human EHF-depleted RWPE-1 cells. Furthermore, using multiplex RNA *in situ* hybridization (RNAScope), we detected the presence of hybrid epithelial cells expressing EpCAM and COL1A1 mRNA or EpCAM and Pecam1/CD31 mRNA in the prostate of cEHF-KO mice and absent in WT mice (Figure 3M and Figure S4I). Together, these data sustained the high plasticity state achieved by murine prostate epithelial cells after deletion of EHF, with the enhanced ability to acquire aberrant non-epithelial cell lineage characteristics.

### EHF loss cooperates with ERG gene fusion by promoting phenotypic plasticity

The TMPRSS2:ERG gene fusion is the most frequent genetic rearrangement in prostate cancers ^16^. The fusion between the TMPRSS2 gene promoter and the ERG coding exons leads to abnormal expression of the ETS transcription factor ERG in prostate epithelial cells ^16^. The ERG gene fusion is associated with indolent disease in both human and mouse models. Hence, unleashing the full oncogenic potential of ERG in prostate tumors may need additional cooperating events ^8,17,25^. As we previously reported, ERG overexpression and EHF loss co-occur frequently in primary prostate cancers, raising the possibility that the depletion of EHF could synergize with ectopic expression of ERG and favor the evolution toward a more aggressive phenotype ^8,9,11^. To provide additional support to this hypothesis, analyzing bulk RNA sequencing data, we found that about 56% and 68% of ERG-positive primary tumors and CRPC/NEPC, respectively, have a mild or drastic reduction of EHF expression (Figure S5A-B),

To determine the impact of EHF loss on ERG-induced oncogenesis, we crossed the *Pb-Cre4; EHF^flox/flox^*(cEHF-KO) with *Rosa26^ERG/ERG^* (R26*^ERG^*) mice to generate the *Pb-Cre4;EHF^flo^*^x/*flox*^*;R26^ERG^*mice (hereafter cEHF-KO/R26*^ERG^*) carrying both the EHF deletion and TMPRSS2:ERG gene fusion (Figure 4A). The newborn mice were viable without gross abnormalities or defects (Figure S5C-D). The size of the prostates of the combined cEHF-KO/R26*^ERG^* mice was similar to that of cEHF-KO and WT mice (Figure S5D). Histological evaluation showed signs of transformation in all age groups examined, with a higher frequency of mPIN at an early age (8-12 weeks) and a higher number of adenocarcinomas at later times (>13 weeks), in line with an increased penetrance of the combination of ERG overexpression and EHF deletion (Figure 4B-C and Figure S5E). ERG was highly expressed, and the number of Ki67-positive cells was significantly higher in the prostate of cEHF-KO/R26*^ERG^* mice than in WT mice (Figure S5F). Consistent with previous reports^16,17^R26*^ERG^* mice exhibited a weak phenotype with no histopathological signs of malignant transformation (Figure 4C). The pathological changes in cEHF-KO/R26*^ERG^* mice with their increased frequency and intensity are, therefore, related to the combined effects of loss of EHF and gain of ERG, with EHF loss having a permissive role for ERG-induced transformation in the mouse prostate, an impact analogous to the PTEN deletion in the R26*^ERG^* mice^14^.

**Figure 4.**
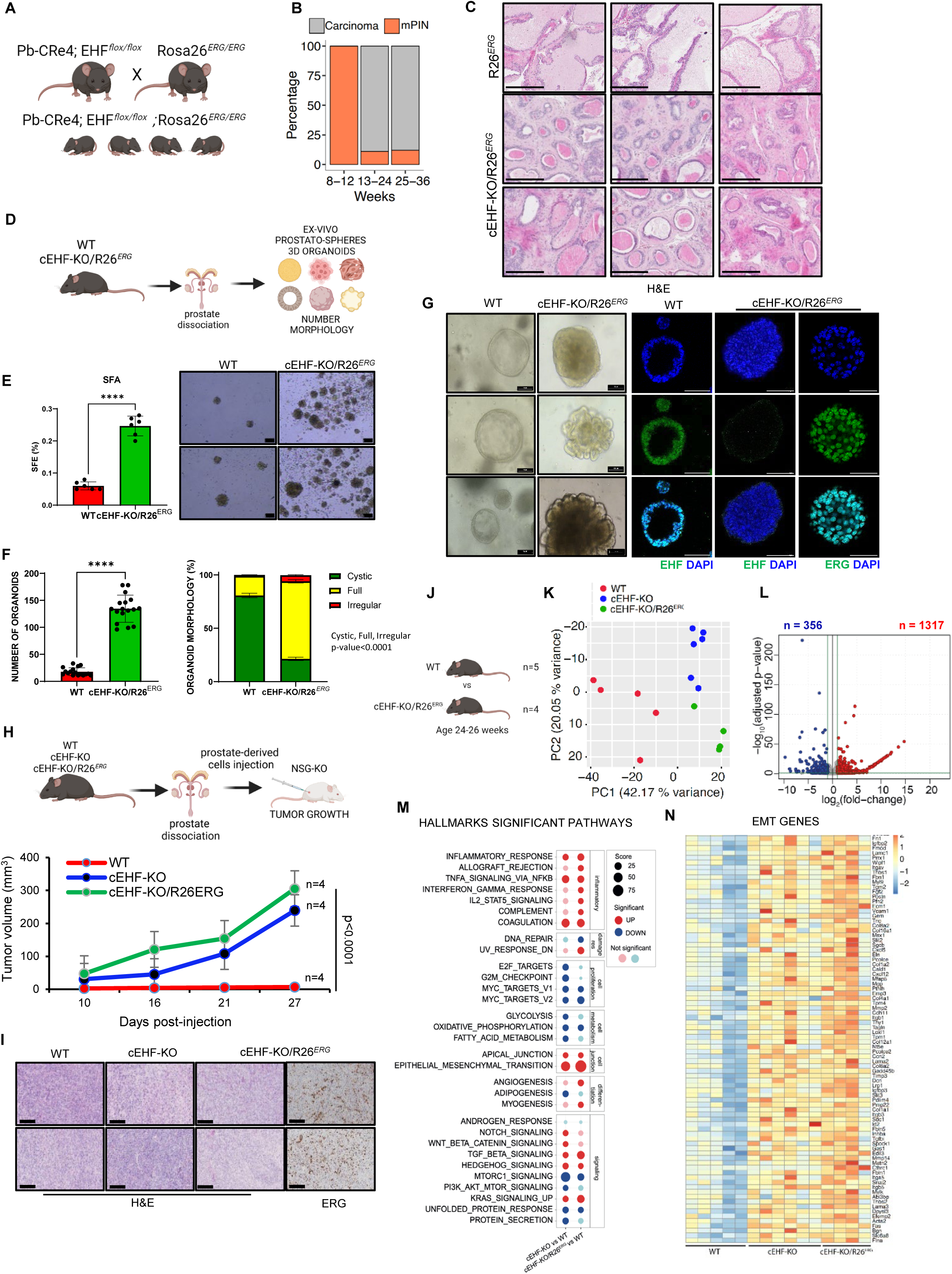
Enhanced transformation and transcriptional reprogramming in cEHF-KO/R26*^ERG^* combined mice. **A** Strategy for generating prostate-specific cEHF-KO/R26*^ERG^* transgenic mice. **B** Most advanced histologic alterations in cEHF-KO/R26*^ERG^* mice. 8-12 weeks (n=4); 13-24 weeks (n=18); 25-36 weeks (n=8). **C** H&E staining of prostate lobes in R26*^ERG^* and cEHF-KO/R26*^ERG^* mice. Scale bar: 200µm (20X). **D** Schematic of the experimental plan for ex-vivo evaluation of tumorspheres and 3D organoids formation from indicated mice models. **E** Percentage (left) and representative images (right) of prostate-spheres from WT and cEHF-KO/R26*^ERG^* mice (n=6 biological replicates). Scale bar: 50µm (4X). SFA stands for Sphere Formation Assay; SFE stands for Sphere Formation Efficiency. **F-G** Number (left) and morphological scoring (right) (**F**) of representative images and IF stain of ERG and EHF (**G**) in organoids derived from WT and cEHF-KO/R26*^ERG^* mice (n=16 biological replicates). Scale bar: 200µm left (4X), 300µm right (63X). **H** Schema (top) and subcutaneous engraftment of prostatic cells from WT, cEHF-KO and cEHF-KO/R26*^ERG^* in NSG-KO mice (n=4 mice/group), and tumor growth evaluation (bottom). **I** Representative images of tumor xenografts and ERG stain. Scale bar: 200µm (20X). **J** Schematic of bulk RNA-Sequencing experiment comparing WT (n=5), and cEHF-KO/R26*^ER^*^G^ (n=4), mice. **K** PCA plot of WT, cEHF-KO and cEHF-KO/R26*^ERG^* transcriptomic profiles. **L** Volcano plot showing genes significantly up- (red) and down-(blue) regulated (|log_2_FC| > 1) in cEHF-KO/R26*^ERG^* vs WT mouse prostates. **M** Hallmark pathways are significantly enriched in cEHF-KO/R26*^ERG^* and cEHF-KO vs WT mice. **N** EMT genes were significantly upregulated in cEHF-KO/R26*^ERG^* and cEHF-KO vs WT mice. Scale bar: 100µm. Data represent the mean ±SEM. Statistical significance was determined by a two tailed unpaired t-test, **** P<0.0001. See also Figures S5, S6, and S7.

In *ex vivo* experiments, prostatic cells from cEHF-KO/R26*^ERG^* mice formed more prostate spheres than WT mice (Figure 4D-E), indicative of higher stem cell potential. cEHF-KO/R26*^ERG^* mouse prostatic cells also generated *ex vivo* higher numbers of organoids, mostly with irregular and full structures and marked slithering, compared to WT mice (Figure 4F-G). Furthermore, when engrafted in NSG mice, both cEHF-KO and cEHF-KO/R26*^ERG^*-derived cells formed tumors, while WT-derived cells did not (Figure 4H-I). To further support the role of EHF loss in ERG-induced tumorigenesis, we transduced *ex vivo* isolated prostatic cells from R26*^ERG^* mice with a lentiviral shRNA construct targeting the murine EHF mRNA (Figure S5G). When implanted subcutaneously in NSG mice, the EHF-depleted tumor cells had a significantly higher growth rate than control R26*^ERG^* tumor cells (Figure S5H). Consistently, EHF expression and Ki67 staining increased considerably in the EHF-depleted R26*^ERG^* tumor allografts (Figure S5I-J). EHF depletion in R26*^ERG^* allografts also favored the growth of organoids *ex vivo* with aberrant morphologies compared to control cells (Figure S5K). Conversely, supplementation of EHF to WT and EHF KO murine organoids by transfecting a plasmid encoding full-length EHF resulted in significantly decreased numbers of organoids and reduced fractions of aberrant morphologies in EHF KO models, without any relevant effect in the WT counterpart (Figure S6A-E).

Bulk RNA-seq showed marked differences between cEHF-KO/R26^ERG^ and WT mice (Figure 4J-L). Transcriptomic profiling revealed similarities with cEHF-KO mice, with EMT and inflammatory signaling activated significantly in both models (Figure 4M). Several EMT and stem cell-related genes were consistently activated in both mice (Figure 4N). However, many changes in EMT gene expression and pathway enrichment were consistently more robust in cEHF-KO/R26^ERG^ than in cEHF-KO mice, indicating a significant contribution of ERG to transcriptional reprogramming in the mouse prostate. Notably, this enhanced transformation in the combined model was even more evident with scRNA-seq (Figure 5A). scRNA seq analysis revealed a substantial redistribution of the epithelial cell population in cEHF-KO/R26^ERG^ mice relative to both WT and R26*^ERG^* mice. Indeed, epithelial cells represented only 1.8% of the total cell population in cEHF-KO/R26*^ERG^* mice compared to 48.7% in R26*^ERG^* and 36.8% in WT mice. Thus, as seen in cEHF-KO mice, a substantial fraction of epithelial cells likely transited to a hybrid phenotypic state. R26*^ERG^* mice showed limited changes with a relative expansion of the epithelial clusters and restriction of the stromal-like compartment compared to WT mice. (Figure 5A, right panel). The clustering analysis restricted to epithelial, stromal, and endothelial cells confirmed the marked reduction of all the major epithelial cell clusters in cEHF-KO/R26*^ERG^* mice and suggested a massive transition of epithelial cells to a stromal-like state (Figure 5B). Interestingly, analysis of the scRNA-seq data identified a major trajectory of epithelial cell clusters transiting towards the stromal cell compartment in both cEHF-KO and cEHF-KO/R26*^ERG^* mice, sustaining the propensity of EHF-deleted prostate epithelial cells to dedifferentiate and convert to a mesenchymal lineage (Figure S7A).

**Figure 5.**
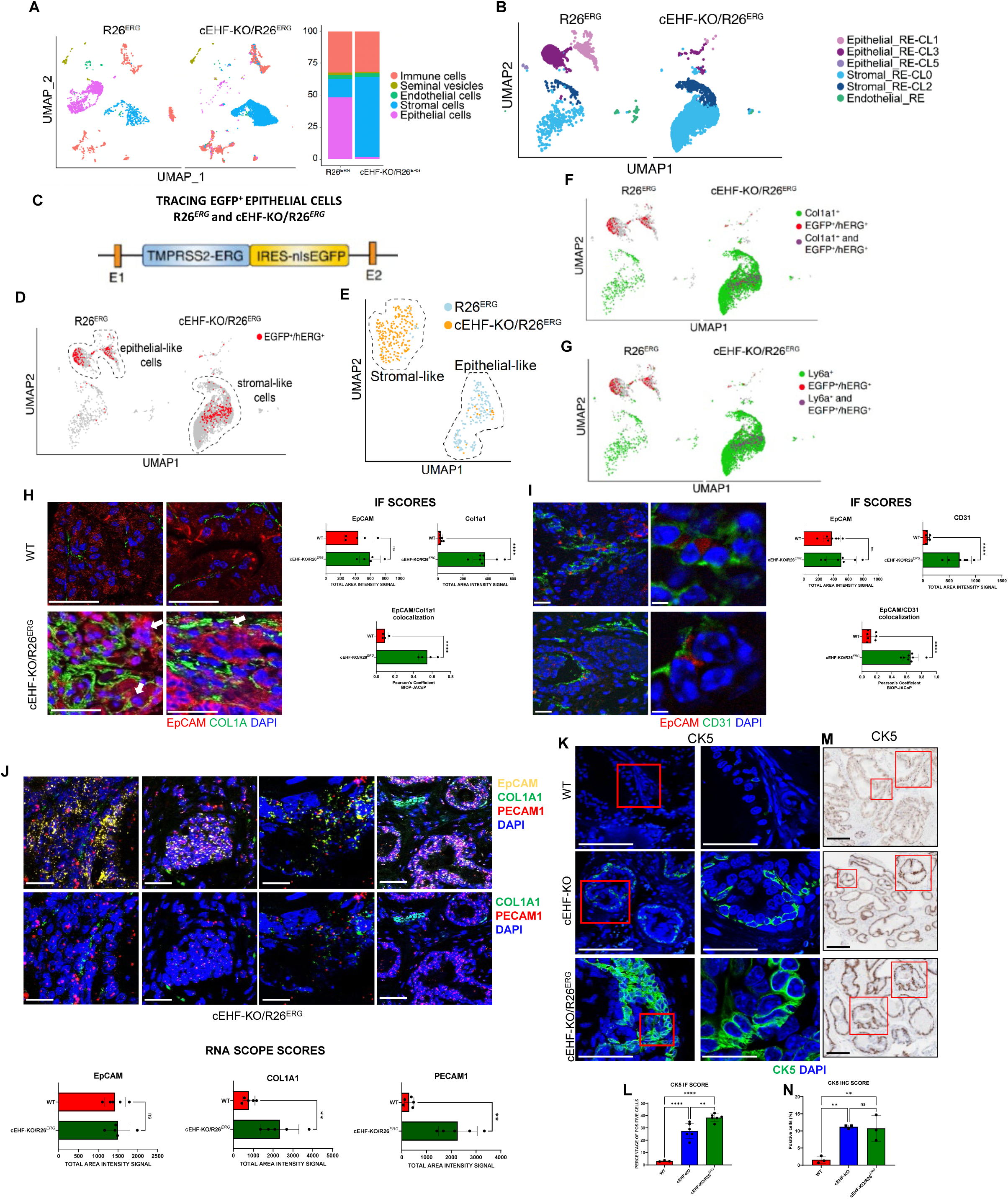
Single-cell transcriptomic and cell tracing of EGFP^+^/hERG^+^ cells provide evidence of phenotypic transition in cEHF-KO/R26*^ERG^* mice. **A** UMAP of R26*^ERG^* (n=1 biological replicates) and cEHF-KO/R26*^ERG^* (n=2 biological replicates) samples (left) and distribution of the main assigned cell clusters (right). **B** Re-clustering of the epithelial, endothelial and stromal clusters in R26*^ERG^* and cEHF-KO/R26*^ERG^* samples. **C** TMPRSS2-ERG and EGFP transgene construct in the R26*^ERG^* and cEHF-KO/R26*^ERG^* mice. **D** Distribution of EGFP^+^/hERG^+^ cells (red) in R26*^ERG^* and cEHF-KO/R2*6^ERG^* cell clusters. **E** Re-clustering of EGFP^+^/hERG^+^ cells in R26*^ERG^* and cEHF-KO/R26*^ERG^* samples. **F** Distribution of EGFP^+^/hERG^+^ (red), Col1a1^+^ (green), and double-positive (violet) cells in R26*^ERG^* and cEHF-KO/R26*^ERG^* cell clusters. **G** Distribution of EGFP^+^/hERG^+^ (red), Ly6a^+^ (green), and double-positive (violet) cells in R26*^ERG^* and cEHF-KO/R26*^ERG^* cell clusters. **H-I** Immunofluorescence detection of EpCAM and Col1a1 (**H**) and EpCAM and Pecam1/CD31 (**I**) positive cells in WT and in cEHF-KO/R26*^ERG^* mice. Scale bar: 30µm (63X); 300µm (I left) (63X); 50µm (I right) (63X). Quantitative assessment of the IF is shown in the bottom panels. **J** Detection of EpCAM, Col1a1 and Pecam1/CD31 positive cells in cEHF-KO/R26*^ERG^* prostatic tissue samples by RNAScope (n=3 biological replicates). Scale bar: 30µm. Quantitative assessment of the RNAscope is shown in bottom panels. **K-N** Representative images of immunofluorescence confocal microscopy and IHC (**K,M**) and quantification (**L,N**) of cells positive to CK5 in mouse prostates with the indicated genotypes (n=3 and 6 biological replicates/group). Scale bar: 300µm (63X left panels; 63X zoom on selected areas right panels); 200µm (M) (20X). Data represent the mean ±SEM. Statistical significance was determined by a two-tailed unpaired t-test. ** P<0.01, ****, P<0.001. n.s. not significant. See also Figure S7 and S8.

To provide direct evidence of the phenotypic transitions occurring in mouse prostate epithelial cells with EHF deletion, we examined scRNA-seq data to trace the fate of the genetically tagged epithelial cell population in cEHF-KO/R26*^ERG^* and R26*^ERG^* mice. Upon conditional Cre-mediated recombination, the prostate epithelial cells in cEHF-KO/R26*^ERG^* and R26*^ERG^* mice express the human ERG (hERG) and nuclear-targeted EGFP, which are both present in the targeting construct (Figure 5C). Hence, tracking the EGFP+/hERG+ cells should provide unambiguous information on the state of the mouse prostate epithelial cells in these mice. The EGFP^+^/hERG^+^ cells mapped exclusively within the prostate epithelial cell clusters in R26^ERG^ mice (Figure 5D). Conversely, a relevant number of EGFP^+^/hERG^+^ epithelial cells mapped within the stromal-like clusters in cEHF-KO/R26*^ERG^* mice. Thus, the genetically tagged EGFP^+^/hERG^+^ prostate epithelial cells acquired mesenchymal features and became akin to stromal-like cells in cEHF-KO/R26*^ERG^* mice. Furthermore, clustering the EGFP^+^/hERG^+^ epithelial cells in both cEHF-KO/R26*^ERG^* and R26*^ERG^* mice separated distinctly the cells retaining the epithelial phenotype from those that gained mesenchymal-like features, with the majority of EGFP^+^/hERG^+^ cells with EHF knockout mapping within the stromal-like compartment in cEHF-KO/R26*^ERG^* mice and in the epithelial clusters of R26*^ERG^*mice (Figure 5E).

In line with the gain of an undifferentiated and mesenchymal state by EHF-deleted prostate epithelial cells, we found EGFP^+^/hERG^+^ cells that were positive for the stem cell and mesenchymal markers Col1A1 (Figure 5F) and Ly6a/Sca1 (Figure 5G) in cEHF-KO/R26*^ERG^* mice, and absent in R26*^ERG^* mice. Furthermore, hybrid cells co-expressing EpCAM and Col1a1 or EpCAM and Pecam1/CD31 were detected in the prostate of cEHF-KO/R26*^ERG^* mice by multiplex RNA *in situ* hybridization (RNAScope) and immunofluorescence confocal microscopy (Figure 5H-J). Double-positive EGFP^+^ and Pecam1/CD31^+^ cells were also found in the prostate of cEHF-KO/R26*^ERG^* mice by confocal immunofluorescence microscopy (Figure S7B). Thus, in both EHF-KO mouse models, EHF deletion led prostate epithelial cells to an undifferentiated, progenitor/stem cell-like state with high cell plasticity and increased propensity to transit to other cell lineages.

In further support of this concept, we used cell surface markers (i.e., Lin, Sca-1, CD49f) to discriminate progenitor/stem-like and differentiated cells in the mouse prostate. We found a substantial increase of progenitor/stem-like cells (LSCmed, 45.7% and 35.4% vs. 3.88%) in cEHF-KO and cEHF-KO/R26ERG compared to WT mice (Figure S7C). Furthermore, LSCmed cells in cEHF-KO and cEHF-KO/R26*^ERG^* mice were mostly TROP2 positive compared to WT, in line with their progenitor/stem cell phenotype (Figure S7D). In addition to distinct cell surface markers and upregulation of various EMT and stem cell markers, prostatic progenitor/stem cells exhibit altered expression of cytokeratins, such as cytokeratin 5 (CK5) and cytokeratin 8(CK8)^28^. Indeed, we observed substantially increased staining of CK5, a basal/stem cell marker, in organoids from cEHF-KO and cEHF-KO/R26*^ERG^* mouse prostate compared to WT prostate (Figure S7E-F). Moreover, we found increased numbers of CK5-positive epithelial cells in the prostate of cEHF-KO and cEHF-KO/R26*^ERG^* mice compared to WT mice by both immunofluorescence microscopy (Figure 5K-L) and immunohistochemistry (Figure 5M-N). Concomitantly, the expression of CK8, a luminal cytokeratin, also moderately increased in the mouse organoids (Figure S7G-H) and EHF-KO mouse prostates (Figure S7I-J). Hence, along with the evidence from bulk and scRNA-seq data, the cell surface markers and cytokeratin expression pattern consistently argue in favor of the expansion of progenitor/stem-like cells with an undifferentiated or intermediate basal-luminal high plasticity state in the mouse prostate after EHF deletion.

### EHF controls the luminal cell identity and androgen responsiveness

The transcriptomic and phenotypic analyses of EHF-deleted epithelial cells indicated a switch to an undifferentiated state with the acquisition of progenitor/stem cell-like features and increased propensity to lineage trans-differentiation. To gain more insights into this phenomenon of phenotypic plasticity, we performed differential gene expression and functional annotation analysis focusing on the genetically tagged EGFP^+^/hERG^+^ epithelial cells from cEHF-KO/R26*^ERG^* and R26*^ERG^* mice. The EMT was the top-activated pathway, and many EMT and stem cell-related genes (e.g., COL1A1, Ly6a, Vimentin) were significantly upregulated in the cEHF-KO/R26*^ERG^* mice (Figure 6A and Figure S8A), corroborating the induction of a highly plastic, stem-like state in EHF-deleted epithelial cells. The activated pathways in EGFP^+^/hERG^+^ cells in the EHF-KO mice also included cytokine signaling, angiogenesis, and hypoxia, sustaining the propensity of epithelial cells to acquire mesenchymal and endothelial lineage features (Figure 6A). Furthermore, among the genes significantly upregulated (p<0.001) in EGFP^+^/hERG^+^ epithelial cells in cEHF-KO/R26*^ERG^* compared to R26*^ERG^* mice, we found many TFs implicated in stemness and cell differentiation. The top-upregulated TFs included multiple members of the AP-1, Klf, and Tcf families, as well as STAT3 and ATF3 (Figure 6B). Interestingly, the members of the AP-1 (Fos, Fosl2, Junb), Klf (Klf2, Klf4), and Tcf (Tfc4) families activated in epithelial cells from EHF-KO mice were among the key TFs discriminating AR-independent CRPC subtypes characterized by the stem cell-like (SCL) and Wnt/β-catenin (WNT) molecular signatures and driven by TCF/LEF TFs and AP-1 family TFs, respectively ^29^. Notably, bulk RNA-seq showed a similarly large network of TFs significantly activated in mouse prostates from cEHF-KO and cEHF-KO/R26*^ERG^* mice, confirming that EHF loss in epithelial cells could unleash the expression of multiple regulators of cell plasticity and stemness (Figure S8B).

**Figure 6.**
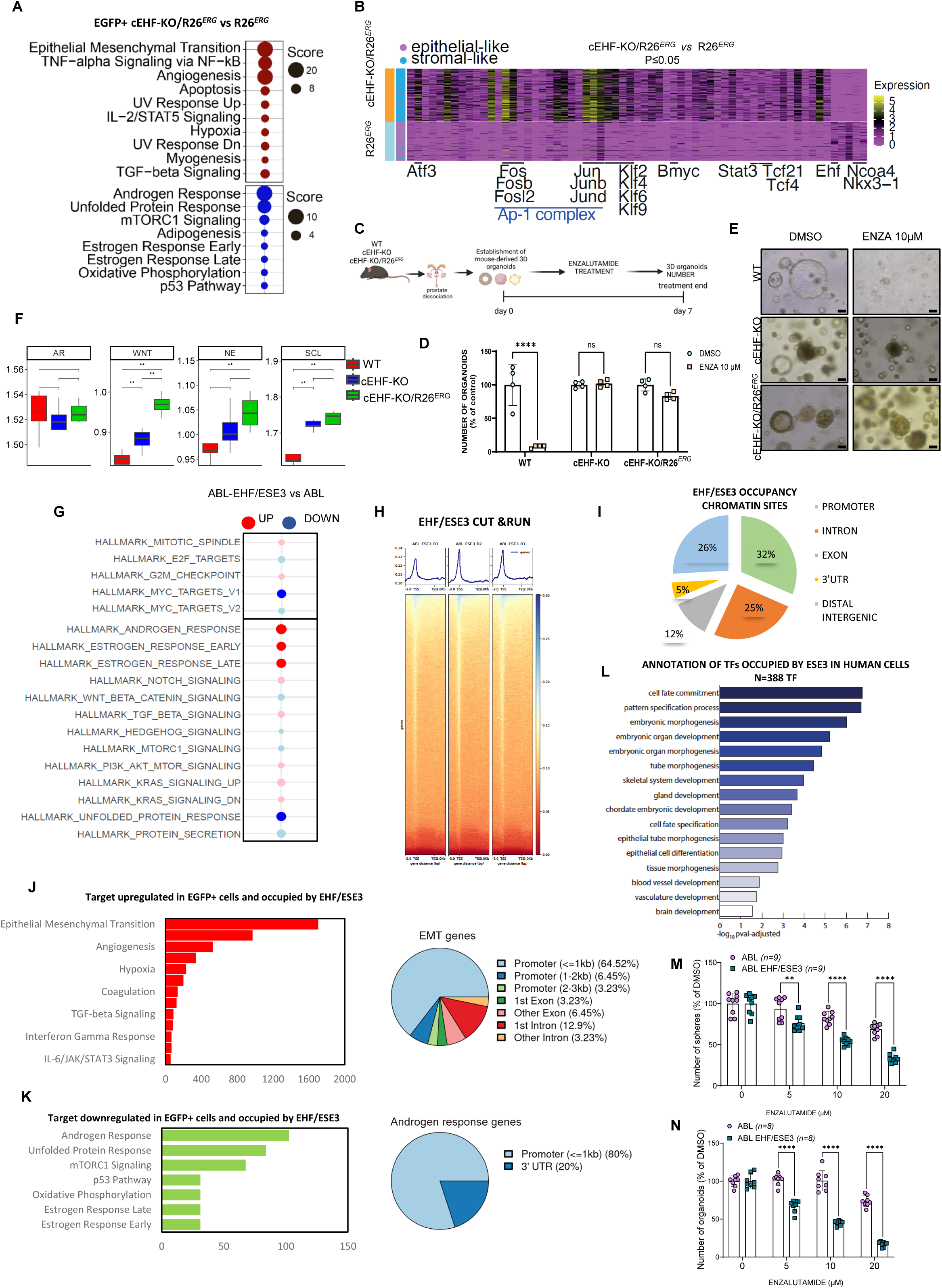
EHF orchestrates the direct control of the transcriptional landscape of prostate cell identity. **A** Hallmark pathways significantly enriched in the signature extracted from EGFP^+^/hERG^+^ cEHF-KO/R26*^ERG^* versus R26*^ERG^* prostate epithelial cells. **B** Heatmap showing TFs significantly deregulated (p<0.05) in EGFP^+^/hERG^+^ cells from the comparison of cEHF-KO/R26*^ERG^* versus R26*^ERG^*. **C** Schematic of the experimental plan for Enzalutamide treatment in GEMM-derived organoids. **D-E** Number (**D**) and representative images (**E**) of organoids derived from WT, cEHF-KO and cEHF-KO/R26*^ERG^* mice treated with the indicated concentration of Enzalutamide (n=4 biological replicates). Scale bars: 50µm (4X). **F** Application of clinical CRPC subtype-specific gene signatures in the transcriptome of WT (n=5), cEHF-KO (n=6) and cEHF-KO/R26*^ERG^* (n=4) mice. Subtype-specific signatures: AR, androgen receptor; SCL, Stem cells-like; NE, neuroendocrine; WNT, Wnt pathway. Cumulative gene expression level of AR, WNT, NE, SCL lists in WT, cEHF-KO/R26*^ERG^* and cEHF-KO samples; Statistical test: Pairwise Wilcoxon test. **G** Hallmark enrichment analysis of ABL-pESE3 versus ABL-pCTRL cells, performed with cameraPR function. Down-regulated pathways (blue), up-regulated (red). **H** Cut & Run analysis and peaks density plot of human EHF in LNCaP-ABL cells stably expressing human full-length EHF construct. **I** Peak distribution of EHF occupancy in human cells. **J** Hallmarks enriched in EHF putatively repressed targets (upregulated in EGFP+ cells and occupied by EHF). **K** Hallmarks enriched in EHF putatively induced targets (downregulated in EGFP+ cells and occupied by EHF). **L** Functional annotation of TFs occupied by EHF extracted by the Cut & Run analysis of EHF consensus peaks. **M-N** SFA (**M**) and 3D organoid assay (**N**) in indicated cell lines treated with increasing doses of Enzalutamide (n=9 and n=8 biological replicates/group). Scale bar: 100µm. Data represent the mean ±SEM. Statistical significance was determined by a two-tailed unpaired t-test, or one-way, ANOVA. ** P<0.01, **** P<0.0001. ns, no significance. See also Figure S9.

The transcriptomic analysis of the EGFP^+^/hERG^+^ cell population revealed a marked attenuation of the androgen response pathway in cEHF-KO/R26*^ERG^* mice (Figure 6A). NKX3.1, an AR target gene and transcriptional cofactor involved in canonical AR signaling, was among the significantly downregulated genes in EHF-deleted cells, even if AR was expressed (Figure 6B and S8A). Interestingly, AR immunostaining in prostatic tissue from WT, cEHF-KO, R26ERG, and cEHF-KO/R26ERG mice revealed similar levels of expression across all genotypes and age groups (Figure S8C). Similarly, we found that organoids from WT and EHF KO mice were also all positive for AR (Figure S8D). The attenuated downstream androgenic signaling associated with the expression of EMT and stem cell-related TFs and activation of alternative pathways could mark tumor cells with a high tendency to androgen indifference and resistance to anti-androgenic therapy^25^. In line with this hypothesis, we found that, despite the presence of AR, organoids from cEHF-KO and cEHF-KO/R26*^ERG^* mice were insensitive to the AR antagonist enzalutamide, which instead strongly suppressed the growth of WT organoids (Figure 6C-E). Furthermore, the addition of DHT (100 nM) stimulated the growth of WT organoids but did not affect the EHF KO organoids (Figure S8E). The association between EHF deletion and an AR-independent phenotype was further supported by an integrative bioinformatic analysis, in which we applied the transcriptional gene signatures of distinct human CRPC subtypes^29^ to the transcriptomic profiles of the murine EHF-KO prostate tumors. We found substantial activation of the SCL, WNT, and neuroendocrine (NE) signatures characterizing AR-independent CRPC subtypes in both EHF-KO models, with more marked enrichment in the combined cEHF-KO/R26^ERG^ mice (Figure 6F). These results, thus, matched the attenuated androgenic signaling seen in human EHF^low^ tumors and EGFP^+^/hERG^+^ epithelial cells in EHF-deleted mice. The AR signature, characteristic of CRPC retaining an active AR signaling (AR-CRPC), was not activated in EHF KO mice, indicating that these EHF-deleted tumors exhibited predominantly an androgen-independent phenotype with upregulation of the non-AR CRPC signatures. We also applied an AR-regulated gene signature extracted from primary tumors with low EHF expression^16^. This signature reflected a set of putatively AR-regulated genes activated in the context of low EHF expression, and therefore, associated with androgen insensitivity, high Gleason scores, and more aggressive clinical behaviors. Intriguingly, we observed significant upregulation of this non-canonical androgen-regulated gene signature activated in high-risk primary PC both in EHF-KO and cEHF-KO/R26*^ERG^* mouse tumors (Figure S8F). Thus, the loss of EHF in both murine and human prostate tumors promoted features of aggressive AR-independent CRPC subtypes characterized by activation of alternative transcriptional programs and impaired AR signaling.

To understand the mechanisms underlying the broad transcriptomic and phenotypic changes caused by EHF loss, we explored the possibility that EHF could directly supervise a network of genes influencing prostate epithelial cell fate, luminal identity, and androgenic response. To this end, we expressed the human full-length EHF in LNCaP-ABL cells that are insensitive to androgen deprivation and AR antagonists^25^ (Figure S9A). LNCaP-ABL cells derive from the parental androgen-responsive LNCaP cells, lack endogenous EHF, and thus represent a good model to assess the impact of EHF gain-of-function on the transcriptomic landscape, AR signaling, and castration resistance. RNA-seq showed robust transcriptomic changes with impact on several pathways upon EHF expression in LNCaP-ABL cells (Figure 6G). Notably, the androgen response gene set was the top-activated pathway in EHF-expressing LNCaP-ABL cells, indicating a prominent role of EHF in controlling the AR signaling and luminal identity (Figure 6G). Next, we examined EHF chromatin binding by CUT&RUN experiments (Figure 6H-I). We found a wide distribution of binding sites with peaks located in promoters, distal intergenic, and intronic regions (Figure 6I). Intriguingly, integrating the information on the EHF genomic binding sites with the genes deregulated in EGFP^+^/hERG^+^ cells in cEHF-KO/R26^ERG^ mice, we found that many activated and repressed genes (63 and 55%, respectively) were occupied by EHF, supporting a direct role of EHF in their control. Notably, the predicted EHF target genes upregulated in the mouse EHF-KO EGFP^+^/hERG^+^ epithelial cells were preferentially associated with EMT, whereas the downregulated ones were related to androgen signaling (Figure 6J-K). Overall, most of the EHF binding sites for activated and repressed genes were in promoters and proximal regulatory regions (Figure S9B), particularly in the case of EMT and AR responsive genes, which were among the most activated and repressed targets, respectively (Figure 6J-K). These findings were in line with previous studies showing that EHF can act both as a transcriptional repressor (e.g., IL-6, STAT3, EZH2) and activator (e.g., NKX3.1) of distinct sets of target genes^16^. Further integrative analysis of EHF occupancy and bulk RNA-seq in the mouse prostates of cEHF-KO mice revealed a similar and robust convergence, with 56% and 51% of activated and repressed genes, respectively, being predicted EHF targets. Notably, the putatively EHF-bound and repressed genes in the EHF-KO mice consistently included a large number of EMT and stem cell genes (Figure S9C).

Relevantly, we also found that EHF occupied regulatory regions of an extensive network of TFs (n=388). Functional annotation of this EHF TFs network revealed a significant association with cell fate commitment, pattern specification, morphogenesis, and development, sustaining the relevance of EHF as a cell lineage-determining factor (Figure 6L). In line with the transcriptomic data, EHF re-expression in LNCaP-ABL cells significantly reduced the ability to form tumorspheres and organoids (Figure S9D), indicative of reduced stemness, self-renewal, and tumor-initiating potential. EHF expression also reduced the ability of LNCaP-ABL cells to grow as tumor xenografts after subcutaneous implantation (Figure S9E-F). Moreover, in line with the impact of EHF on luminal cell identity and androgenic signaling, EHF-expressing LNCaP-ABL cells were more responsive to enzalutamide than parental LNCaP-ABL cells in tumorsphere and organoid assays (Figure 6M,N) Overall, these data provided a mechanistic explanation for the relevant role of EHF in orchestrating the transcriptional landscape and luminal identity of prostate epithelial cells, with its loss driving aberrant phenotypic features, androgen indifference and resistance to anti-androgen therapy.

### Cell plasticity and tumor progression in EHF-depleted prostate cancers

We reasoned that examining the transcriptional profiles of the individual cell clusters identified by scRNA-seq in cEHF-KO, cEHF-KO/R26*^ERG,^* and WT mice could help to determine the pathways specifically affected by EHF depletion and relevant for epithelial phenotypic plasticity (Figure 7A). The epithelial clusters (e.g., RE-CL3) exhibited the most evident changes in terms of the number of deregulated genes and activated pathways in both cEHF-KO and cEHF-KO/R26*^ERG^* mice. Many of these pathways were significantly downregulated in the stromal cell clusters, confirming the relevance of the specific changes in the epithelial cell population. We found significant activation of inflammatory signaling (e.g., IL-6/JAK/STAT3), EMT, and other oncogenic pathways (e.g., hypoxia, TGF-β, KRAS) in the epithelial clusters of cEHF-KO and cEHF-KO/R26*^ERG^* mice compared to WT and R26*^ERG^* mice (Figure 7A, B). Furthermore, the pathways activated in the mouse EHF-KO epithelial clusters (e.g., IL6-JAK-STAT3, EMT) substantially reflected those enriched in human EHF^low^ prostate tumors, particularly CRPC and NEPC, sustaining the relevance of EHF loss in the most advanced and aggressive tumors. Consistently, using epithelial-specific transcriptional signatures of EHF-KO mouse models (i.e., genes activated in the RE-CL3 epithelial cluster), we found a significant signature enrichment in both human primary and castration-resistant EHF^low^ tumors compared to EHF^high^ tumors (Figure 7C).

**Figure 7.**
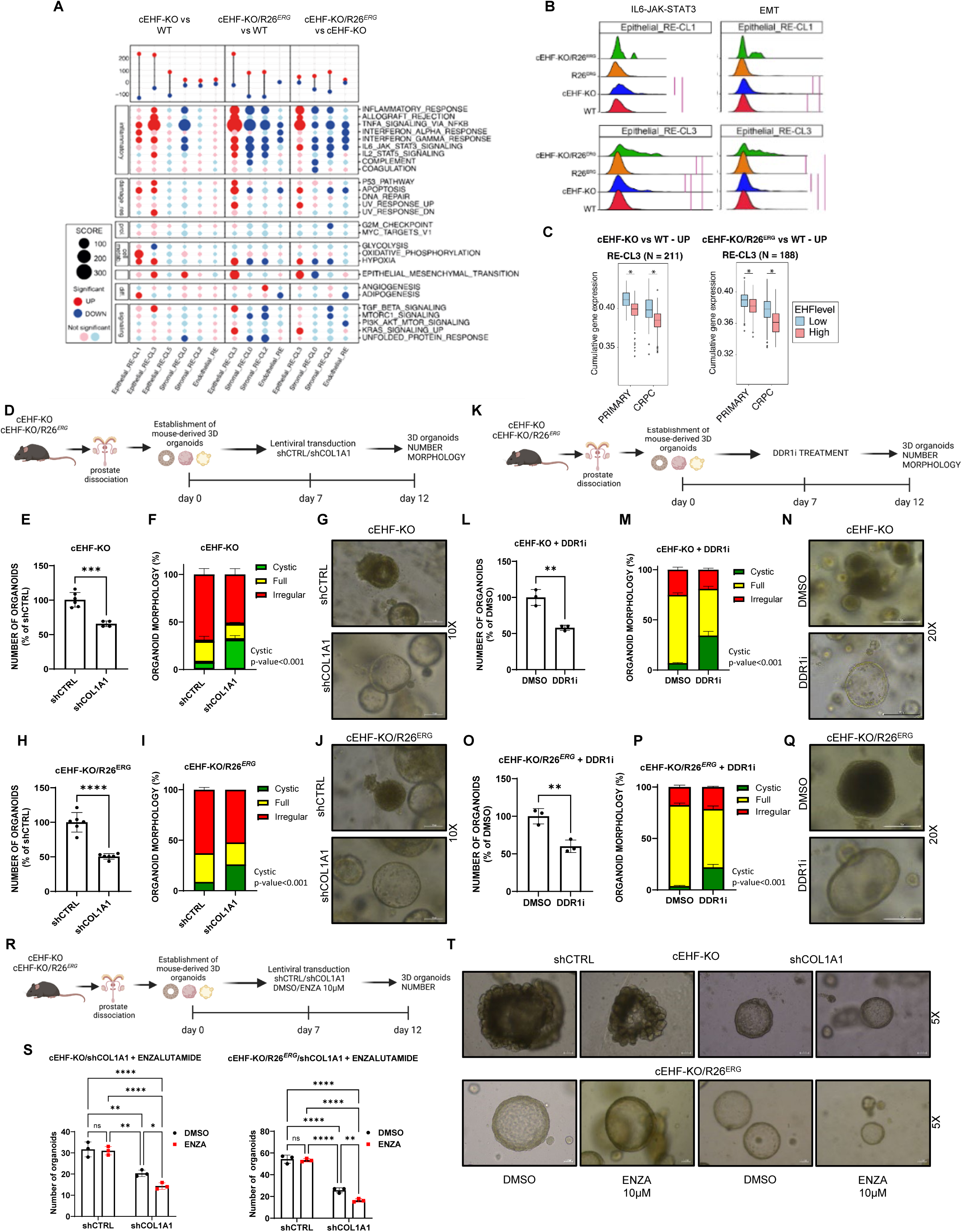
COL1A1 sustains loss of cell identity and androgen-indifferent status in cEHF-KO mice. **A** Top panel, number of significantly up-regulated (in red) and down-regulated (in blue) genes and (bottom panel) hallmark pathways significantly enhanced (in red) and repressed (in blue) for the indicated comparisons. **B** Ridge plots of the distribution of cumulative score computed for each cell for IL6/Jak/Stat3 and EMT gene sets in indicated mice. **C** Cumulative gene set activated in the epithelial cluster RE-CL3 in EHFlow primary and CRPC human prostate tumors **D** Schematic of the experimental plan for e*x vivo* establishment and evaluation of mouse-derived 3D organoids upon lentiviral transduction against COL1A1. **E-G** Number (**E**) morphology scoring (**F**) and representative images (**G**) of organoids from cEHF-KO in control (shCTRL) and COL1A1 ablated samples (shCOL1A1) (n=6 biological replicates). Scale bars: 100µm (10X). **H-J** Number (**H**) morphology scoring (**I**) and representative images (**J**) of organoids from cEHF-KO/R26*^ERG^* mice in control (shCTRL) and COL1A1 ablated samples (shCOL1A1) (n=6 biological replicates). Scale bars: 100µm (10X). **K** Schematic of the experimental plan for e*x vivo* establishment and evaluation of mouse-derived 3D organoids following treatment with the DDR1 inhibitor (DDR1i). **L-N** Number (**L**) morphology scoring (**M**) and representative images (**N**) of organoids from cEHF-KO following treatment with the DDR1i (n=3 biological replicates). Scale bars: 100µm (20X). **O-Q** Number (**O**) morphology scoring (**P**) and representative images (**Q**) of organoids from cEHF-KO/R26*^ERG^* mice following treatment with the DDR1i (n=3 biological replicates). Scale bars: 100µm (20X). **R** Schematic of e*x vivo* establishment and evaluation of mouse-derived 3D organoids upon lentiviral transduction against COL1A1 and Enzalutamide treatment. **S-T** Number of organoids from cEHF-KO and cEHF-KO/R26*^ERG^* (**S**) and representative images (**T)** following lentiviral transduction and treatment with Enzalutamide (ENZA) (n=3 biological replicates). Scale bars: 100µm (5X). Data represent the mean ±SEM. Statistical significance was determined by a two-tailed unpaired t-test, or one-way, ANOVA. * P<0.05, ** P<0.01, *** P<0.001, **** P<0.0001. ns, no significance. See also Figure S10.

Together, these findings suggested that EMT and inflammatory signaling could be the central drivers of the phenotypic alterations emerging in EHF-depleted prostate cancer cells. To test this hypothesis, we focused on collagen-encoding genes. We found that many collagen genes, including COL1A1, had evidence of EHF genomic binding and could be putatively regulated by EHF (Figure S10A). COL1A1 mediates relevant oncogenic signals, underlying ECM interactions, cell-cell communications, tumor-initiating and metastatic properties^30,31^. Notably, COL1A1 signaling is associated with stemness and pluripotency in both mammary and prostate glandular epithelial cells^32^. COL1A1 was activated in all the EHF-depleted human and mouse models (i.e., EHF KO murine cells and RWPE-1_EHFKd cells) and repressed in EHF-expressing LNCaP-ABL, arguing in favor of a direct control by EHF. This tight association was also consistent with the presence of COL1A1+ epithelial cells, which represented a subpopulation of epithelial cells that had undergone a phenotypic transition and had acquired stem-like properties upon EHF deletion. Hence, we hypothesized a direct link between COL1A1 and phenotypic plasticity with the induction of a progenitor/stem cell-like and an androgen-indifferent state due to COL1A1 expression in EHF-depleted epithelial cells. To test this, we knocked down COL1A1 by lentiviral transduction with a murine-specific shRNA in cEHF-KO and cEHF-KO/R26*^ERG^* mouse organoids (Figure 7D and Figure S10B). COL1A1 knockdown led to a significant reduction in the number of organoids in both models. We also observed a shift toward a cystic morphology and luminal-like phenotype in organoids from both cEHF-KO (Figure 7E-G) and cEHF-KO/R26*^ERG^* (Figure 7H-J) mice, in line with a relevant role of COL1A1 in promoting the aberrant phenotypes of EHF-depleted epithelial cells.

Secreted COL1A1 can bind to the cell surface receptor DDR1 and act in an autocrine/paracrine manner, mediating the ligand-dependent activation of intracellular downstream effectors^31^. Hence, we tested whether COL1A1-DDR1 signaling had a role in the context of EHF loss, evaluating the impact of a DDR1 inhibitor on the growth of EHF-KO tumor organoids (Figure 7K). Notably, inhibiting DDR1 reduced organoid growth and reversed their aberrant morphology, increasing the fraction of cystic organoids in both cEHF-KO (Figure 7L-N) and cEHF-KO/R26*^ERG^* (Figure 7O-Q) mice. Together, the genetic and pharmacological interventions sustained the relevance of COL1A1 in the epithelial cell phenotypic plasticity in the EHF-deleted context. Moreover, consistently with the shift toward a more luminal phenotype, ablation of COL1A1 was associated with increased response to the AR antagonist enzalutamide in both murine models (Figure 7R-T). Notably, co-treatment with a low dose of DDR1i (2.5 µM) also rendered the EHF-KO tumor organoids more sensitive to enzalutamide (Figure S10C-D). Together, these data support a previously unrecognized link between COL1A1-induced phenotypic plasticity and the acquisition of an androgen-indifferent castration-resistant state and suggest novel therapeutic strategies in the context of EHF loss.

Cell-autonomous activation of inflammatory signaling pathways (e.g., IL-6/JAK/STAT3) was another acquired characteristic of EHF-deleted prostate epithelial cells (Figure 7A-B). Indeed, in line with the transcriptomic data in EHF-deleted epithelial cells, we found significant activation of the IL-6/JAK/STAT3 pathway with increased phosphorylation of STAT3 at tyrosine 705 (pSTAT3) in the prostate of cEHF-KO and cEHF-KO/R26*^ERG^* mice compared to WT mice (Figure 8A-B). We had previously reported various EHF-dependent mechanisms leading to IL-6/JAK/STAT3 activation in prostate cancers^33,34^. We hypothesized that COL1A1/DDR1 signaling could also contribute to the altered IL6-JAK-STAT3 signaling in EHF-deleted prostate epithelial cells by enhancing JAK/STAT signaling and STAT3 phosphorylation^31^. In line with this hypothesis, COL1A1 depletion resulted in reduced pSTAT3 in cEHF-KO and cEHF-KO/R26*^ERG^* mouse organoids (Figure 8C). Inhibiting DDR1 had the same effect on pSTAT3 (Figure 8D), indicating a direct impact of the COL1A1/DDR1 axis on STAT3 activation in these EHF-KO models. On the other hand, the JAK/STAT3 inhibitor ruxolitinib, which inhibited pSTAT3, reduced tumor organoid growth and significantly increased the fraction of cystic, luminal-like organoids at the expense of the irregular and full structures (Figure 8E-I), indicating JAK/STAT3 as a critical pathway downstream of the EHF and COL1A1/DDR1 axis. Intriguingly, JAK/STAT3 inhibition also affected COL1A1 levels in EHF-KO-derived tumor organoids (Figure 8F), hinting at a feed-forward loop reciprocally controlling these key pathways. We observed similar results in human RWPE-1 cells with EHF knockdown. RWPE-1 shEHF/ESE3 cells, particularly the prostate sphere-forming cell fraction, had high expression of both COL1A1 and pSTAT3 compared to RWPE-1 shCTRL control cells (Figure 8J-K). COL1A1 knockdown in RWPE-1_EHF-Kd cells inhibited prostate-sphere growth and migration (Figure 8L-N). Similarly, ruxolitinib reduced the growth of tumor-spheres of RWPE-1 shEHF/ESE3 cells (Figure S11A-B). Furthermore, ruxolitinib reduced both pSTAT3 and COL1A1 levels in RWPE-1 shEHF/ESE3 cells, sustaining the link between the STAT3 activation and COL1A1 expression (Figure S11C). These data support a signaling axis between EHF, COL1A1/DDR1, and JAK/STAT3 that, upon EHF loss, disrupts luminal cell identity and drives stemness and phenotypic plasticity in prostate cancer. Moreover, inhibiting COL1A1/DDR1 and JAK/STAT3 signaling reverted the phenotypic plasticity in EHF-depleted models, sustaining the relevance and therapeutic potential of this axis in prostate cancer.

**Figure 8.**
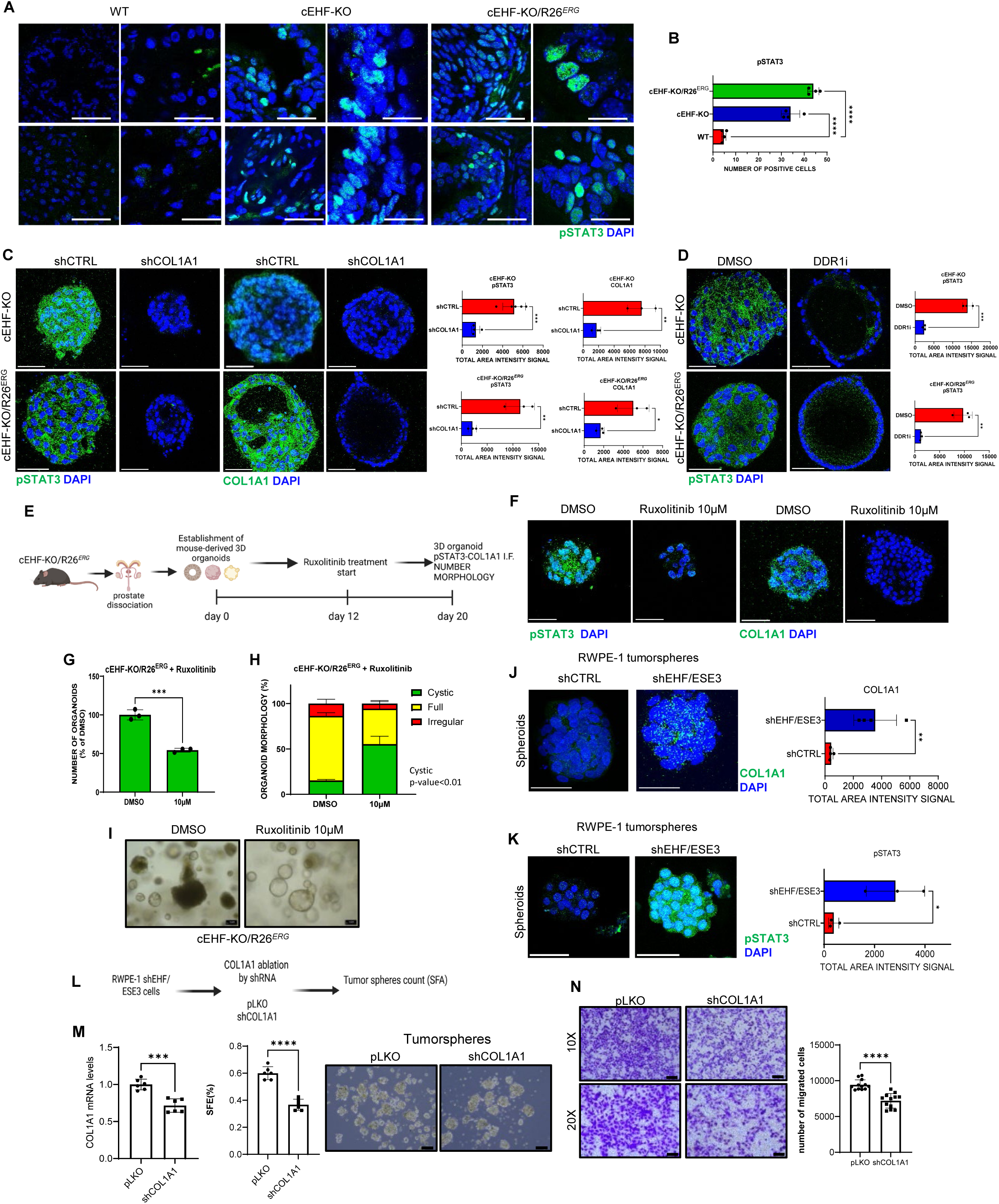
Crosstalk between JAK-STAT3 and COL1A1 sustains cell plasticity in EHF depleted context. **A-B** Immunofluorescence staining (**A)** and quantitative assessment (**B**) of phosphorylated STAT3 in mouse prostates with indicated genotypes (n=4 biological replicates). Scale bars: 100µm (63X left panels; 63X zoom on selected areas right panels). **C-D** Detection by immunofluorescence and confocal microscopy of phosphorylated STAT3 and COL1A1 in cEHF-KO and cEHF-KO/R26^ERG^ mouse organoids transduced with lentiviral targeting COL1A1 (**C**) or treated with DDR1i or DMSO (**D**) (n=5 and n=3 biological replicates). Scale bar: 100µm (63X). Quantitative assessment of the IF is shown in C-D. **E** Experimental outline for testing phenotypic reversal by JAK/STAT inhibition using Ruxolitinib in mouse organoids. **F-I** Detection by immunofluorescence and confocal microscopy of phosphorylated STAT3 (left) and COL1A1 (right) in cEHF-KO/R26*^ERG^* mouse organoids treated with Ruxolitinib or DMSO (**F**). Scale bar: 100µm (63X). Number **(G)** and morphological scoring (**H**) and representative images (**I**) of cEHF-KO/R26*^ERG^* mouse organoids treated with Ruxolitinib or DMSO (n=3 biological replicates). **J** Representative images (left) of immunofluorescence confocal microscopy of tumorspheres positive to pSTAT3 and quantification (right) in RWPE-1 shEHF/ESE3 cells. Scale bar: 100µm (63X). **K** Representative images (left) of immunofluorescence confocal microscopy of tumorspheres positive to COL1A1 and quantification (right) in RWPE-1 shEHF/ESE3 cells. Scale bar: 100µm (63X). **L** Schematic of lentiviral transduction against COL1A1 for SFA and migration assay. **M** COL1A1 mRNA levels evaluated by qRT-PCR (left) in RWPE-1 shEHF/ESE3 cells after shCOL1A infection (n=6 biological replicates). Prostate-spheres counting (middle) and representative images (right) of RWPE-1 shEHF/ESE3 after COL1A1 ablation. Scale bar: 50µm (10X) (n=6 biological replicates). **N** Representative images (left) and quantification (right) of migrating RWPE-1 shEHF/ESE3 upon shCOL1A infection in Boyden chamber assay at 24 hours. Scale bar: 100µm (10X), 100µm (20X) (n=12 biological replicates/group). Data represent the mean ±SEM. Statistical significance was determined by a two-tailed unpaired t-test, or one-way, ANOVA. * P<0.05, ** P<0.01, *** P<0.001, **** P<0.0001. See also Figure S10 and Figure S11.

## Discussion

Phenotypic plasticity is a hallmark of cancer and has relevant implications for tumor progression and treatment resistance^2,35,36^. In this study, we identify a novel regulatory axis centered on the ETS factor EHF and describe the cascade of events unleashed by the disruption of this essential transcriptional regulator of epithelial cell lineage identity. EHF acts as an epithelial-specific master regulator that defines the fate of prostate epithelial cells and restricts malignant transformation, cell plasticity, and phenotypic transitions to other cell lineages. We show that both in human and murine prostate epithelial cells, the loss of EHF is sufficient to induce a progenitor/stem cell-like and highly plastic state, disrupting the cell lineage integrity and leading to malignant transformation. These findings are clinically relevant as they identify EHF loss as a critical event that unlocks cancer cell plasticity early in prostate cancer evolution and increases the tendency toward multi-lineage phenotypic transitions and acquisition of androgen-independent aggressive phenotypes. Moreover, reduced level of EHF and attenuated androgen signaling were evident in treatment-naïve primary tumors in addition to CRPC, supporting a role of EHF already at early disease stages and independent of AR-targeting therapies. Indeed, our data from human samples and murine models indicate that EHF loss bypasses androgen stimulation and canonical AR signaling, even in the presence of AR, creating a state of androgen indifference or independence and promoting resistance to AR-targeted therapies.

Using bulk RNA- and scRNA-seq data, we found that a significant fraction of prostate tumors, particularly aggressive CRPC subtypes, had low expression of EHF. At the single-cell level, epithelial cells in normal prostate and many primary tumors express EHF. In contrast, most epithelial cells in CRPC samples had drastically reduced expression of EHF and a non-luminal phenotype, underscoring the link between EHF, luminal identity, and tumor progression. Thus, EHF appears as a barrier to malignant transformation and aberrant transdifferentiation of prostate epithelial cells. Indeed, low EHF expression marked primary and advanced tumors with high epithelial-mesenchymal plasticity, reduced androgenic response, and increased tendency to androgen independence and castration resistance. These findings are highly relevant, providing a potentially discriminating factor for identifying tumors with a higher risk of AR-independent evolution while suggesting new insights for possible therapeutic strategies.

Unlocking cell plasticity due to the reduced expression of EHF is an epithelial cell-intrinsic process, which we observed by multiple approaches in cultured cells, tumor allografts, and transgenic mouse models. Strikingly, human prostate epithelial cells with EHF knockdown exhibited a progenitor/stem cell-like phenotype, enhanced cell plasticity, and ability to undergo multi-lineage transitions. Indeed, EHF-depleted epithelial cells underwent a broad reprogramming of their transcriptome when exposed to different environmental stimuli. Under conditions set to promote mesenchymal and neuronal differentiation, EHF-depleted epithelial cells upregulated lineage-specific markers and reduced expression of typical epithelial markers. These transcriptional changes mirrored the acquisition of functional phenotypes with progenitor/stem-like and mesenchymal capabilities (e.g., enhanced tumor sphere formation and migration). Intriguingly, when grown under hypoxia and vasculogenic conditions, EHF-depleted prostate epithelial cells with stem-like properties could undergo EET and vascular mimicry with the formation of extensive tubular vascular-like networks, a critical step favoring early tumor vascularization. Thus, EHF-depleted epithelial cells, through these multi-lineage transitions, acquire multiple capabilities, including self-renewal, migratory, invasive, and vasculogenic properties, essential for tumor development and progression.

RNA-seq data in the EHF-KO mice corroborated in vivo the enhanced plasticity and phenotypic transitions observed in human epithelial cells. Intriguingly, EHF knockout in mouse prostates promoted phenotypic changes, generally observed in late-stage tumors with multiple genetic hits, like p53 and pRb mutations^3^. Furthermore, ERG knock-in combined with EHF deletion accelerated the onset of malignant lesions in the context of enhanced phenotypic plasticity caused by the inactivation of EHF. Thus, loss of EHF had a permissive role for ERG-induced malignant transformation in the prostate. Specifically, the loss of luminal identity and multi-lineage transition capability in the single and combined models depended strictly on the EHF deletion and did not occur in the single R26*^ERG^* transgenic mice. Direct tracing of the genetically tagged EGFP+/hERG+ cells in cEHF-KO/R26*^ERG^* mice using scRNA-seq data provided an unambiguous demonstration of the epithelial cell phenotypic plasticity and transitions to a mesenchymal-like state. Indeed, most EGFP^+^/hERG^+^ epithelial cells in the EHF KO mice transited within the stromal-like clusters, whereas they mapped exclusively with the epithelial clusters in R26*^ERG^* mice. Numerous EGFP^+^/hERG^+^ epithelial cells in cEHF-KO/R26*^ERG^* mice were positive for the mesenchymal and stem cell markers (e.g., Col1A1 and Ly6a/Sca1). Furthermore, we identified hybrid cells co-expressing mesenchymal or endothelial markers (e.g., EpCAM and Col1a1 or EpCAM and Pecam1) in the prostate of EHF KO mice using orthogonal approaches. In line with the gain of an undifferentiated and mesenchymal state, we also observed a substantial increase of prostate epithelial cells with progenitor/stem cell-like phenotype in EHF-KO mouse prostates based on cell surface markers and cytokeratin expression.

Phenotypic plasticity induced by EHF ablation is mainly a cell-autonomous phenomenon driven by the activation of cell-intrinsic pathways, although microenvironmental and cell-cell interactions might also contribute. The transcriptomic analysis of the epithelial cell clusters in EHF KO mice provided in-depth information on the pathways activated in the plastic epithelial cells. Notably, many deregulated pathways in the prostate epithelial cells in EHF KO mice (e.g., EMT, inflammatory signaling, and JAK/STAT3) were similar to the pathways enriched in human primary and castration-resistant EHF^low^ tumors, sustaining the relevance of these findings to the clinical setting. Furthermore, the integration of the genomic EHF binding sites and transcriptomic data showed that many genes activated in EHF-KO models were putative EHF targets repressed in the normal prostate. Indeed, we found that EHF directly controlled a vast network of transcriptional regulators and downstream genes, including many EMT- and stemness-related factors. Interestingly, most of the deregulated TFs in EHF-KO mice (e.g., AP-1, Klf, Tcf family TFs) were among the key TFs known to be differentially activated in AR-independent CRPC subtypes. Furthermore, putative EHF target TFs based on genomic occupancy of regulatory sites were strongly associated with cell fate specification, morphogenesis, and development, consistent with the prominent role of EHF in controlling the expression of multiple regulators of cell plasticity, stemness, and differentiation in epithelial cells.

We also found that COL1A1, a critical ECM component and cell signaling mediator^30^, was among the direct targets of EHF and one of the top-activated genes in EHF-KO mice and both human and murine epithelial cells. COL1A1 expression is associated with stemness and pluripotency in prostate epithelial cells^32^. Furthermore, various TFs, including AP-1 family TFs, control COL1A1 expression in different contexts^32^. Here, we discovered a novel upstream regulatory mechanism leading to COL1A1 overexpression in prostate cancer. We found that EHF negatively impacted COL1A1 transcription in prostatic cells. Notably, we observed that EHF was directly bound to COL1A1 promoter regulatory sites and repressed its transcription. On the other hand, AP-1 TFs and other TFs (e.g., STAT3) controlling COL1A1 expression were also downstream targets of EHF and were activated upon EHF deletion. Thus, our data positions EHF upstream of both AP-1 TFs and other TFs and as a tight repressor of COL1A1 transcription.

Importantly, we found that COL1A1 was a key driver of phenotypic plasticity, epithelial-mesenchymal transition, and stemness in both human and murine EHF-depleted models. Genetic depletion of COL1A1 and pharmacologic inhibition of the COL1A1 receptor DDR1, which mediates its intracellular downstream effects, reduced stemness potential, reversed phenotypic transitions, and restored responsiveness to the AR antagonist enzalutamide. Intriguingly, inhibiting the COL1A1/DDR1 axis also affected the JAK/STAT3 signaling, which was activated in both human and murine models with EHF ablation. Activation of the JAK/STAT3 pathway has a central role in prostate cancer^6,34^. We show that genetic and pharmacological inhibition of COL1A1 signaling led to the inhibition of pSTAT3 in EHF-KO mouse organoids and human epithelial cells, indicating that COL1A1 contributed to the JAK/STAT3 activation in these models. Furthermore, inhibiting pSTAT3 with the JAK inhibitor ruxolitinib reproduced the effects on organoid growth and luminal differentiation seen by inhibiting COL1A1/DDR1. Thus, inhibiting either COL1A1/DDR1 or JAK/STAT3 signaling reverted epithelial cell plasticity and blocked the effects of EHF ablation, sustaining the relevance and therapeutic potential of these pathways.

Collectively, these data reveal a previously unrecognized link between the ETS factor EHF, multiple transcriptional regulators, and signaling pathways that sustain stemness, cell plasticity, and phenotypic transitions in prostate cancer. Furthermore, these cell-intrinsic alterations induced by EHF loss with activation of downstream pathways like inflammatory signaling, cytokine production, JAK/STAT3, and Col1A1/DDR1 are likely to affect immune and stromal cells in the surrounding microenvironment and contribute to a pro-tumorigenic and immuno-suppressive state. Together, these findings have relevant implications for drug discovery and cancer therapy. The critical pathways involved, like the COL1A1/DDR1 and JAK/STAT3 axis, are targetable with investigational or clinically approved drugs. In addition, the in vitro and in vivo human and mouse EHF-KO models described here could represent optimal experimental platforms to test innovative therapeutic strategies for restoring EHF function, reversing phenotypic plasticity, and restoring epithelial lineage integrity in prostate cancer. Here, we propose a central role for EHF in maintaining prostate epithelial cell differentiation and preventing cell plasticity and aberrant phenotypic transitions. Furthermore, this work opens new perspectives and fills an important gap in the understanding of the timing, mechanisms, and implications of loss of lineage identity and cell plasticity in prostate cancer and, perhaps, other epithelial cancers.

## Materials and Methods

### Generation of novel murine models Pb-Cre4; EHF^flox/flox^ (cEHF-KO) and Pb-Cre4;EHF^flo*x/flox*^;R26*^ERG^* (cEHF-KO/R26^ERG^) mice

Animal studies were approved by the Institutional Authority (Swiss Veterinary Federal Authority) and performed in accordance with national and international guidelines and regulations. cEHF-KO mice were generated using the Knockout-first allele, promoter-driven selection cassette approach (EUCOMM;(Ehf^tm1a(EUCOMM)Hmgu^). In loxP mice mutants, the critical exon (targeted region of EHF: exon 3), are flanked by loxP site which allows for excision with Cre recombinase. To generate prostate-specific EHF knockout mice (*Pb-Cre4; EHF^flox/flox^),* we crossed conditional ready null/knockout mice for the EHF (*EHF^flox/flox^*) with *Pb-Cre4* mice ^37^ to knockout *EHF* specifically in the prostate. The EHF mutation occurred upon crossing and was detected by end-point PCR performed with specific primers. The Pb gene is expressed in the prostatic epithelium post-puberty (the gene is androgen-responsive). Thus, excision of the EHF gene occurs only in the prostatic epithelium post-puberty. This avoids developmental effects due to the complete inactivation of EHF during prostate organogenesis. Rosa26^ERG/ERG^ (R26^ERG^) were obtained from Charles Sawyer’s laboratory and has been previously described^16^. To generate the double transgenic/knockout *Pb-Cre4;EHF^flox/flox^;R26^ERG^*, we crossed Rosa26^ERG/ERG^ (R26^ERG^) mice with *Pb-Cre4; EHF^flox/flox^*.

### Cell lines, plasmids and shRNA

Immortalized normal prostate epithelial RWPE-1 cells were maintained in Keratinocyte Serum-Free Growth Medium (KSF; 17005075 Thermo Fisher Scientific) with specific supplements ^11^. RWPE-1 cell lines with stable EHF knockdown (RWPE-1 shEHF/ESE3) and control RWPE-1 shCTRL were established and maintained as previously described ^11^. UGSM-2 cells were obtained from ATCC and maintained as previously described^25^. LNCaP-abl ^38^, were maintained in RPMI-1640 no-phenol red, supplemented with 10% charcoal-stripped serum (CSS) and 1% Pen/Strep. We stably expressed EHF in LNCaP-abl cells and generated the LNCaP-abl pESE3 cells. To this end, we employed a plasmid vector that comprises the open reading frame of EHF, which is transcribed under a pRC-CMV promoter^9^. The expression of EHF is maintained stable in the cells thanks to the presence of a neomycin/G418 (antibiotic resistance sequence)^39^.

To ablate COL1A1 in RWPE-1 cells we used lentiviral transduction with shRNA targeting human murine COL1A1 (clones ID number: NM_000088; TRCN0000062558 and TRCN0000062562).

### Multilineage Phenotypic Transition assay

RWPE-1 shEHF/ESE3 cells were cultivated in 2 different specific growth conditions to test their plasticity. For mesenchymal stimulation, cells were cultivated in Mesenchymal SC Growth Medium (MSCGM) BulletKit (PT-3238+PT-4105, Lonza) for 7 days. For neural stimulation, the following mediums was used: STEMdiff™ Neural Induction Medium (cat. number: 05835, StemCell Technologies) for 14 days. For the not stimulated condition (control group), RWPE-1 shEHF/ESE3 were cultivated in the KSF growth medium. At the end of stimulation, RNA was extracted for RNA-sequencing analysis.

### Sphere Forming Assay

For sphere-forming assay (SFA), cells were plated in Poly-HEMA (Poly-2-hydroxyethyl methacrylate, 1X) (Sigma-Aldrich) coated 6 well-plates in serum-free Mammary Epithelial Basal Medium (MEBM, Cambrex) supplemented with B27 Factor (1X), Follicular Growth Factor (20ng/ml), Epidermal Growth Factor (10ng/ml), Insulin (0.4mg/ml) and 1% Pen/Strep (Thermo Fisher Scientific, cat. 15070063). 1.000 cells/mL were seeded in SFA conditions. RWPE-1 spheres were counted after 7 days. Representative pictures were taken using Zeiss Microscope with Canon EOS 450D.

For ex-vivo SFA, single cells were seeded in precoated Poly-2-hydroxyethyl methacrylate 6-well plates at a density of 5.000 cells/ml in 2 ml of SFA medium (10.000 cells/well) in triplicate as previously described ^11^. After 14 days in culture, spheroids bigger than 50 µm in diameter were pictured and counted by optical microscope.

### Boyden chamber assay

For Boyden chamber assay, 50.000 cells/insert were plated in a Transwell® permeable support, in a 24-well plate with 5.0 µm polycarbonate membrane. RWPE-1 shEHF/ESE3 and shCTRL cells were plated in KSF without EGF and BPE and a regular growth medium was used as a chemoattractant. After 24 h, cell migration was blocked and migrating cells were stained with crystal violet in 0.01% distilled water for 10 minutes. The excess color was washed out with tap water. Cell migration was evaluated using ImageJ. Each experiment was carried out in triplicate.

### Vascular mimicry assay

Vascular mimicry assay was performed as follows: 18×18 mm glass coverslips (Cat. N°. 0101030, Marienfeld, Lauda-Königshofen, Germany) were ethanol-washed, air-dried, and placed in 12-well culture plates (Cat. N°. 140675, Thermo Scientific, Waltham, MA). After drying, coverslips were loaded with 50 μL/coverslip of Matrigel and air-dried for 45-60 min at room temperature. Prostatospheres derived from RWPE-1 shEHF/ESE3 were harvested, filtered to exclude structures below 40 μm, resuspended in 200 μL of KSF medium, and seeded onto matrigel-coated coverslips. They were incubated at 37°C for 1 hour and finally covered with 3 ml of KSF medium. Fresh medium was added every 3 days ^40^. Morphometric Image analysis for tubular-like structures quantification was performed using the plug-in “Angiogenesis Analyzer” software developed for ImageJ. The “Angiogenesis Analyzer” software was created in the macro language of ImageJ, as described in Carpentier et, al. ^41^.

### Hypoxia conditions and VEGF stimulation

To evaluate the impact of hypoxia on vasculogenic mimicry process, spheroids structures were generated in normoxic conditions and at day 3 post-seeding were seeded in vasculogenic mimicry assay, as previously described. Hypoxic conditions were set in a standard KSF medium with hypoxic exposure in a NUAIRE 5741E Air-Jacketed Automatic CO2 Incubator (NuAire, Inc. 2100 Fernbrook Lane Plymouth, Mn, USA) with 1.5% oxygen, 5% CO2, balanced nitrogen for 4 days. To test the consequences of VEGF-exposure in hypoxic condition in our RWPE-1 cellular model, vascular mimicry was performed in parallel using endothelial transdifferentiation medium adapted from a paper described by Zhao et al. ^42^. The transdifferentiation medium was composed as follows: Keratinocyte serum-free growth medium (KSF; 17005042, Gibco) supplemented with 2.5 μg EGF Human Recombinant (Cat. No. 10450-013), 25 mg Bovine Pituitary Extract (Cat. No.13028 -014), and 1% Penicillin/Streptomycin, VEGF (Invitrogen, Cat. No. 100-20-10UG, USA), 1/50 B27 supplement, 10 ng/ml EGF, 5 ng/ml bFGF (Sigma-Aldrich). For all conditions, fresh medium was added every 3 days. Morphometric analysis and quantification of tubular-like structures was performed after 7 days post-seeding, whereas branching phenotype was evaluated after 10 days post-seeding. Representative pictures were captured using Zeiss Microscope with Canon EOS 450D.

### Organoid establishment

To establish 3D organoids from GEMM and human cell lines, we followed Drost, J. et al. ^22^. In brief, murine single cells from GEMM prostates were obtained after tissue digestion and dissociation. After trypan blue viable cell counting, we seeded 240.000 cells resuspended in 120 μl of mouse prostate culture medium (MPCM) ^22^ and combined with 360 μl of Matrigel, Growth Factor Reduced (GFR), Phenol Red-free (BD, cat. no. 356231). Domes were prepared by combining 75% of Matrigel and 25% of MPCM to obtain a 480 μl of final volume for a total of 12 domes. Each dome was composed of 40 μl of combined Matrigel and cells. Each dome was seeded in the middle of one well of a 24-well plate and incubated for 1 hour at 37°C upside down. After dome solidification, 250 μl of MPCM was added to each well ^22^. After 4 days, MPCM was replaced with fresh one and approximately after 7 days, organoids were harvested from 12 domes of 24-well plate in the culture medium and transferred to a 15 ml Falcon tube. Organoids were separated from Matrigel matrix by trituration with a blue tip pipette. After pipetting up and down 15 – 20 times, 5 ml of ice-cold adDMEM/F12 +/+/+ was added to dissolve residual Matrigel matrix. Centrifugation at 150 g for 5 min at 4°C was performed, and the supernatant was aspirated to obtain a resuspended pellet for downstream application. Organoids were counted using an inverted microscope after 7-23 days, based on their growth kinetics and their morphologies were evaluated as previously described^25^. For selected experiments, organoids were stained with H&E and Ki67, and quantification was performed using APERIO. For human cell lines-derived 3D organoid experiments, we followed the same protocol described above with the following modifications. After trypan blue viable cell counting, human EHF-KD cells were seeded in Matrigel matrix domes and cultured in their KSF regular complete growth medium. To ablate COL1A1 in murine organoids we used spin-infection with shRNA targeting murine COL1A1 (clone ID number: NM_007742; TRCN0000090503).

### Drug treatments of organoids

GEMM-derived and human cell lines-derived cells were seeded in non-tissue treated 96-well plates as 5.000 cells/well and were grown following culture medium composition and volumes as described by Drost et al., ^22^. Each drug treatment condition was performed in quadruplicate as described above. Ruxolitinib (MedChem Express, cat numb. HY-50856), Enzalutamide (MDV-3100) MedChemExpress, cat. Numb. HY-70002), DDR1i (DDR1-2, SELLECKCHEM, cat. numb. S6817), and DHT (5-alpha-Dihydrotestosterone, sigma Aldrich, cat. numb. D-073-1ml) were dissolved in DMSO and were added to each well in the correct dilution in 100 μl of mouse prostate culture medium (MPCM).

### EHF/ESE3 DNA supplementation to murine organoids

WT, cEHF-KO, and cEHF-KO/R26ERG 3D organoids were established in a 48-well plate as described above. After 8 days, murine organoids were transfected with EHF/ESE3 plasmid (pESE3) and control plasmid (pCMV) using the jetPRIME® transfection kit (Polyplus) following manual instructions. 2 µg of DNA, 50 µl of jetPRIME® buffer and 2 µl of jetPRIME® reagent were used. Organoids count and morphology were evaluated 3 and 5 days post-transfection using a Leica Microscope.

### Mice for subcutaneous allografts

NOD.Cg-*Prkdc^scid^ Il2rg^tm1WjI^*/SzJ (NSG) male, mice used to establish subcutaneous allografts of mouse-derived prostatic cells, were purchased from Charles River Laboratories. Transgenic *R26^ERG^* (ERG) and Transgenic/knockout *Pb-Cre4;Pten^flox/flox^;R26^ERG^* (ERG/PTEN) mice with combined prostate-specific deletion of PTEN and overexpression of ERG were described previously ^16^.

### Flow cytometry

All steps for flow cytometry were performed in PBS supplemented with 0.5% BSA, and 2 mM EDTA. For analysis, single cells derived from GEMM prostate, were stained with anti-human CD45 (1:500, CD45 APC-Cy7, clone 30-F11, cat. number 2255516, Invitrogen), anti-human CD31 (1:500, FITC, Monoclonal Antibody (390), cat. number 2175435, Invitrogen) and EpCAM (1:200, PE-Cy7, G8.8, cat. number 2295626, Invitrogen). Samples were measured with a FACS Fortessa (BD Biosciences) and analyzed with FlowJo 10.8.0 software. For stem/progenitors fractions evaluation, single cells derived from GEMM prostate were stained for Lineage negative (CD45, CD31, CD49f (Thermo Fisher Scientific, clone eBioGoH3, dilution 1:200), Sca-1 (Biolegend, clone D7, dilution 1:300) and Trop2 (Bio-Techne AG, cat. number FAB1122A, dilution 1:100) antibodies.

### Establishment of in vivo subcutaneous allografts from mouse prostate-derived cells

Prostates from WT and cEHF-KO mice were dissected, minced into small pieces, and incubated in Hanks Balanced Salt solution (Sigma: cat. number H9394) supplemented with Collagenase Type 1 (1 mg/ml, Sigma: cat. number C0130) for 1 hour. Cell suspensions were passed through a 40 µm cell strainer (Falcon: cat. number 352340) to collect single cells. Cells were centrifuged (2000 rpm for 5 min), washed twice in PBS1X and counted by trypan blue for viability. Next, an equal number of viable cells were mixed (ratio 1:1) with Matrigel matrix and UGSM-2 cells (ratio 1:4): 200.000 cells combined with 50.000 UGSM-2 cells and subcutaneously injected in the flanks of NSG male mice for tumor growth in vivo.

### Infection of mouse-derived cells with shRNA lentivirus and establishment of in vivo subcutaneous allografts

To ablate EHF in WT and ERG transgenic background (Rosa26-ERG^flox/flox^ and Rosa26-ERG^flox/flox^/PTEN^flox/flox^ we used spin-infection with shRNA targeting murine EHF (clones ID number: NM_007914.3-513s21c1 and NM_007914.3-774s21c1). Briefly, Rosa26-ERG^flox/flox^ and Rosa26-ERG^flox/flox^/PTEN^flox/flox^-derived mice prostates were dissociated as described above. Viable single cells were divided into 3 polystyrene FACS tubes in a total volume of 500 μl per tube. 8 μg/ml (final concentration) of Polybrene (Sigma: cat. number <u>H9268</u>) was added to each tube. Next, the appropriate high-titer lentivirus (pLKO-CTRL shRNA and 2 mouse pLKO-EHF #231212 and #pLKO-EHF #257160) were used in a typical MOI (multiplicity of infection) in the range of 5-10. The tubes were placed at 37 °C in a sterile tissue culture incubator for 1 hour and mixed every 15 min by flicking. After 1 hour the tubes were centrifuged at 1800 rpm (754g) for 1 hour at 25 °C (spinfection) ^43^ ^23^. Tubes were washed three times with PBS 1X to remove any unbound lentivirus. Infected cells were seeded for ex-vivo SFA assay in T25 flasks. After 24 hours post-seeding in SFA, 1 µg/ml of puromycin (SIGMA; P8833) was added for 3 days to ensure only infected cells remained. Spheroids-forming infected cells cultured in SFA were counted for cells viability by trypan blue. Equal number of infected cells 200’000/injection were mixed (ratio 1:1) with Matrigel matrix and mixed with 50.000 UGSM-2 cells (ratio 1:4), for in vivo subcutaneous injections in NSG-KO mice and tumor growth was monitored by Caliper measurements.

### RNA extraction from mouse prostates

GEMM prostate glands were harvested and snap frozen in RNase-free conditions. Total RNA was extracted from GEMM prostates as described in following details. Briefly, the entire mouse prostates were disrupted with pestle (Axygen PES-15-B-SI Disposable Tissue Grinder Pestle) in 1.5 ml Eppendorf tubes loaded with 350 µl of RLT buffer (RNeasy® Mini Kit, Qiagen, cat. number 74104). Then, RNA was extracted by following RNeasy® Mini Kit instructions. The RNA quality was verified by the spectrophotometric analysis by using 2100 Bioanalyzer System (Agilent, cat. number G2939BA). Only samples with R.I.N. (RNA integrity number) 8-10 were used for Poly-A RNA sequencing analysis.

### RNA extraction and qRT-PCR

Total RNA was extracted after the resuspension of cells in Trizol (Ambition by Life Technologies, cat. number 15-596-026) with the Direct-zol RNA-MiniPrep kit (Zymo Research, cat. number R2051). All RNA samples were treated with DNase I (included in the kit from Zymo Research, cat. number R2051) to remove any contaminant genomic DNA. Elution of extracted RNA was performed using 35 μL of UltraPure DEPC Treated Water (Invitrogen, cat. number 10977035). Quantitative real-time PCR (qRT-PCR) was performed using 20 ng of RNA as template for SYBR Green RT-PCR one-step qRT-PCR kit (Qiagen, cat. number 204243) on the ABI 7000 machine (Applied Biosystem). Samples were analyzed in triplicate. The level of each gene was calculated by comparing the Ct value in the samples normalized to the amount of β-actin. Next, the relative quantities were normalized to the control samples. Sequences of all PCR primer sets used are shown in Key Resources Table.

### Immunoblotting

Cell lysates were prepared using RIPA buffer with protease inhibitor cocktail (cOmplete PIC, cat. number 4693116001, Roche) and phosphatase inhibitor cocktail (PhosStop, cat. number 4906845001, Roche). Total cell extracts were separated by SDS-PAGE and transferred to PDFV membranes (cat. number NBA085C001EA, PROTRAN). Antibodies directed to the following proteins were used for immunoblotting analysis: EHF (α-rat, 5A.5 Ab, provided by Antonio Tugores, 0.172 mg/mL, diluted 1:500), α-Tubulin (cat. number CP06, Calbiochem).

### Histological evaluation of mice and immunohistochemistry

Prostate sections from wild type (WT), heterozygous (cEHF-HET), homozygous cEHF-KO and double transgenic/knockout cEHF-KO/R26^ERG^ mice at different time points (from 8 to 36 weeks) have been stained with hematoxylin and eosin (H&E) and evaluated by trained pathologists for histological lesions.

Immunohistochemistry (IHC) on histological mice prostates tissues samples was performed using an antibody detecting EHF (1:25 dilution, Abcam, cat. number ab105375 Rb Polyclonal), AR (1:300 dilution, Abcam, cat. number ab133273 Rb Monoclonal), CHA (1:25 dilution, Abcam, cat. number ab15160 Rb Polyclonal), anti-ERG (Abcam, cat. number ab92513), anti-Ki67 antibody (Lab Vision Corporation, Clone SP6, cat. number RT-9106-R7), anti-Cytokeratin 5 [EP1601Y] (1:400 dilution, cat number ab52635, Abcam), anti-Cytokeratin 8 (1:400 dilution, cat. number ab59400, Abcam). The specificity of the antibodies was previously confirmed by western blot analysis. Cell nuclei were counterstained with a hematoxylin solution. Positive samples for each antibody and negative samples, in which the primary antibody was omitted, were used as controls. APERIO Software was used for quantification.

### Immunofluorescence

For immunofluorescence, prostate samples were fixed in 10% formalin (Thermo Scientific, cat. number 5701) and embedded in paraffin. Once dried, the sections were treated with OTTIX plus solution (Diapath, cat. number X0076) and OTTIX shaper solution (Diapath, cat. number X0096) to dewax and rehydrate the sections. Antigen retrieval was performed using pH 6 solutions at 98°C for 20 mins and non-specific binding sites with Protein-Block solution (DAKO Agilent technologies, cat. number X0909) for 10 min. Sections were stained with anti-Cytokeratin 5 [EP1601Y] (cat. number ab52635, Abcam) 1:400 antigen retrieval pH9 EDTA, anti-Cytokeratin 8 (cat. number ab59400, Abcam) 1:400 pH9 EDTA, anti-CD326 (EpCAM) Antibody (Thermo Scientific, cat. number 14-5791-81) 1:200 and CD31 (PECAM-1) anti-rat/mouse (R&D System, cat. number AF3628) 1:50 for 1h at room temperature, followed by 3 washes with 1X PBS-T. Alexa Fluor 594-conjugated anti-rat IgG secondary antibody (Thermo Scientific, cat. number A-21209, dilution 1:400), Alexa Fluor 488-conjugated anti-goat IgG secondary antibody (Thermo Scientific, cat. number A-110555, dilution 1:400), were incubated at room temperature for 30 min. To remove the autofluorescence the samples were treated with Ready Probes Tissue Autofluorescence

Quenching Kit (Invitrogen, cat. number R37630) and then the slides were mounted with ProLong Gold antifade reagent with DAPI (Invitrogen, cat. number P36931). For EGFP staining, the same protocol above described was used. For COL1A1 (E8F4L) XP® Rabbit mAb (Cell Signaling, cat. number 72026) 1:200 was used for 1h at RT, followed by 3 washes with 1X PBS-T. Alexa Fluor 488-conjugated anti-rabbit IgG secondary antibody (Thermo Scientific, cat. number A-21202, dilution 1:400).

### Quantification of the Immunofluorescence (IF) stain

For quantification of the Immunofluorescence (IF) stain in mice, images (n=5/7 for each genotype) were selected. Epithelial cells were identified for morphology and for positivity to EpCAM protein.

Quantification was performed using Fiji ImageJ software and the PEARSON’S COEFFICIENT plug-in (BIOP-BIOP-JACoP Plug-in); the following steps were applied: 1-click open LIF file and split channels for the desired image; 2 – open LIF file for red channel; 3-open LIF file for green channel; click Image, then color, then merge channels; 4-depict desired area to be analyzed for the Region of Interest (ROI); 5-click plugins, then BIOP, then Image Analysis, then BIOP JCACoP; 6-Pearsons’s Coefficient values were reported in excel file.

### Immunofluorescence on organoids and spheroids

Ibidi slides (µ-Slide, 18 well ibiTreat) were used for the establishment of Matrigel embedded organoid. Formed organoids were fixed by adding formaldehyde 4% (100 µL/well) on ice at 4°C for 3 hours. Then, formaldehyde was removed and 1.5 mg/ml glycine in PBS 1X at RT was added at 4°C until staining. Staining was performed by permeabilizing and blocking 30 min with 0.3 % Triton-X in 3 % BSA, at RT. Primary antibody dissolved in 0.3% Triton-X in 3% BSA was incubated at 4°C for ON or 48 hours. Organoids were washed once with BSA 3% 5 min at RT and secondary antibody were incubated for 45 min protected from light in 0.05% Triton-X+BSA 3% at RT. BSA 3% for 5 min at RT was used for washing once and PBS 1X was added with diluted Hoechst in 1:1000 for 1 min in PBS protected from light. For phospho STAT3 (Y705) anti-Rabbit mAb (Cell Signaling, cat. number 9131S) 1:100 was used for 48h at RT, followed by 3 washes with 1X PBS-T. Alexa Fluor 488-conjugated anti-rabbit IgG secondary antibody (Thermo Scientific, cat. number A-21202, dilution 1:500). For COL1A1 (E8F4L) XP® Rabbit mAb (Cell Signaling, cat. number 72026) 1:200 was used. Alexa Fluor 594-conjugated anti-rat IgG secondary antibody (Thermo Scientific, cat. number A-21209, dilution 1:400). Images were acquired by a confocal microscope Leica SP5, with an oil-immersion objective (633/1.4 NA Plan-Apochromat; Olympus), using laser excitation at 405, 488, or 594 nm. Images were processed using ImageJ software. Confocal images were obtained with the Leica TCS SP5 confocal microscope using 3 10/1.25 oil. Three to five fields of view from three independent samples were analyzed. The total area intensity signal was calculated using the analyze-analyze particles ImageJ software. Quantification of selected markers was performed as described in the previous paragraph.

### RNAScope

RNAScope for Col1A1, EpCAM and PECAM-1 was carried-out according with manufacturer’s protocol (RNAScope Multiplex Fluorescent Reagent Kit v2 Assay). Briefly, after organs paraffin inclusion, sequential sections of 4 μm were cut and baked in a dry oven at 60[for 1 hour. Slides were deparaffinized using two steps of 5 min in OTTIX Plus (Diapath, cat. number X0076) and one Step of 5 min in OTTIX Shaper (Diapath, cat. Number X0096). For the pretreatment steps, sections were covered for 10 min at room temperature with RNAScope Hydrogen Peroxide, incubated for 15 min in RNAScope Target retrieval reagent (ACDBio, cat. number 322000) and then treated 30 min at 40[with RNAScope Protease Plus (RNAscope H2O2 and protease reagents ACDBio, cat. number 322381). A mix of the probes for Col1A1-C1 (RTU ACDBio, cat. number 319371), EpCAM-C2 (1:50 dilution, ACDBio, cat. number 418151-C2) and PECAM1-C3 (1:50 dilution, ACDBio, cat. number 316721-C3) was prepared according to the manufacturer’s protocol and incubated with the samples for 2h at 40[. The following steps of Amplification and development of the signal were also performed following the RNAScope Multiplex Fluorescent Reagent Kit v2 Assay protocol (ACDBio, cat. number 323110). Fluorophores were TSA Vivid 520 for Col1A1 (1:1000 dilution, ACDBio, cat. number 323271), TSA Vivid 570 for EpCAM (1:1500 dilution, ACDBio, cat. number 323272) and TSA Vivid 650 for PECAM1 (1:1500 dilution, ACDBio, cat. number 323273). The slides were mounted with ProLong Gold antifade reagent with DAPI (Invitrogen, cat. number P36931).

### Bulk RNA sequencing

For RNA sequencing we used mouse prostate from WT (n=5), cEHF-KO (n=6) and cEHF-KO/R26*^ERG^*(n=4) 21-24 weeks-old. RNA was marked with Illumina total prep 96 RNA amplification kit (Ambion). RNA sequencing for all experiments was performed using Next Ultra II Directional RNA Library Prep Kit for Illumina starting from 800 ng of total RNA from each sample and sequenced on the Illumina NextSeq500 with single-end, 75 base pair long reads. After the evaluation of sequencing read quality was accessed through FastQC (v.0.11.9) ^44^, reads were aligned to the GRCm38.p6 release of the mouse genome (mouse samples) and to GRCh38.p14 release of the human genome (cell lines). The alignment and quantification were performed using STAR aligner (v.2.6.1c) in two-pass mode ^45^.

### Single-cell RNA sequencing (10X scRNA sequencing). Preparation of the samples

For single-cell RNA sequencing we used mouse prostate from WT (n=2), cEHF-KO (n=2) and cEHF-KO/R26*^ERG^* (n=2) 21-24 weeks-old. Prostates were dissected in cold PBS1X containing RNAi (ratio 1:1000) Superase RNase inhibitor (Invitrogen, cat. Numb. AM2694) in cold small petri dish (Nunc, cat. Numb. 150462) and disrupted mechanically in small pieces with blades. The minced tissues were moved to 3 ml of digestion medium: RPMI-1640 (SIGMA, cat. Numb. R 0883-500ML; 10% FBS; Pen/Strept 1%; HEPES 25 mM (SIGMA, cat numb. H0887; Collagenase D, 1 mg/ml (SIGMA, cat. Numb. 11088858001); DNase I Solution (2500 U/mL, cat numb. 90083). Tissues were incubated for 50 min at 37°C on orbital shaker. After, digestion was blocked by adding 3 ml of stopping buffer (RPMI-1640 with 10% FBS and 1% Pen/Strept). Samples were homogenized with a 1 ml pipette and filtered through a 100 µm nylon mesh and rinsed with 4 ml of RPMI-1640 medium. Cell suspension was incubated on ice for 4 min and then filtered through 40 µm nylon mesh strainer. After centrifugation at 1500 rpm for 5 minutes at 4°C, cells were resuspended in 200 µL of stopping buffer and vortexed in FACS sorting tubes (Falcon™ Round-Bottom Polystyrene Test Tubes with Cell Strainer Snap Cap, 5mL, cat. numb. 352235). Cell suspension was incubated for 5 minutes with 7-AAD (7-Aminoactinomycin D, cat. numb. A1310) before sorting was performed with FACS ARIA (BD). At the end of sorting, viable sorted cells were collected in low binding 1.5mL eppendorf tubes pre-coated with BSA 3% and 10 µl RNAi and 15 µl of RPMI-1640 with FBS 10%. To sequence the transcriptome, the recommended amount of cell suspension was loaded into 10x Chromium single cell Controller (10× Genomics, Pleasanton, CA. USA), and the Chromium Next GEM Single Cell 3’ v3 Reagent Kit was adopted. All the procedures were performed following the manufacturer’s instructions reported in the Chromium Next GEM Single Cell 3’ guide.

### Downstream analysis of bulk transcriptomic data

RNA-Seq analysis was carried out using the DESeq2 pipeline in R statistical environment (DESeq2 v1.36.0, R v.4.2.1) ^46^. Differential expression analysis was performed using the Independent filtering procedure to discard genes expressed at low levels; to normalize data, the variance stabilizing transformation method (vst) was applied to the raw count matrix ^47^. Computed p-values were adjusted for multiple testing using the Benjamini-Hochberg (FDR) correction method and the subsequent gene-set enrichment analysis was performed with CAMERA algorithm (limma package v.3.52.3 ^48^: specifically CAMERA was run in pre-ranked mode, where input genes were sorted according to the Wald statistic (DESeq2 package). All tested gene sets were retrieved from the Molecular Signature Database (MSigDB). To functionally annotate custom gene lists we adopted the enrichR package v.3.1^49^. Single-sample enrichment analysis was performed with GSVA with ssgsea mode.

### Analysis of Single-cell RNA sequencing data

Gene expression quantification. Raw sequencing data were demultiplexed with cellranger mkfastq (cellranger v.6.0.0), and input reads were subsequently aligned and quantified at gene level using a custom-mouse reference genome (GRCm39 vM28 + EGFP + TMPRSS2-ERG FUSION) through cellranger count function, resulting in the generation of a gene-cell matrix in which Gene Symbols were used as identifiers.

#### Data pre-processing and filtering

Data were analyzed with Seurat (v.4.1.1) ^50^ in R statistical environment ^46^. Samples were individually analyzed to assess the good quality of sequenced reads and to detect and discard transcriptional outliers. To pre-process the data, the following steps were applied for each sample: normalization of the data (Seurat::NormalizedData, default parameters), identification of the most variable features (Seurat::FindVariableFeatures, default parameters), data scaling (Seurat:: ScaleData, default parameters), PCA generation, clustering (Seurat::FindCluster, with 10 different resolutions from 0.1 to 1). Clusters containing high fraction of cells with high expression of mitochondrial, ribosomal, hemoglobin genes or with low counts were discarded. Furthermore, the same parameters were exploited to compute the skewness-adjusted Multivariate Outlyingness (robustbase::adjOutlyingness) for each cell, and the median-absolute deviation was considered to identify outlier cells. To predict the presence of doublets, DoubletFinder method ^51^, that includes pK Identification and the Homotypic Doublet Proportion Estimate, was applied according to the guidelines. Empty-droplet identification was done with DropUtils package (v.1.18.1). After cell outlier filtering, samples were merged in a Seurat Object and were pre-processed again starting from raw counts. Then, genes that were expressed in less than 10 cells were discarded, resulting in 21187 final genes and 24312 cells.

### Data processing and integration

Before performing the integration, data were normalized (Seurat::NormalizeData) and then, ratio of mitochondrial, ribosomal, and hemoglobin genes were regressed out in a second non-regularized linear regression (Seurat: SCTransform). Integration of 5 samples (2 WT, 2 cEHF-KO, 1 R26*^ERG^*) was performed through the identification of the 3000 most variable features and the definition of anchors (Seurat Select Integration Features, Seurat Find Integration Anchors, Seurat Integrate Data, normalization.method = SCT). Next, we computed the principal component analysis up to the top 50 components on the integrated Seurat Object. In order to establish the number of components to be exploited to determine the K-nearest neighbors of each cell (Seurat::FindNeighbors), we choose the minimum between 2 parameters that were computed as follow: (a) the PC that exhibits cumulative percent greater than 90% and a percentage of variation associated with the PC as less than 5; (b) the PC that differs from the previous with a percentage less than 0.1%. Next, we defined the clusters for 10 resolutions from 0.1 to 1 (Seurat::FindClusters) on the integrated object containing WT, cEHF-KO, R26*^ERG^* samples. The resolution set for the subsequent analyses was 0.3 and provided 17 clusters. The final object containing WT, cEHF-KO and R26*^ERG^* samples were used as reference for subsequent integration of the cEHF-KO/R26*^ERG^* samples. (sampleSeurat::MapQuery function (reference. reduction = pca, reduction.model = umap). Data were annotated for the cell types as described in the following paragraph. Then, the reclustering of epithelial, stromal and endothelial cells was carried out. For reclustering, data were processed starting from raw counts as previously described, except for the integration step, which was replaced by merging the data, to preserve differences among samples and highlight condition-specific gene expression patterns.

Cell-type annotations. UMAP was implemented to visualize all identified clusters. The cell types were associated with the clusters according to the expression of canonical markers (see Figure S4C). For epithelial cell type identification, we considered a range of canonical markers for basal and luminal epithelial cells previously described in details ^26,27^. To confirm the obtained assignment, other strategies were applied. Marker genes, specific to individual clusters and conserved across different samples (WT, cEHF-KO, R26*^ERG^* and cEHF-KO/R26*^ERG^*) were obtained (Seurat::FindConservedMarkers, only positive markers, logfc.threshold = 1) and enriched with enrichR (“Descartes Cell Types and Tissue” database)^49^. Moreover, predicted cell type of each individual cell was inferred with SingleR (v.6.1), which provides a labelled reference dataset containing 18 main cell types (celldex::MouseRNAseqData()) ^52^.

Differential gene expression analysis. Differential gene expression analysis was performed to compare one cluster to the others (considering all the samples) and to compare clusters of different mouse models. These analyses were performed with FindConservedMarkers (only.pos = TRUE, logfc.threshold = 1) and with FindMarkers (logfc.threshold = 0.7, test.use = “MAST”, min.pct = 0.1).

### Trajectory inference

Trajectory analysis was performed through Slingshot tool (v2.6.0) in R environment ^53^ on each genotype separately (WT, R26*^ERG^*, cEHF-KO, cEHF-KOR26*^ERG^*). Along the trajectory, individual cells are positioned based on the similarity of their gene expression profiles, which is synthesized in the “pseudotime” value. Pseudotime is a continuous and quantitative measure of the progression of cells along the reconstructed trajectory. The establishment of a trajectory implied two steps, which were wrapped in the slingshot function: (a) identification of lineage structure with a cluster-based minimum spanning tree (*getLineages* function); (b) construction of smooth representations of each lineage using simultaneous principal curves (*getCurves* function).

### CUT&RUN

For CUT&RUN, the protocol published by Skene and colleagues was followed ^54^. Briefly, LNCaP-abl pESE3 cells were harvested and 300,000 cells/replicate were used. Following the washing steps, cells were incubated overnight at 4°C with anti-EHF (ab105375, Abcam) or IgG (Millipore) antibodies, at a dilution of 1:100. The following day, the cells and antibody mixture were incubated with protein A-MNase fusion protein for 10 minutes at room temperature. Next, digestion was carried on for 30 minutes at 0°C. Finally, the CUT&RUN fragments were released, and DNA extraction was performed using NucleoSpin Gel and PCR Clean-up kit (cat. 740609.250, Macherey-Nagel), according to manufacturer’s instructions.

### CUT&RUN Sequencing Data Processing and Analysis

CUT&RUN sequencing data were processed using the nf-core/cutandrun pipeline (v3.2.2) [https://nf-co.re/cutandrun/3.2.2/], a standardized Nextflow-based workflow for reproducible analysis of CUT&RUN experiments. Raw paired-end FASTQ files were trimmed to remove adapters and low-quality bases using Trim Galore, followed by alignment to the human reference genome (e.g., GRCh38) using Bowtie2 with parameters optimized for short fragment alignment. Duplicate reads were removed, and quality control metrics (including fragment size distribution, read duplication rates, and signal-to-noise estimation) were computed using standard tools embedded within the pipeline, including samtools and deepTools.

Peak calling was performed using SEACR in relaxed mode, with matched IgG controls used for normalization. The resulting peak sets were filtered based on signal enrichment and reproducibility across biological replicates. For downstream differential binding analysis, peak counts were quantified and analyzed using the DiffBind package (v3.16.0) in R (v4.4.1). Peaks were merged across samples to generate a consensus peakset, read counts were normalized and analysed using DESeq2 method. Statistical testing for differential binding was performed using a negative binomial model, with differentially bound regions identified based on a false discovery rate (FDR) < 0.05. Annotation of peaks to nearby genes was performed using the ChIPseeker (v1.42.1) and TxDb.Hsapiens.UCSC.hg38.knownGene (v3.20.0) packages.

### Public genomic data

We took advantage from an integrated database of prostate cancer patients including 174 normal samples, 714 primary tumors, 316 CRPC, and 19 CRPC with neuroendocrine features ^18^. VST-normalized expression data along with its annotations were downloaded from Zenodo repository (https://doi.org/10.5281/zenodo.5546618). To summarize the expression of lists of genes in patient samples, the single sample Gene Set Enrichment Analysis (ssGSEA) through the gsva function was adopted.

### Management of single-cell RNA sequencing data from patient databases

To inspect the transcriptome profile at single-cell level of human samples, we interrogated scRNA seq. datasets. Data from patient datasets were analyzed following the same procedures applied for mouse models, with minor adjustments. Features and barcode information, along with the matrix with the raw counts were retrieved from GEO database (data accessible at NCBI GEO database, accession Primary tumors (GSE157703;GSE181294;GSE193337), CRPC (GSE137829; GSE143791; GSE210358) and NE (GSE210358) ^55^. Samples included, normal adjacent tissue (n=25), primary tumor (n=25), CRPC (n=22) and NEPC (n=3). The pre-processing was carried out as previously described, except for the supervised step, which was deemed unnecessary due to the good quality of the samples; hence, only the empty droplet detection, the unsupervised outlier detection, and the doublet detection steps were done. After the filtering procedure, data were merged, and the clusters were defined with 0.1 resolution. The assignment of cell type was performed with supervised and unsupervised methods, leading to the identification of the 4 broad cellular lineages: epithelial cells, endothelial cells, fibroblasts, and immune cells.

### Statistics and reproducibility

For all comparisons between two groups of independent datasets, two-tailed unpaired t-test, Wilcoxon rank-sum test or Dunn test were performed, p-value, standard deviation (SD) or standard error of the mean (SEM) were calculated. For all comparisons among more than two groups, one-way, two-way ANOVA or Dunn tests followed by multiple comparison were performed. For quantification of positive cell density, images of tumor sections with IF or IHC staining were captured by microscope. Three fields for each sample were randomly selected for positive cell density analysis and statistical analysis was performed by two-tailed unpaired t-test. The two-tailed Pearson correlation between murine samples for ERG, Ki67, EHF, Ck5, CK8 score were calculated using Graphpad Prism 8.0. Histological grade difference was determined by two-tailed c2 test. The predictive value and PFS were univariately analyzed using the Kaplan-Meier method. For genetically engineered mouse modes (GEMMs) analysis, the examinations were performed dependent on animal available. All the in vitro experiments were independently repeated at least three times (with at least 3 biological repeats in total), and the in vivo experiments included at least 4 biological repeats with a similar time course and treatment. Representative data from experiments performed at least in triplicate are shown.

### Materials availability

This study generated two novel prostate-specific mouse models: Pb-Cre4; EHF^flox/flox^ (cEHF-KO) and Pb-Cre4;EHF^flo*x/flox*^;R26*^ERG^* (cEHF-KO/R26*^ERG^*). The EHF^flox/flox^ mouse strain generated in this study has been deposited. The strain is archived and is currently listed on the INFRAFRONTIER website: https://www.infrafrontier.eu/emma/. Material (also derivatives) will be distributed under the conditions from the EUCOMM consortium (EUCOMM MTA).

## Data and code availability

All data and materials that support the findings of this study are available within the article and supplemental information. Supplemental figures and datasets are available as supplemental information. Raw data from RNA sequencing experiments related to RWPE-1 cell line and mouse models are available with the accession number E-MTAB-13229, respectively. Raw data from 10x S.C. sequencing are deposited with the accession number E-MTAB-13230. Any additional information required to reanalyze the data reported in this paper is available from the lead contact upon request.

## Ethical Approval

The research performed in this study complies with all relevant ethical regulations. All mouse experiments were approved by the Swiss Veterinary Federal Authority (license 35411; TI 32/3023) and performed in accordance with the national and international guidelines and regulations. All relevant animal use guidelines and ethical regulations were followed.

## Availability of data and materials

All data are available in the main text or the supplementary information. Any raw data supporting the conclusions of this article will be made available by the authors, without undue reservation.

## Acknowledgments.

The work was supported by the Swiss National Science Foundation (SNSF-310030_189081, SNSF-IZLSZ3_170898 and SNSF-310030L_170182), Swiss Cancer League (KLS-4569-08-2018 and KLS-4899-08-2019), Foundation Nelia and Amadeo Barletta (FNAB), Fondazione Ticinese Ricerca sul Cancro, and Fondazione San Salvatore. Graphical abstract and schematics were created with BioRender.com.

## Author contributions

D.A. performed experiments, analyzed the data, and wrote the manuscript draft. G.S. performed bioinformatics and statistical analysis and wrote the manuscript draft. C.M. performed tumor-spheroid and organoids experiments. E.S. assisted with mouse colonies, in vivo studies and data analysis. G.P. performed in vitro assays and VM assays in human cells and analyzed data. D.I. assisted in sample preparation and FACS. A.C. performed experiments and analyzed data. G.A.C. performed bioinformatics analysis of scRNA seq data from human prostate tumors. A.B. assisted with mouse colonies, in vivo studies. S.M. assisted in histological sample preparation and performed immunofluorescence and RNAScope. G.C. assisted in in vivo studies and FACS. A.R. assisted in RNA samples preparation and performed RNA-Seq, scRNA-Seq and CUT&RUN. C.J. assisted in histopathological evaluation of mouse samples. R.M.H. supervised and performed histopathological evaluation of mouse samples. M.B. supervised all the bioinformatics analysis. C.V.C. supervised the experimental plans, interpreted results, provided insights, resources and funding for the project, wrote and reviewed the manuscript. G.M.C. conceived the project, supervised the experimental plans, mentored participants, analyzed and interpreted the results, provided resources and funding, and wrote and reviewed the manuscript.

