## Supplemental Information for "Loss of ESE3/EHF is sufficient to promote cell plasticity, transformation and androgen-independent status in the early stage of prostate carcinogenesis"

### Slide 1
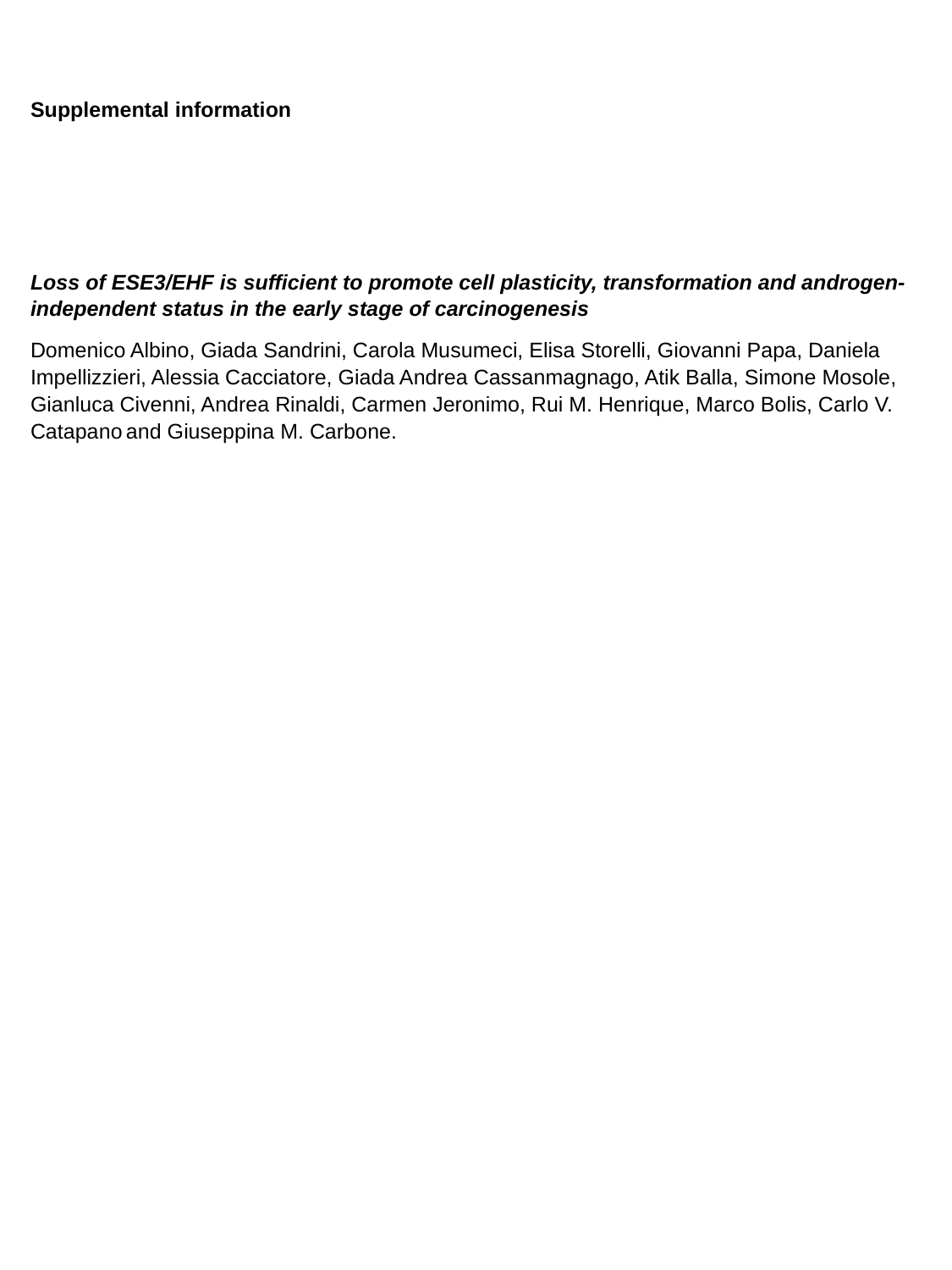

Supplemental information
Loss of ESE3/EHF is sufficient to promote cell plasticity, transformation and androgen-independent status in the early stage of carcinogenesis
Domenico Albino, Giada Sandrini, Carola Musumeci, Elisa Storelli, Giovanni Papa, Daniela Impellizzieri, Alessia Cacciatore, Giada Andrea Cassanmagnago, Atik Balla, Simone Mosole, Gianluca Civenni, Andrea Rinaldi, Carmen Jeronimo, Rui M. Henrique, Marco Bolis, Carlo V. Catapano and Giuseppina M. Carbone.

### Slide 2
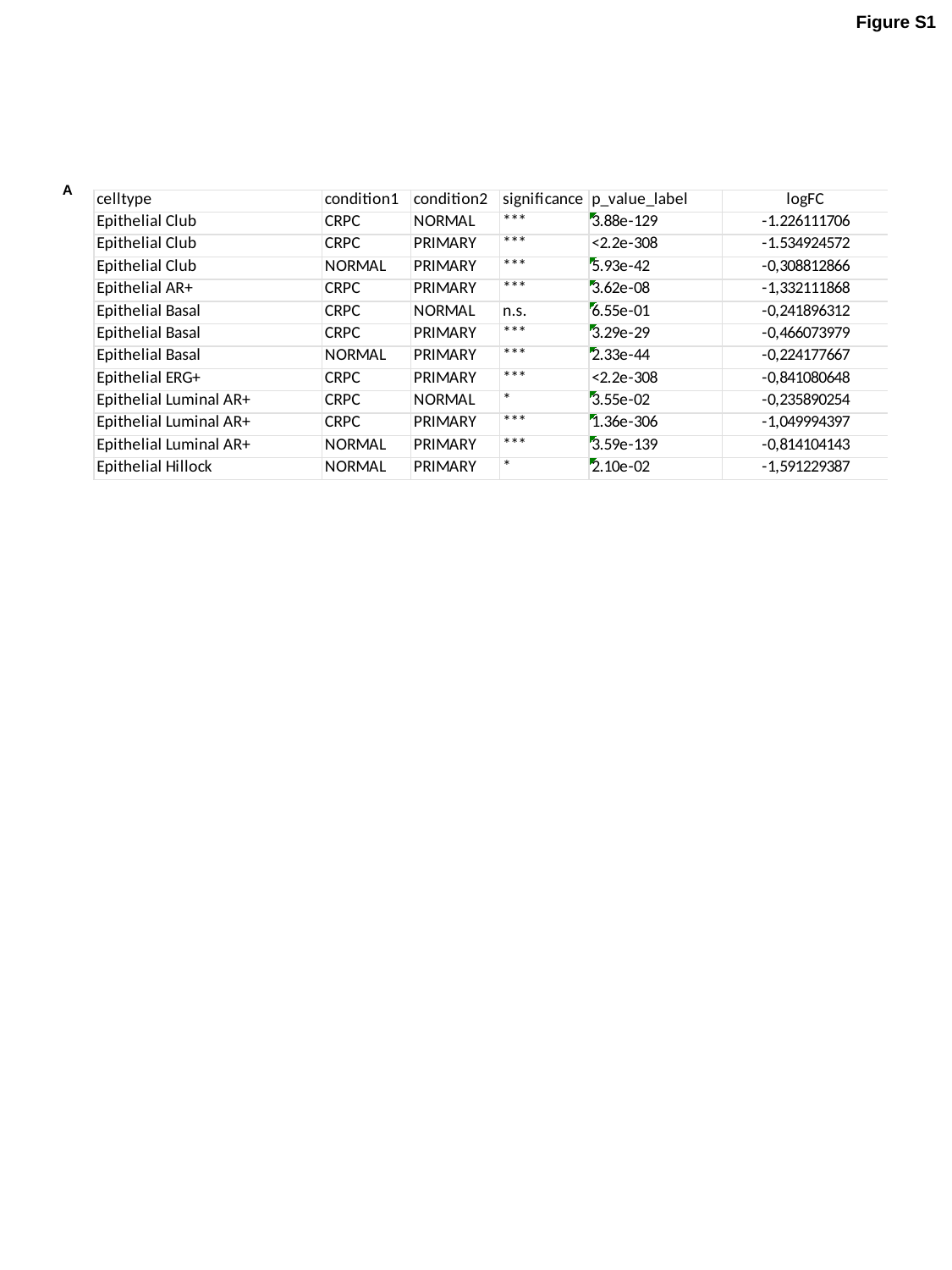

Figure S1
A

### Slide 3
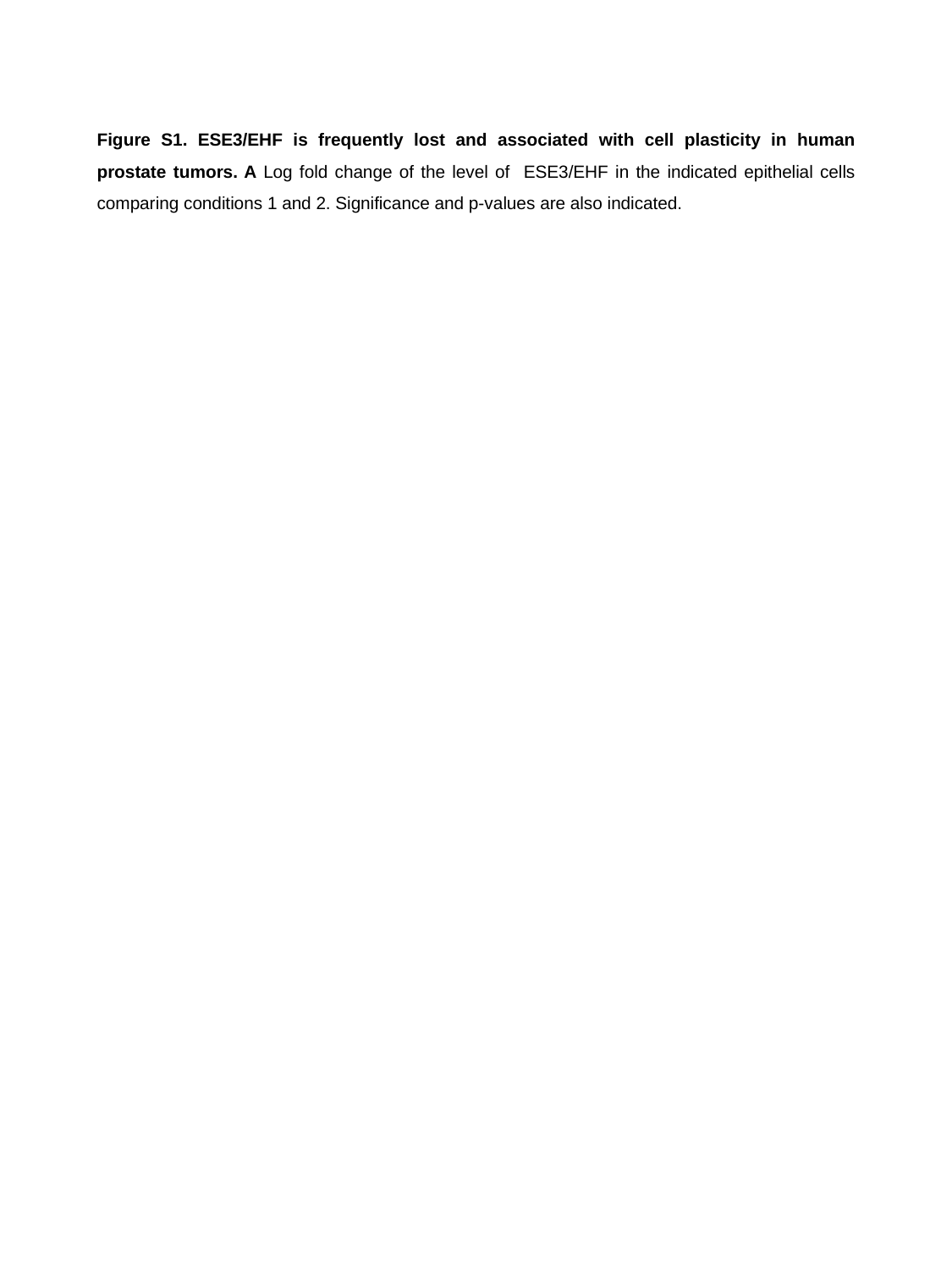

Figure S1. ESE3/EHF is frequently lost and associated with cell plasticity in human prostate tumors. A Log fold change of the level of ESE3/EHF in the indicated epithelial cells comparing conditions 1 and 2. Significance and p-values are also indicated.

### Slide 4
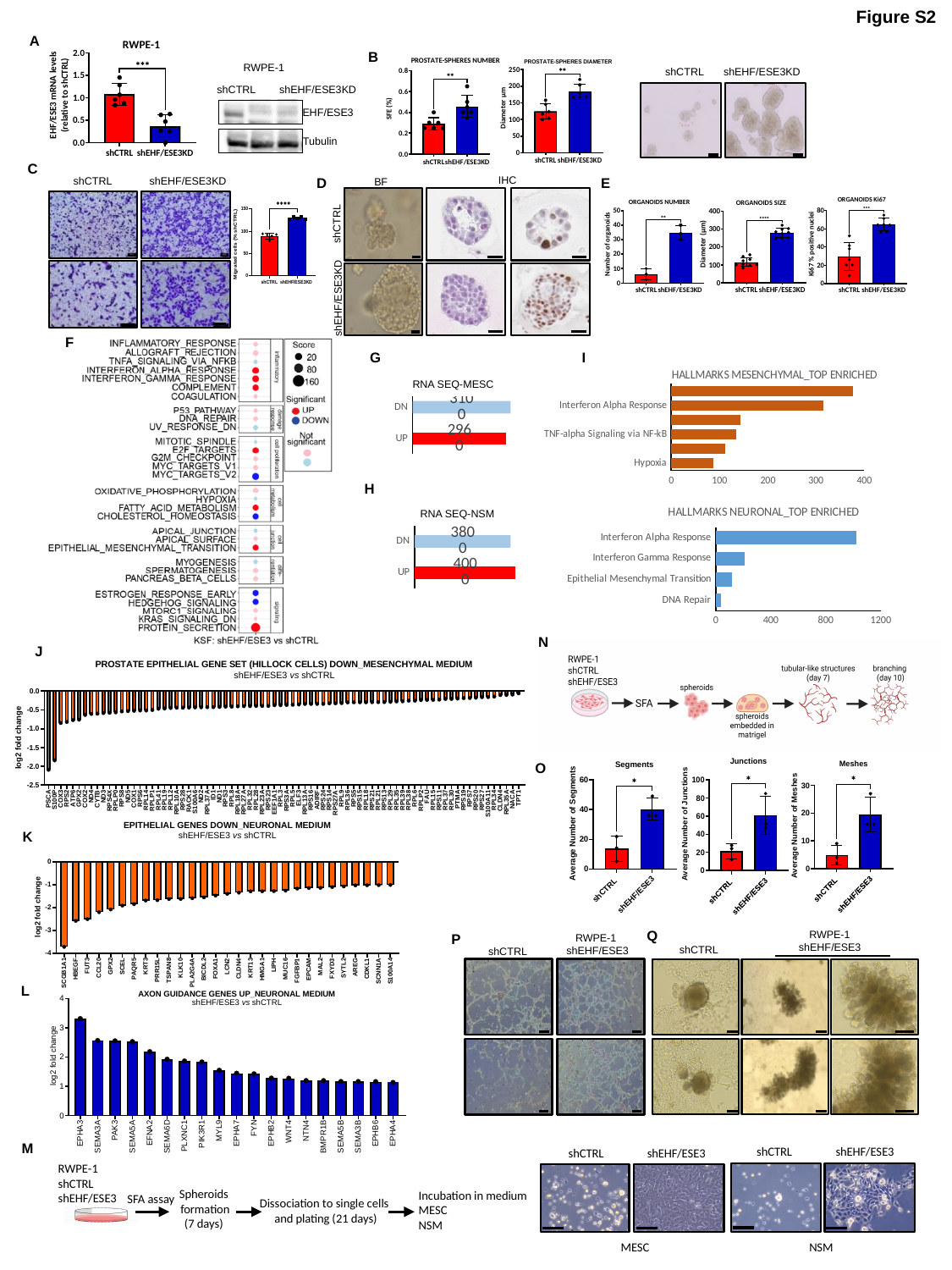

Figure S2
A
B
RWPE-1
shCTRL
shEHF/ESE3KD
EHF/ESE3
Tubulin
shCTRL
shEHF/ESE3KD
C
IHC
D
E
shCTRL
shEHF/ESE3KD
BF
shCTRL
shEHF/ESE3KD
F
G
I
#### Chart: HALLMARKS MESENCHYMAL_TOP ENRICHED
| Category | |
|---|---|
| Hypoxia | 87.77130557716295 |
| TGF-beta Signaling | 112.37697101135161 |
| TNF-alpha Signaling via NF-kB | 134.86054886440687 |
| Interferon Gamma Response | 142.86631995754703 |
| Interferon Alpha Response | 314.5760872543393 |
| Epithelial Mesenchymal Transition | 377.6500485170608 |RNA SEQ-MESC
#### Chart
| Category | |
|---|---|
| UP | 2960.0 |
| DN | 3100.0 |H
#### Chart: HALLMARKS NEURONAL_TOP ENRICHED
| Category | |
|---|---|
| DNA Repair | 37.60455269991727 |
| Epithelial Mesenchymal Transition | 118.0776945055203 |
| Interferon Gamma Response | 209.57738005257733 |
| Interferon Alpha Response | 1022.5610582569275 |RNA SEQ-NSM
#### Chart
| Category | |
|---|---|
| UP | 4000.0 |
| DN | 3800.0 |N
J
O
K
Q
RWPE-1
shEHF/ESE3
shCTRL
P
RWPE-1
shEHF/ESE3
shCTRL
L
M
shEHF/ESE3
shCTRL
shEHF/ESE3
shCTRL
MESC
NSM
RWPE-1
shCTRL
shEHF/ESE3
Spheroids
formation
(7 days)
Incubation in medium
MESC
NSM
Dissociation to single cells
and plating (21 days)
SFA assay

### Slide 5
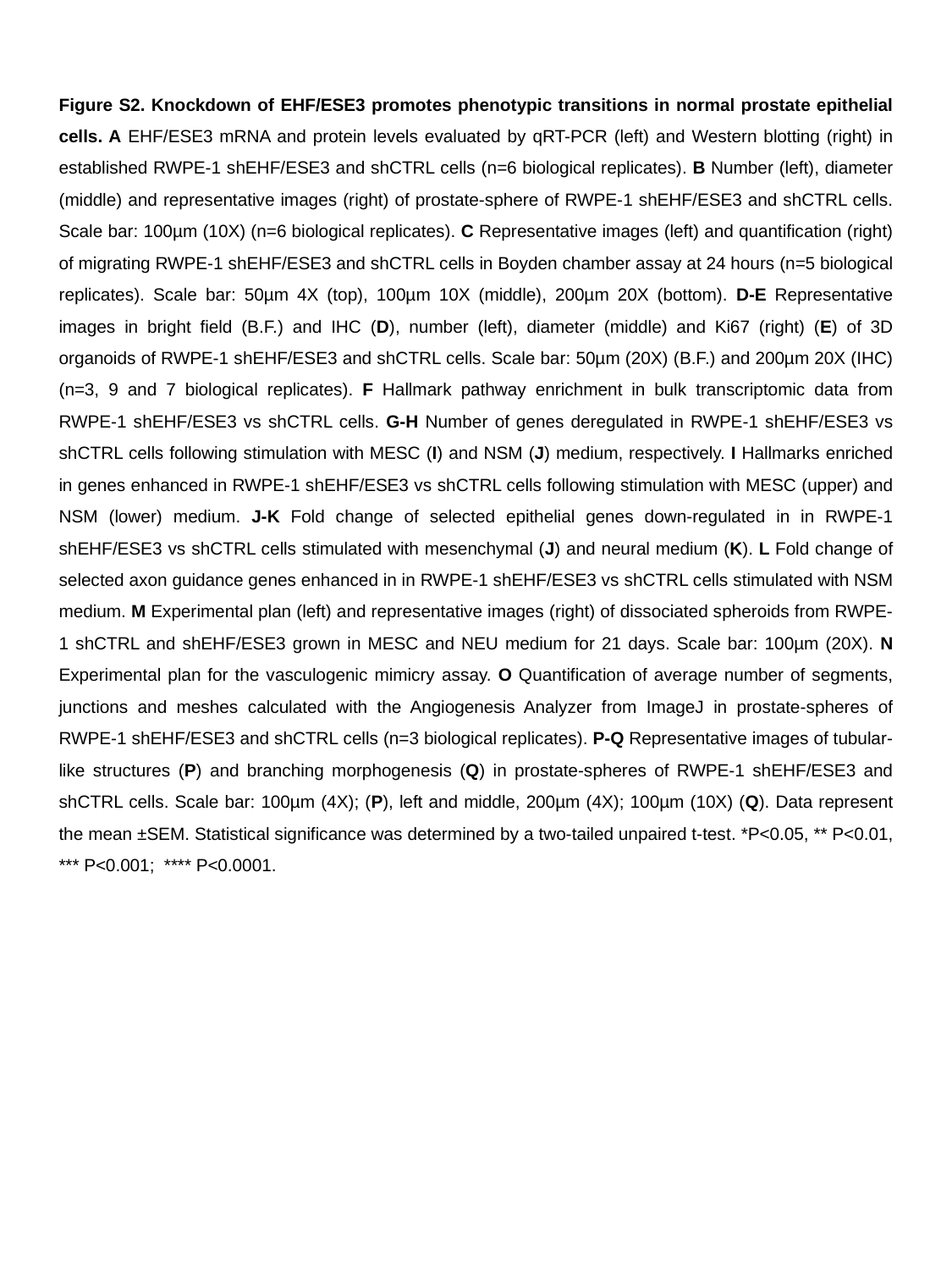

Figure S2. Knockdown of EHF/ESE3 promotes phenotypic transitions in normal prostate epithelial cells. A EHF/ESE3 mRNA and protein levels evaluated by qRT-PCR (left) and Western blotting (right) in established RWPE-1 shEHF/ESE3 and shCTRL cells (n=6 biological replicates). B Number (left), diameter (middle) and representative images (right) of prostate-sphere of RWPE-1 shEHF/ESE3 and shCTRL cells. Scale bar: 100µm (10X) (n=6 biological replicates). C Representative images (left) and quantification (right) of migrating RWPE-1 shEHF/ESE3 and shCTRL cells in Boyden chamber assay at 24 hours (n=5 biological replicates). Scale bar: 50µm 4X (top), 100µm 10X (middle), 200µm 20X (bottom). D-E Representative images in bright field (B.F.) and IHC (D), number (left), diameter (middle) and Ki67 (right) (E) of 3D organoids of RWPE-1 shEHF/ESE3 and shCTRL cells. Scale bar: 50µm (20X) (B.F.) and 200µm 20X (IHC) (n=3, 9 and 7 biological replicates). F Hallmark pathway enrichment in bulk transcriptomic data from RWPE-1 shEHF/ESE3 vs shCTRL cells. G-H Number of genes deregulated in RWPE-1 shEHF/ESE3 vs shCTRL cells following stimulation with MESC (I) and NSM (J) medium, respectively. I Hallmarks enriched in genes enhanced in RWPE-1 shEHF/ESE3 vs shCTRL cells following stimulation with MESC (upper) and NSM (lower) medium. J-K Fold change of selected epithelial genes down-regulated in in RWPE-1 shEHF/ESE3 vs shCTRL cells stimulated with mesenchymal (J) and neural medium (K). L Fold change of selected axon guidance genes enhanced in in RWPE-1 shEHF/ESE3 vs shCTRL cells stimulated with NSM medium. M Experimental plan (left) and representative images (right) of dissociated spheroids from RWPE-1 shCTRL and shEHF/ESE3 grown in MESC and NEU medium for 21 days. Scale bar: 100µm (20X). N Experimental plan for the vasculogenic mimicry assay. O Quantification of average number of segments, junctions and meshes calculated with the Angiogenesis Analyzer from ImageJ in prostate-spheres of RWPE-1 shEHF/ESE3 and shCTRL cells (n=3 biological replicates). P-Q Representative images of tubular-like structures (P) and branching morphogenesis (Q) in prostate-spheres of RWPE-1 shEHF/ESE3 and shCTRL cells. Scale bar: 100µm (4X); (P), left and middle, 200µm (4X); 100µm (10X) (Q). Data represent the mean ±SEM. Statistical significance was determined by a two-tailed unpaired t-test. *P<0.05, ** P<0.01, *** P<0.001; **** P<0.0001.

### Slide 6
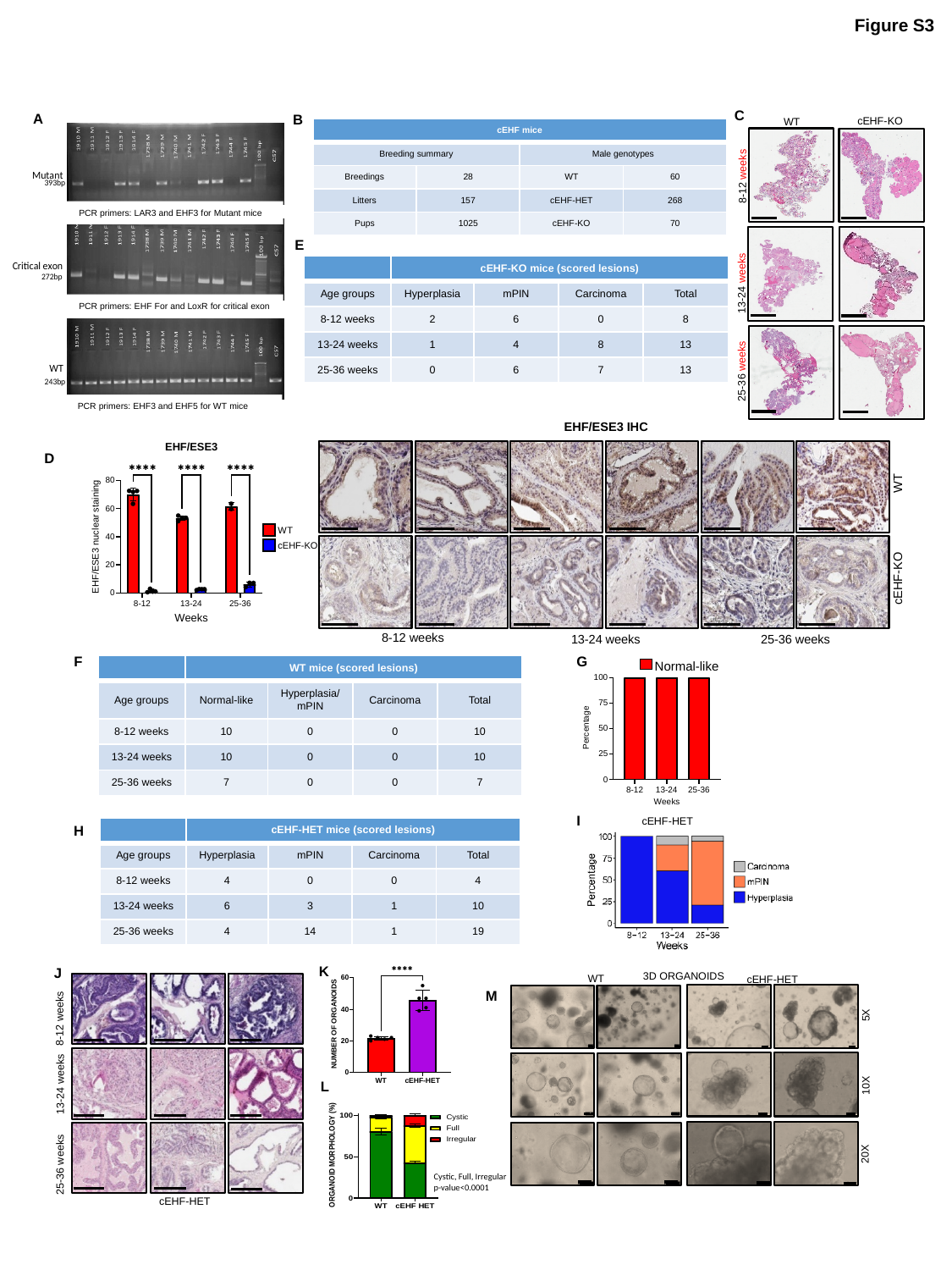

Figure S3
C
A
B
cEHF-KO
WT
8-12 weeks
13-24 weeks
25-36 weeks
| cEHF mice | cEHF mice | | |
| --- | --- | --- | --- |
| Breeding summary | | Male genotypes | |
| Breedings | 28 | WT | 60 |
| Litters | 157 | cEHF-HET | 268 |
| Pups | 1025 | cEHF-KO | 70 |
PCR primers: LAR3 and EHF3 for Mutant mice
E
| | cEHF-KO mice (scored lesions) | | | |
| --- | --- | --- | --- | --- |
| Age groups | Hyperplasia | mPIN | Carcinoma | Total |
| 8-12 weeks | 2 | 6 | 0 | 8 |
| 13-24 weeks | 1 | 4 | 8 | 13 |
| 25-36 weeks | 0 | 6 | 7 | 13 |
PCR primers: EHF For and LoxR for critical exon
PCR primers: EHF3 and EHF5 for WT mice
F
 EHF/ESE3 IHC
WT
cEHF-KO
8-12 weeks
25-36 weeks
 13-24 weeks
D
F
G
| | WT mice (scored lesions) | | | |
| --- | --- | --- | --- | --- |
| Age groups | Normal-like | Hyperplasia/mPIN | Carcinoma | Total |
| 8-12 weeks | 10 | 0 | 0 | 10 |
| 13-24 weeks | 10 | 0 | 0 | 10 |
| 25-36 weeks | 7 | 0 | 0 | 7 |
I
cEHF-HET
H
| | cEHF-HET mice (scored lesions) | | | |
| --- | --- | --- | --- | --- |
| Age groups | Hyperplasia | mPIN | Carcinoma | Total |
| 8-12 weeks | 4 | 0 | 0 | 4 |
| 13-24 weeks | 6 | 3 | 1 | 10 |
| 25-36 weeks | 4 | 14 | 1 | 19 |
K
J
8-12 weeks
13-24 weeks
25-36 weeks
cEHF-HET
3D ORGANOIDS
WT
cEHF-HET
5X
10X
20X
M
L
Cystic, Full, Irregular
p-value<0.0001

### Slide 7
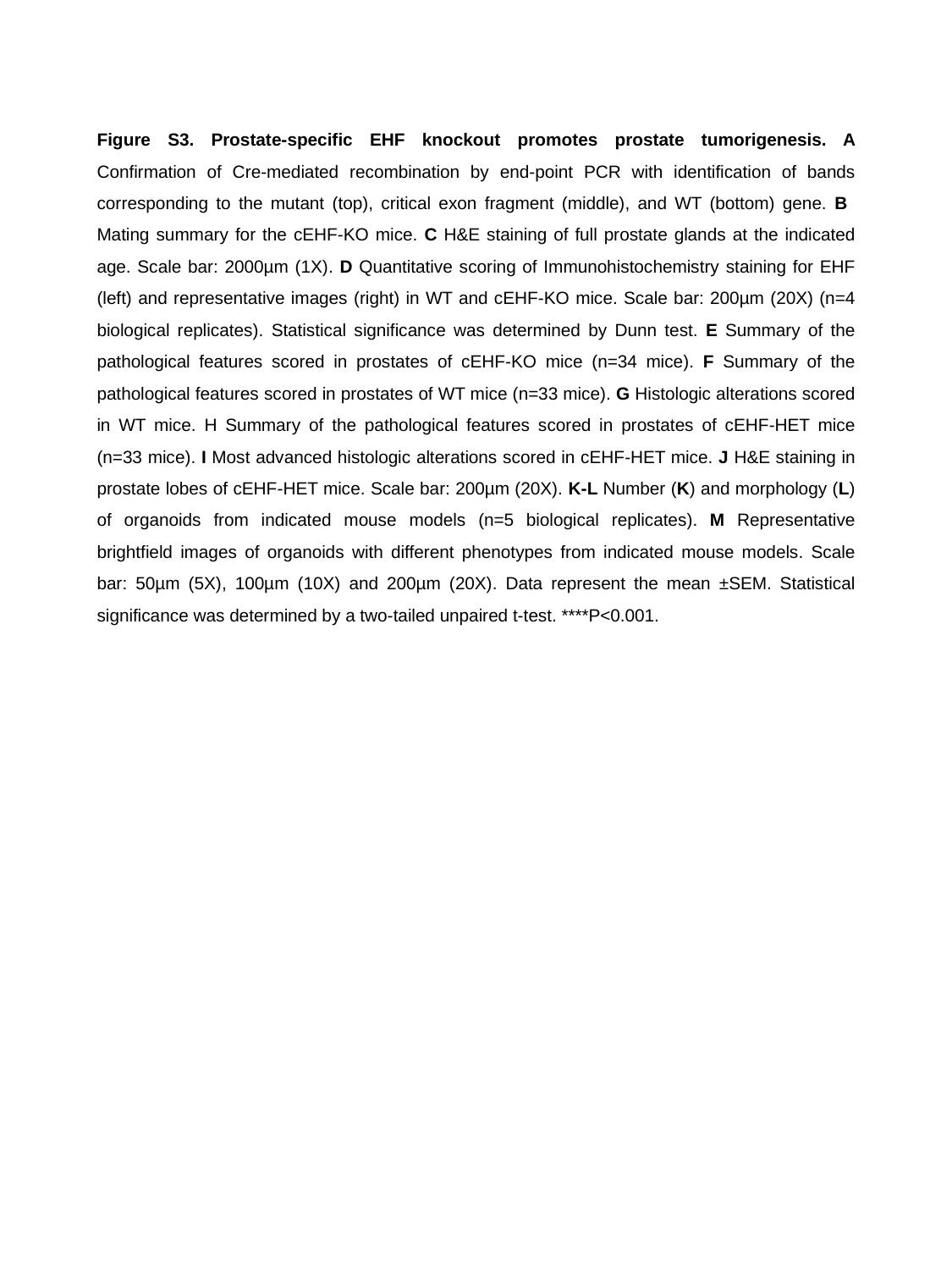

Figure S3. Prostate-specific EHF knockout promotes prostate tumorigenesis. A Confirmation of Cre-mediated recombination by end-point PCR with identification of bands corresponding to the mutant (top), critical exon fragment (middle), and WT (bottom) gene. B Mating summary for the cEHF-KO mice. C H&E staining of full prostate glands at the indicated age. Scale bar: 2000µm (1X). D Quantitative scoring of Immunohistochemistry staining for EHF (left) and representative images (right) in WT and cEHF-KO mice. Scale bar: 200µm (20X) (n=4 biological replicates). Statistical significance was determined by Dunn test. E Summary of the pathological features scored in prostates of cEHF-KO mice (n=34 mice). F Summary of the pathological features scored in prostates of WT mice (n=33 mice). G Histologic alterations scored in WT mice. H Summary of the pathological features scored in prostates of cEHF-HET mice (n=33 mice). I Most advanced histologic alterations scored in cEHF-HET mice. J H&E staining in prostate lobes of cEHF-HET mice. Scale bar: 200µm (20X). K-L Number (K) and morphology (L) of organoids from indicated mouse models (n=5 biological replicates). M Representative brightfield images of organoids with different phenotypes from indicated mouse models. Scale bar: 50µm (5X), 100µm (10X) and 200µm (20X). Data represent the mean ±SEM. Statistical significance was determined by a two-tailed unpaired t-test. ****P<0.001.

### Slide 8
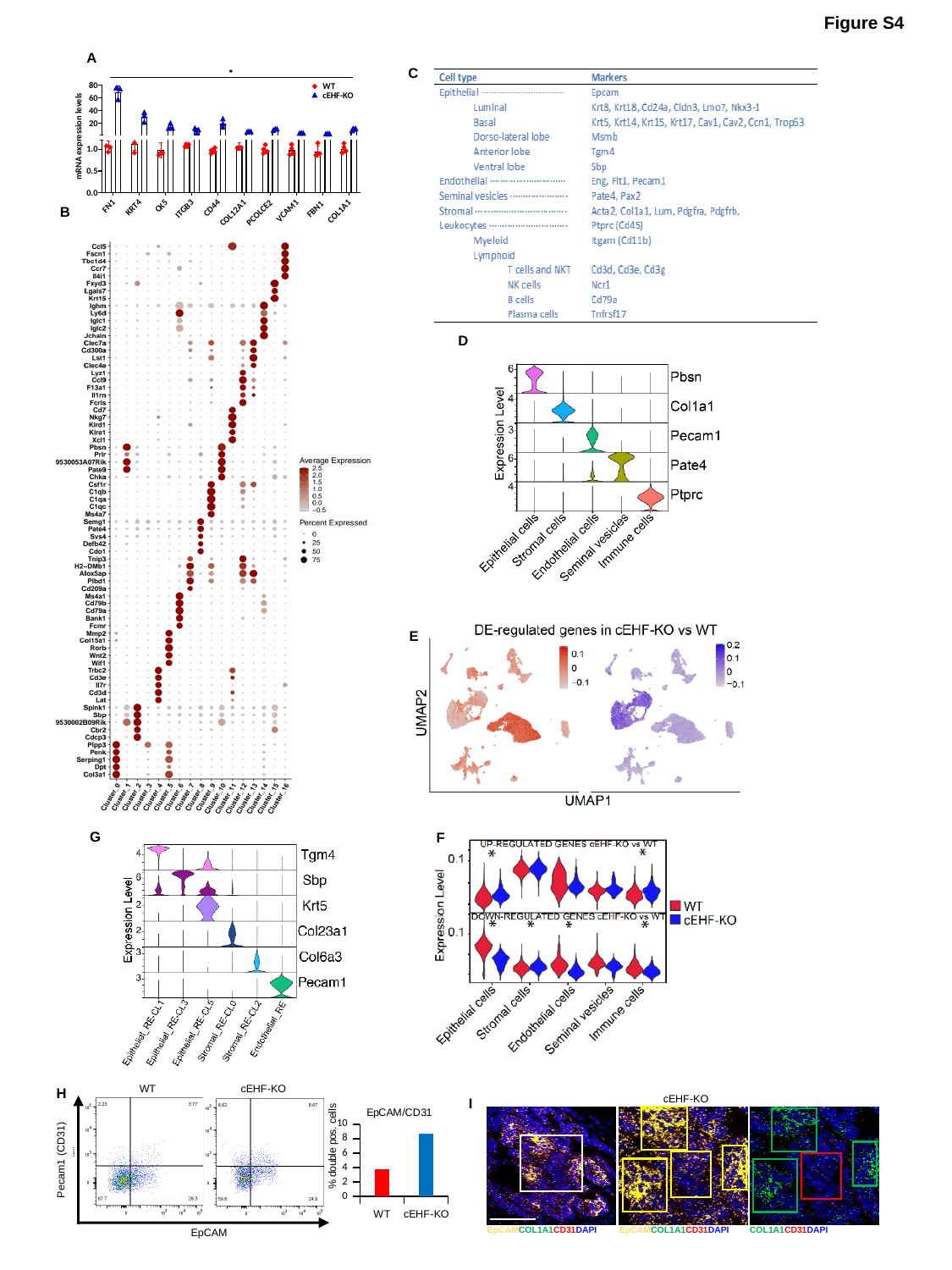

Figure S4
A
C
B
D
E
G
F
WT
Pecam1 (CD31)
EpCAM
cEHF-KO
H
#### Chart: EpCAM/CD31
| Category | |
|---|---|
| WT | 3.77 |
| cEHF-KO | 8.67 |
cEHF-KO
I
EpCAMCOL1A1CD31DAPI
EpCAMCOL1A1CD31DAPI
COL1A1CD31DAPI

### Slide 9
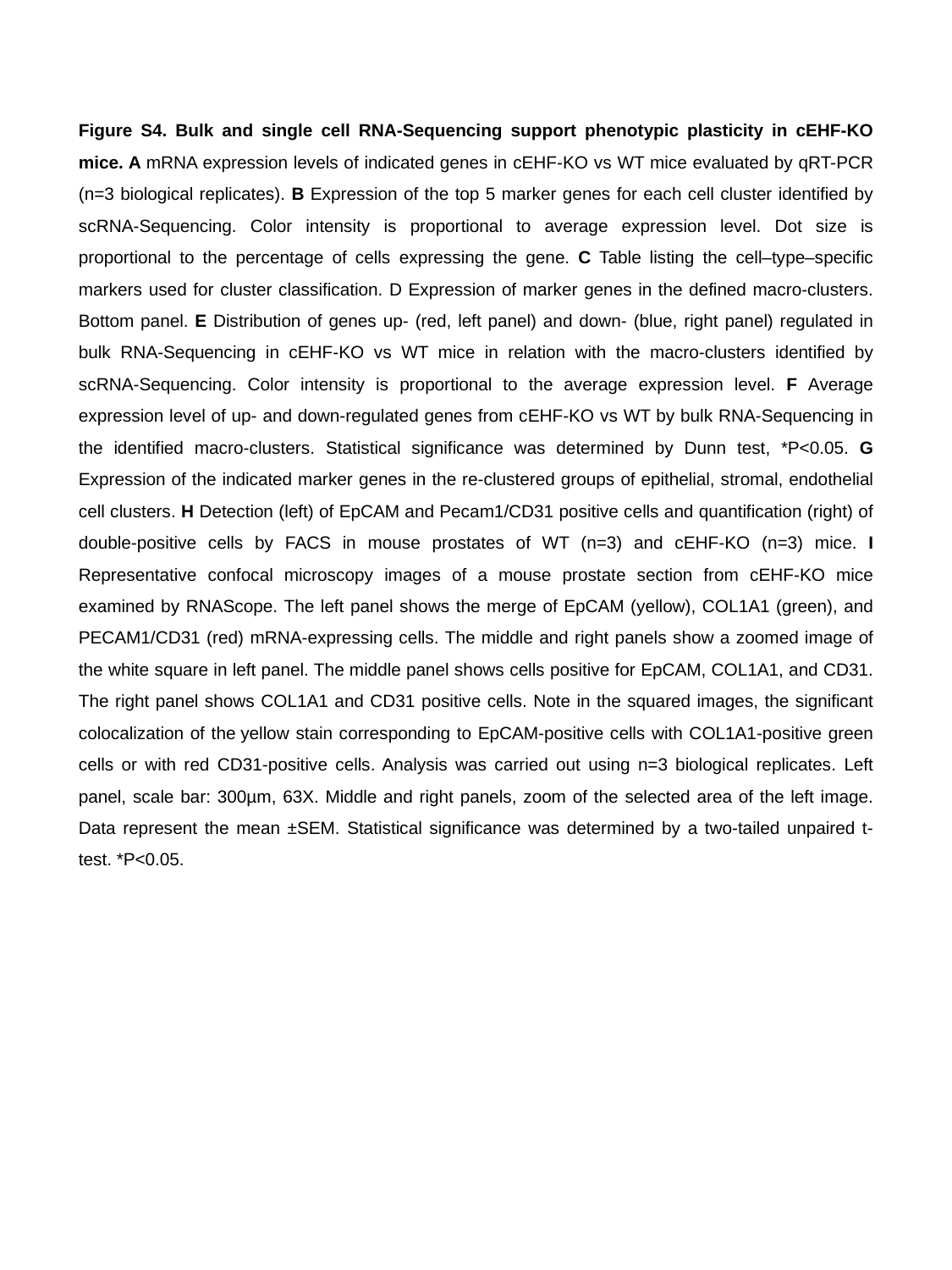

Figure S4. Bulk and single cell RNA-Sequencing support phenotypic plasticity in cEHF-KO mice. A mRNA expression levels of indicated genes in cEHF-KO vs WT mice evaluated by qRT-PCR (n=3 biological replicates). B Expression of the top 5 marker genes for each cell cluster identified by scRNA-Sequencing. Color intensity is proportional to average expression level. Dot size is proportional to the percentage of cells expressing the gene. C Table listing the cell–type–specific markers used for cluster classification. D Expression of marker genes in the defined macro-clusters. Bottom panel. E Distribution of genes up- (red, left panel) and down- (blue, right panel) regulated in bulk RNA-Sequencing in cEHF-KO vs WT mice in relation with the macro-clusters identified by scRNA-Sequencing. Color intensity is proportional to the average expression level. F Average expression level of up- and down-regulated genes from cEHF-KO vs WT by bulk RNA-Sequencing in the identified macro-clusters. Statistical significance was determined by Dunn test, *P<0.05. G Expression of the indicated marker genes in the re-clustered groups of epithelial, stromal, endothelial cell clusters. H Detection (left) of EpCAM and Pecam1/CD31 positive cells and quantification (right) of double-positive cells by FACS in mouse prostates of WT (n=3) and cEHF-KO (n=3) mice. I Representative confocal microscopy images of a mouse prostate section from cEHF-KO mice examined by RNAScope. The left panel shows the merge of EpCAM (yellow), COL1A1 (green), and PECAM1/CD31 (red) mRNA-expressing cells. The middle and right panels show a zoomed image of the white square in left panel. The middle panel shows cells positive for EpCAM, COL1A1, and CD31. The right panel shows COL1A1 and CD31 positive cells. Note in the squared images, the significant colocalization of the yellow stain corresponding to EpCAM-positive cells with COL1A1-positive green cells or with red CD31-positive cells. Analysis was carried out using n=3 biological replicates. Left panel, scale bar: 300µm, 63X. Middle and right panels, zoom of the selected area of the left image. Data represent the mean ±SEM. Statistical significance was determined by a two-tailed unpaired t-test. *P<0.05.

### Slide 10
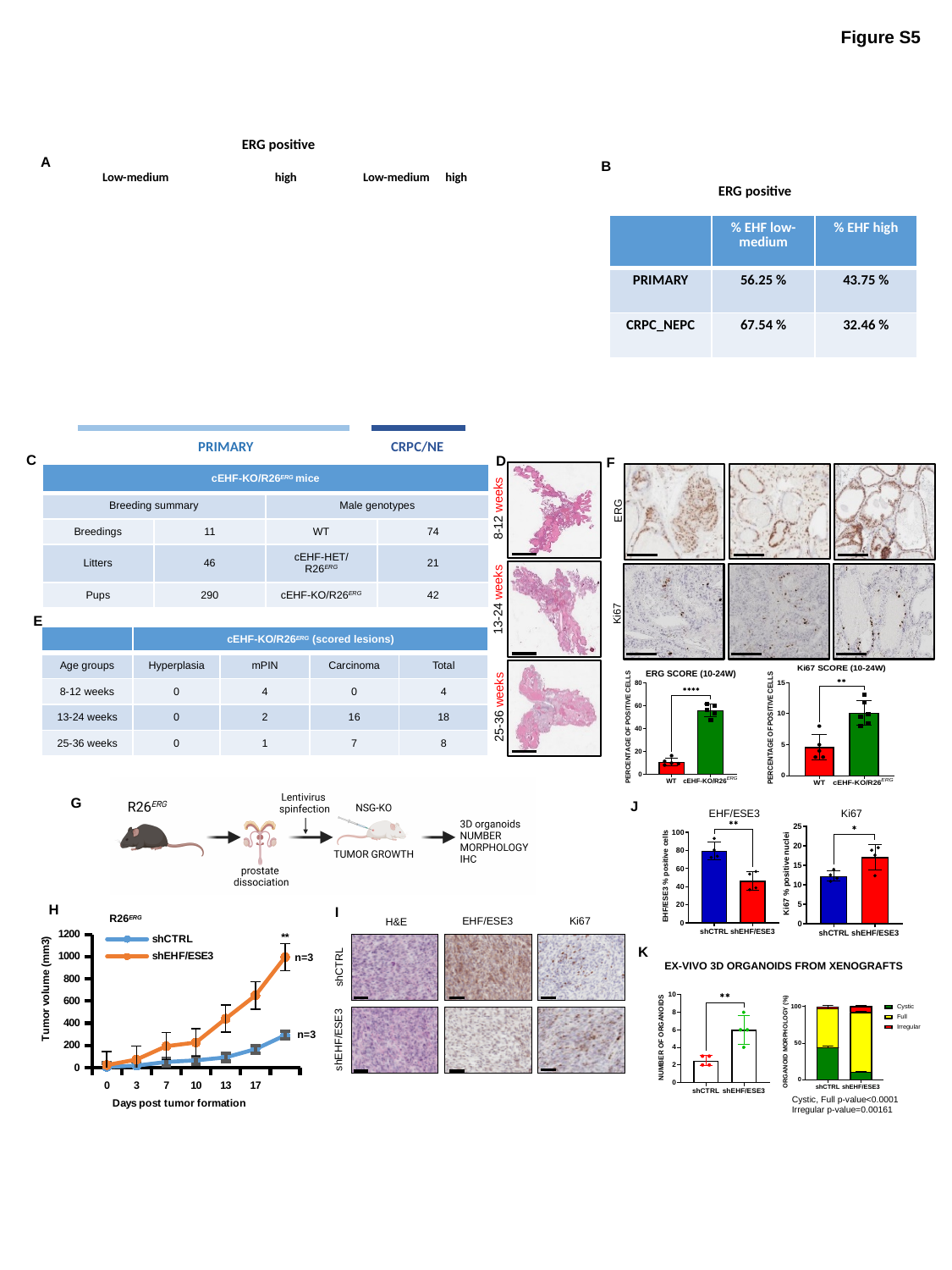

Figure S5
ERG positive
A
 Low-medium
 high
 Low-medium
 high
 PRIMARY
 CRPC/NE
B
ERG positive
| | % EHF low-medium | % EHF high |
| --- | --- | --- |
| PRIMARY | 56.25 % | 43.75 % |
| CRPC\_NEPC | 67.54 % | 32.46 % |
C
D
8-12 weeks
13-24 weeks
25-36 weeks
F
ERG
Ki67
| cEHF-KO/R26^ERG^ mice | cEHF mice | | |
| --- | --- | --- | --- |
| Breeding summary | | Male genotypes | |
| Breedings | 11 | WT | 74 |
| Litters | 46 | cEHF-HET/R26^ERG^ | 21 |
| Pups | 290 | cEHF-KO/R26^ERG^ | 42 |
E
| | cEHF-KO/R26^ERG^ (scored lesions) | | | |
| --- | --- | --- | --- | --- |
| Age groups | Hyperplasia | mPIN | Carcinoma | Total |
| 8-12 weeks | 0 | 4 | 0 | 4 |
| 13-24 weeks | 0 | 2 | 16 | 18 |
| 25-36 weeks | 0 | 1 | 7 | 8 |
G
J
EHF/ESE3
Ki67
H
#### Chart
| Category | shCTRL | shEHF/ESE3 |
|---|---|---|
| 0 | 6.45866 | 22.6954 |
| 3 | 17.34278 | 72.7441 |
| 7 | 50.7676 | 192.92000000000002 |
| 10 | 65.68484000000001 | 226.48366 |
| 13 | 92.05300000000001 | 440.57 |
| 17 | 165.99492 | 650.0 |n=3
n=3
R26^ERG^
I
Ki67
EHF/ESE3
H&E
**
shCTRL
shEHF/ESE3
K
EX-VIVO 3D ORGANOIDS FROM XENOGRAFTS
Cystic, Full p-value<0.0001
Irregular p-value=0.00161

### Slide 11
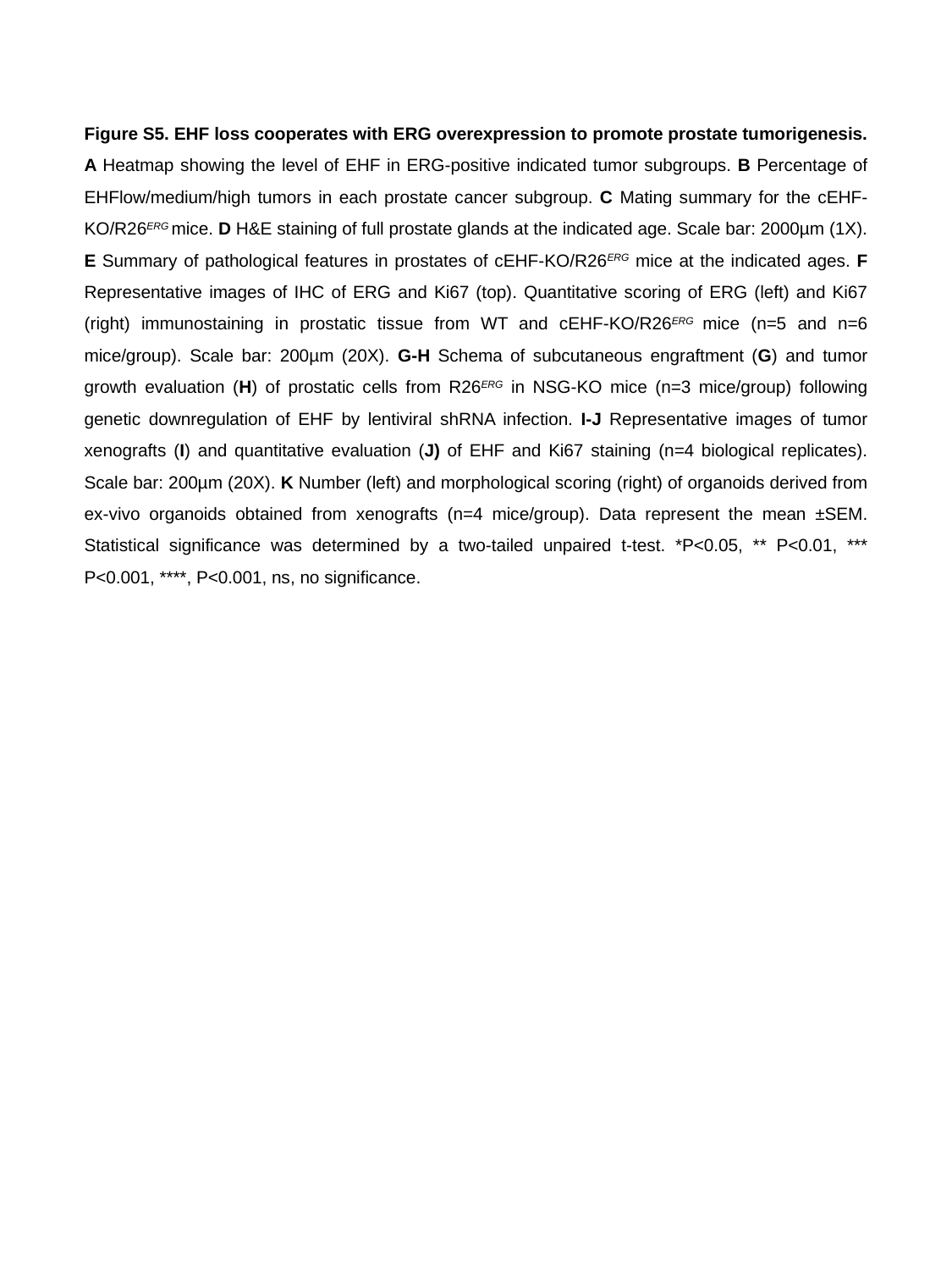

Figure S5. EHF loss cooperates with ERG overexpression to promote prostate tumorigenesis. A Heatmap showing the level of EHF in ERG-positive indicated tumor subgroups. B Percentage of EHFlow/medium/high tumors in each prostate cancer subgroup. C Mating summary for the cEHF-KO/R26^ERG^ mice. D H&E staining of full prostate glands at the indicated age. Scale bar: 2000µm (1X). E Summary of pathological features in prostates of cEHF-KO/R26^ERG^ mice at the indicated ages. F Representative images of IHC of ERG and Ki67 (top). Quantitative scoring of ERG (left) and Ki67 (right) immunostaining in prostatic tissue from WT and cEHF-KO/R26^ERG^ mice (n=5 and n=6 mice/group). Scale bar: 200µm (20X). G-H Schema of subcutaneous engraftment (G) and tumor growth evaluation (H) of prostatic cells from R26^ERG^ in NSG-KO mice (n=3 mice/group) following genetic downregulation of EHF by lentiviral shRNA infection. I-J Representative images of tumor xenografts (I) and quantitative evaluation (J) of EHF and Ki67 staining (n=4 biological replicates). Scale bar: 200µm (20X). K Number (left) and morphological scoring (right) of organoids derived from ex-vivo organoids obtained from xenografts (n=4 mice/group). Data represent the mean ±SEM. Statistical significance was determined by a two-tailed unpaired t-test. *P<0.05, ** P<0.01, *** P<0.001, ****, P<0.001, ns, no significance.

### Slide 12
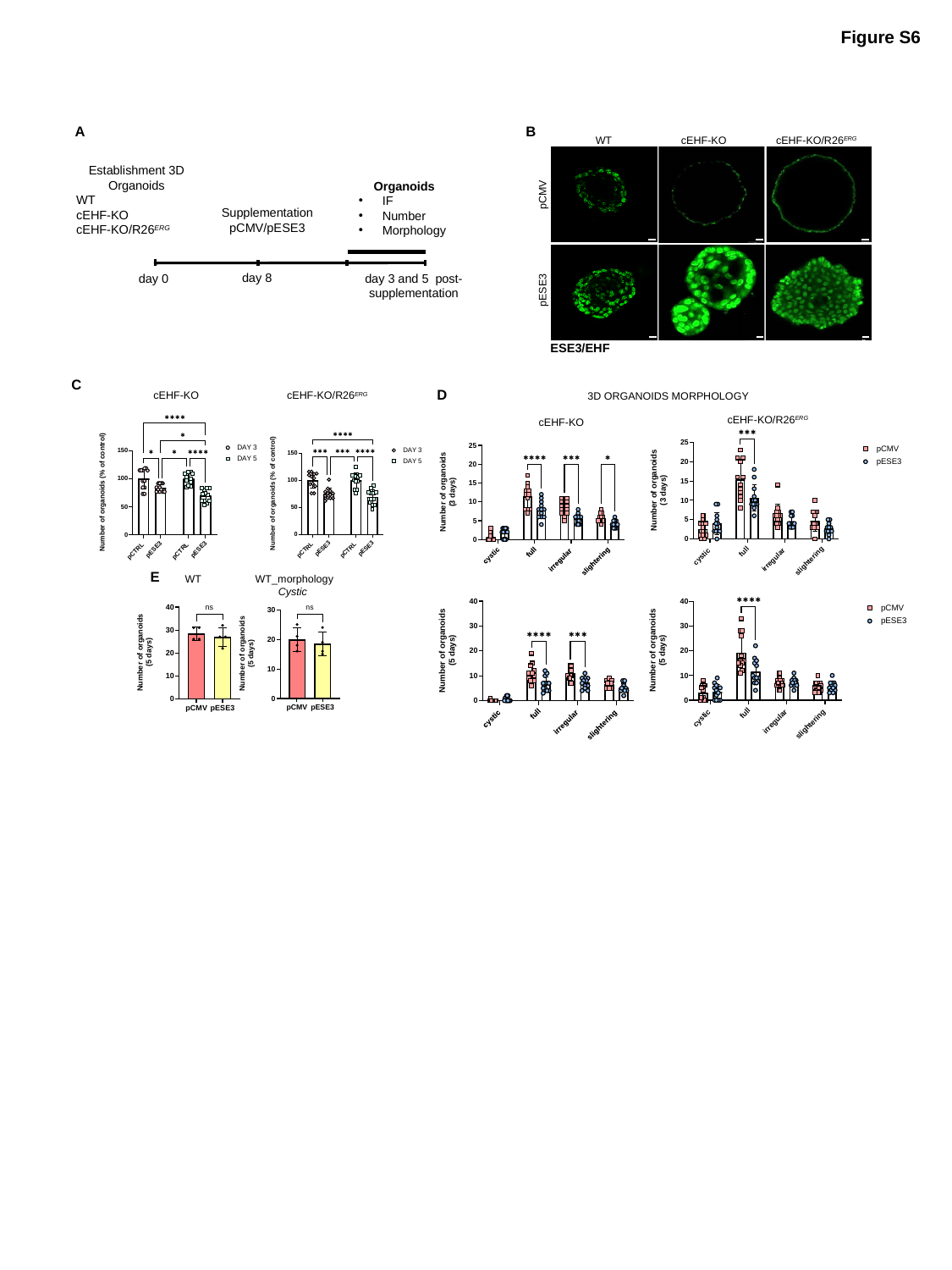

Figure S6
A
B
WT
cEHF-KO
cEHF-KO/R26^ERG^
Establishment 3D Organoids
WT
cEHF-KO
cEHF-KO/R26^ERG^
Organoids
IF
Number
Morphology
Supplementation pCMV/pESE3
day 8
day 3 and 5 post-supplementation
day 0
pCMV
pESE3
ESE3/EHF
C
D
cEHF-KO
cEHF-KO/R26^ERG^
3D ORGANOIDS MORPHOLOGY
cEHF-KO/R26^ERG^
cEHF-KO
E
WT
WT_morphology
Cystic

### Slide 13
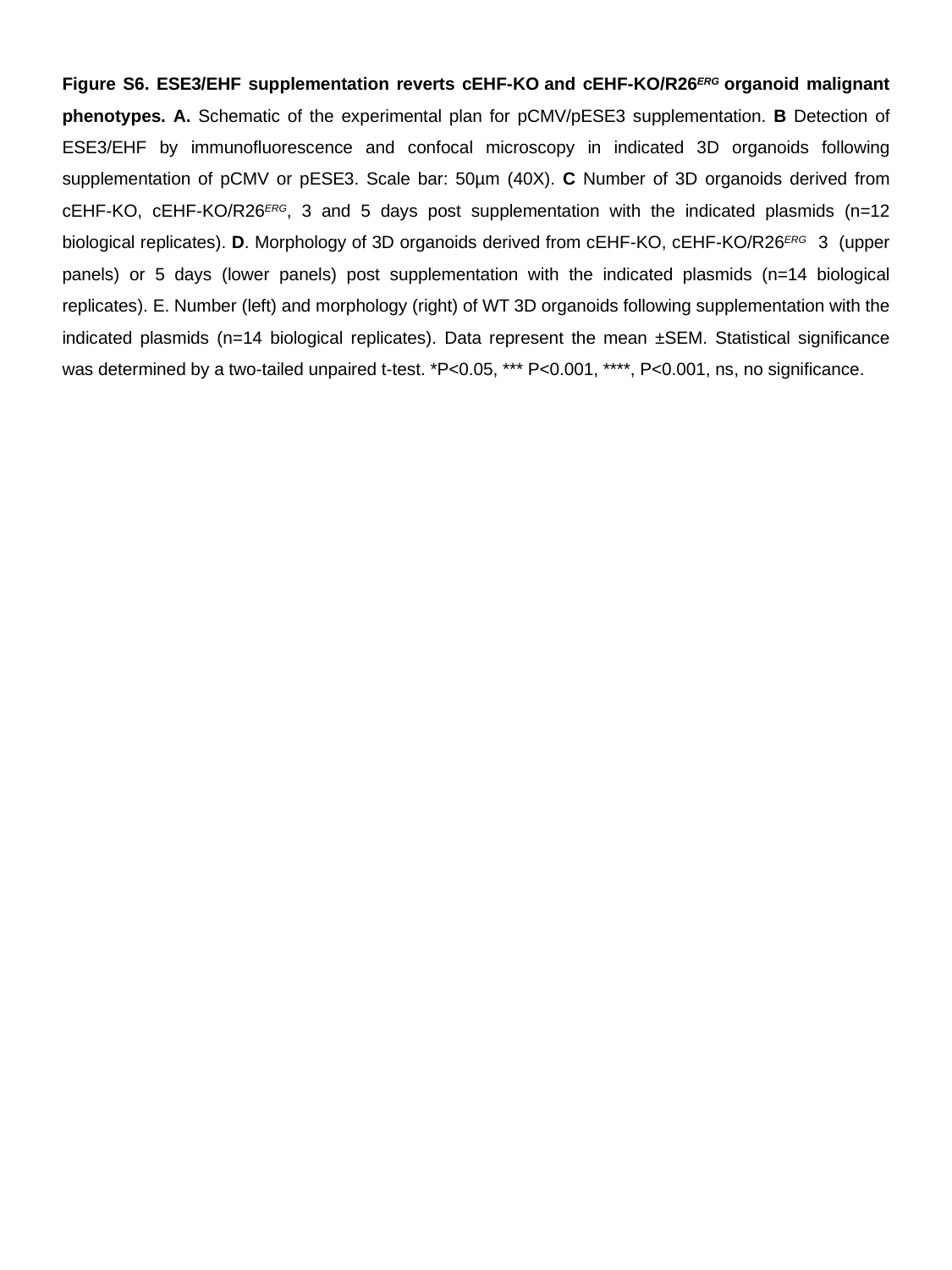

Figure S6. ESE3/EHF supplementation reverts cEHF-KO and cEHF-KO/R26^ERG^ organoid malignant phenotypes. A. Schematic of the experimental plan for pCMV/pESE3 supplementation. B Detection of ESE3/EHF by immunofluorescence and confocal microscopy in indicated 3D organoids following supplementation of pCMV or pESE3. Scale bar: 50µm (40X). C Number of 3D organoids derived from cEHF-KO, cEHF-KO/R26^ERG^, 3 and 5 days post supplementation with the indicated plasmids (n=12 biological replicates). D. Morphology of 3D organoids derived from cEHF-KO, cEHF-KO/R26^ERG^ 3 (upper panels) or 5 days (lower panels) post supplementation with the indicated plasmids (n=14 biological replicates). E. Number (left) and morphology (right) of WT 3D organoids following supplementation with the indicated plasmids (n=14 biological replicates). Data represent the mean ±SEM. Statistical significance was determined by a two-tailed unpaired t-test. *P<0.05, *** P<0.001, ****, P<0.001, ns, no significance.

### Slide 14
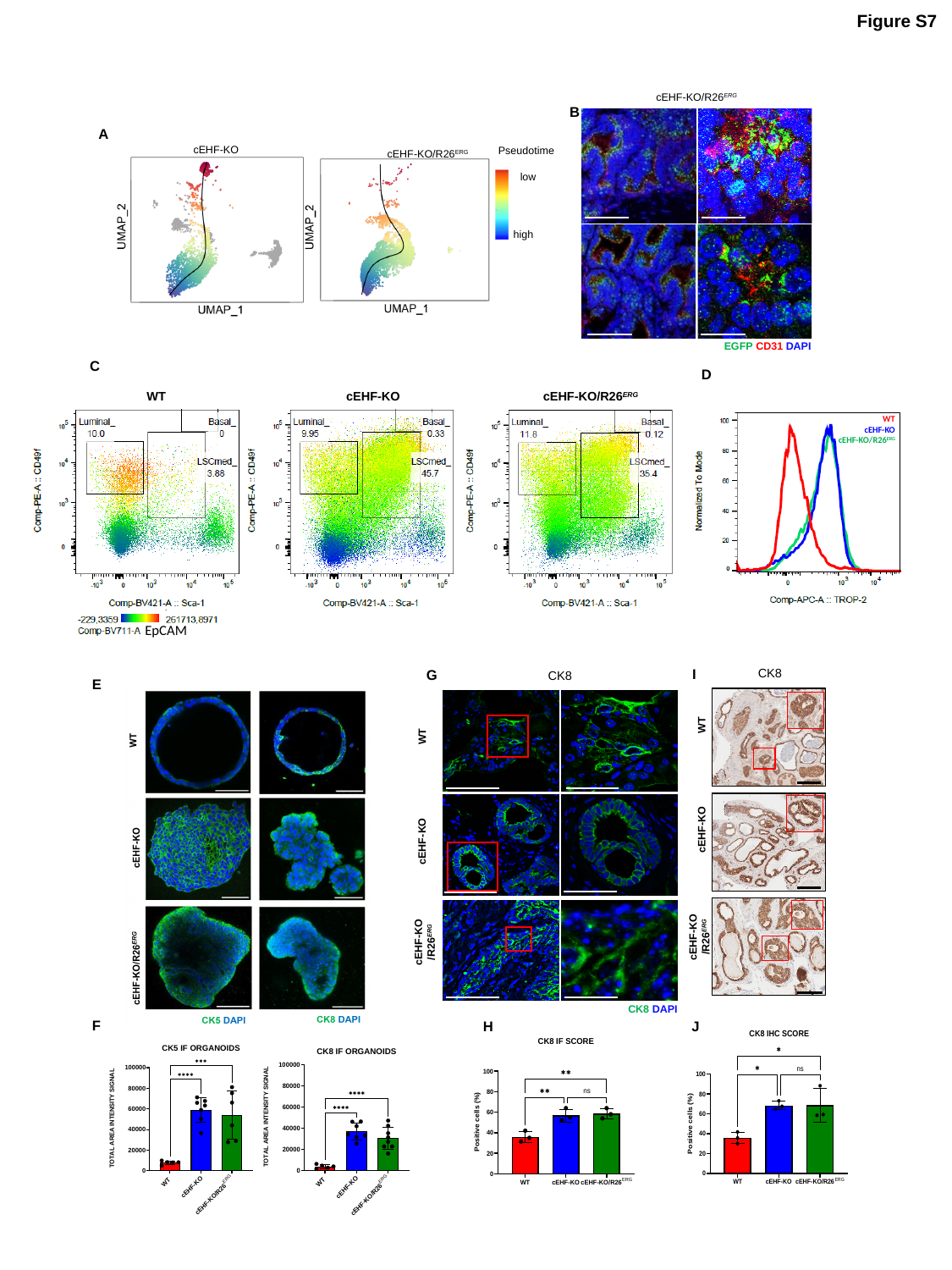

Figure S7
cEHF-KO/R26^ERG^
EGFP CD31 DAPI
B
A
Pseudotime
low
high
cEHF-KO
cEHF-KO/R26^ERG^
C
D
WT
cEHF-KO
cEHF-KO/R26^ERG^
EpCAM
WT
cEHF-KO
cEHF-KO/R26^ERG^
I
CK8
WT
cEHF-KO
cEHF-KO
/R26^ERG^
J
G
CK8
WT
cEHF-KO
cEHF-KO
/R26^ERG^
CK8 DAPI
H
E
F

### Slide 15
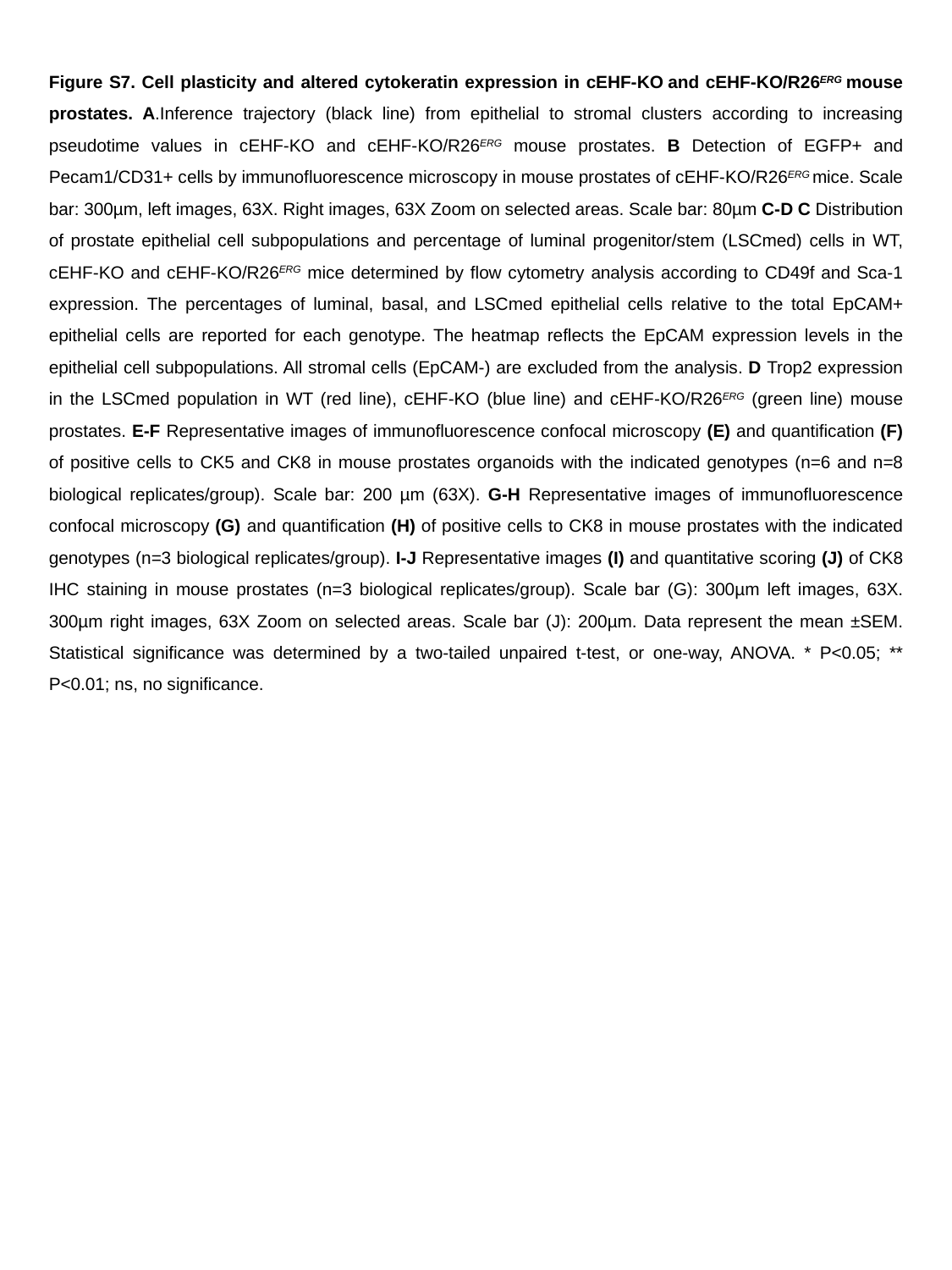

Figure S7. Cell plasticity and altered cytokeratin expression in cEHF-KO and cEHF-KO/R26^ERG^ mouse prostates. A.Inference trajectory (black line) from epithelial to stromal clusters according to increasing pseudotime values in cEHF-KO and cEHF-KO/R26^ERG^ mouse prostates. B Detection of EGFP+ and Pecam1/CD31+ cells by immunofluorescence microscopy in mouse prostates of cEHF-KO/R26^ERG^ mice. Scale bar: 300µm, left images, 63X. Right images, 63X Zoom on selected areas. Scale bar: 80µm C-D C Distribution of prostate epithelial cell subpopulations and percentage of luminal progenitor/stem (LSCmed) cells in WT, cEHF-KO and cEHF-KO/R26^ERG^ mice determined by flow cytometry analysis according to CD49f and Sca-1 expression. The percentages of luminal, basal, and LSCmed epithelial cells relative to the total EpCAM+ epithelial cells are reported for each genotype. The heatmap reflects the EpCAM expression levels in the epithelial cell subpopulations. All stromal cells (EpCAM-) are excluded from the analysis. D Trop2 expression in the LSCmed population in WT (red line), cEHF-KO (blue line) and cEHF-KO/R26^ERG^ (green line) mouse prostates. E-F Representative images of immunofluorescence confocal microscopy (E) and quantification (F) of positive cells to CK5 and CK8 in mouse prostates organoids with the indicated genotypes (n=6 and n=8 biological replicates/group). Scale bar: 200 µm (63X). G-H Representative images of immunofluorescence confocal microscopy (G) and quantification (H) of positive cells to CK8 in mouse prostates with the indicated genotypes (n=3 biological replicates/group). I-J Representative images (I) and quantitative scoring (J) of CK8 IHC staining in mouse prostates (n=3 biological replicates/group). Scale bar (G): 300µm left images, 63X. 300µm right images, 63X Zoom on selected areas. Scale bar (J): 200µm. Data represent the mean ±SEM. Statistical significance was determined by a two-tailed unpaired t-test, or one-way, ANOVA. * P<0.05; ** P<0.01; ns, no significance.

### Slide 16
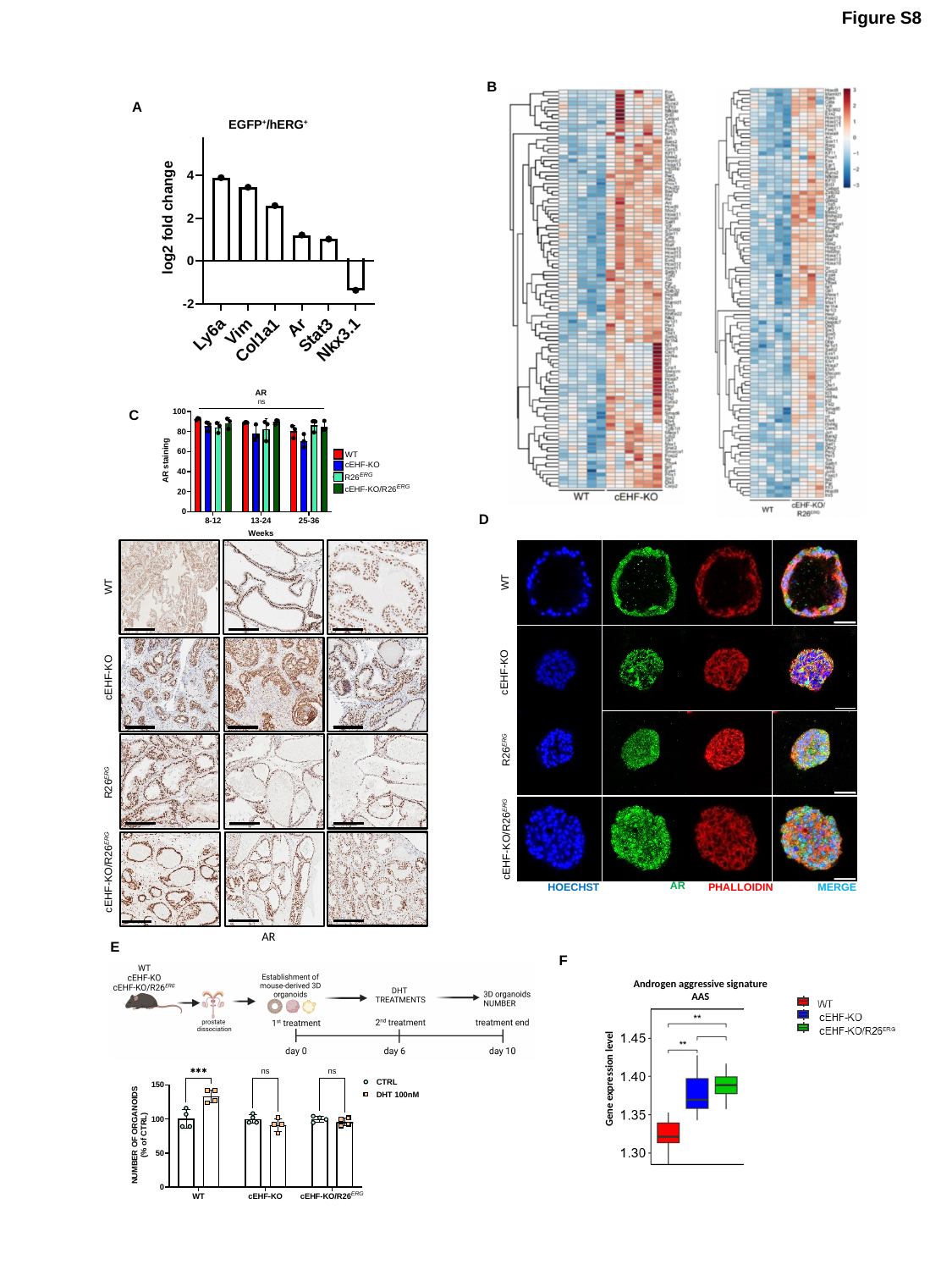

Figure S8
B
A
EGFP^+^/hERG^+^
C
D
WT
 cEHF-KO
 R26^ERG^
cEHF-KO/R26^ERG^
AR
HOECHST
PHALLOIDIN
MERGE
WT
cEHF-KO
R26^ERG^
cEHF-KO/R26^ERG^
AR
E
F
Androgen aggressive signature
AAS
Gene expression level

### Slide 17
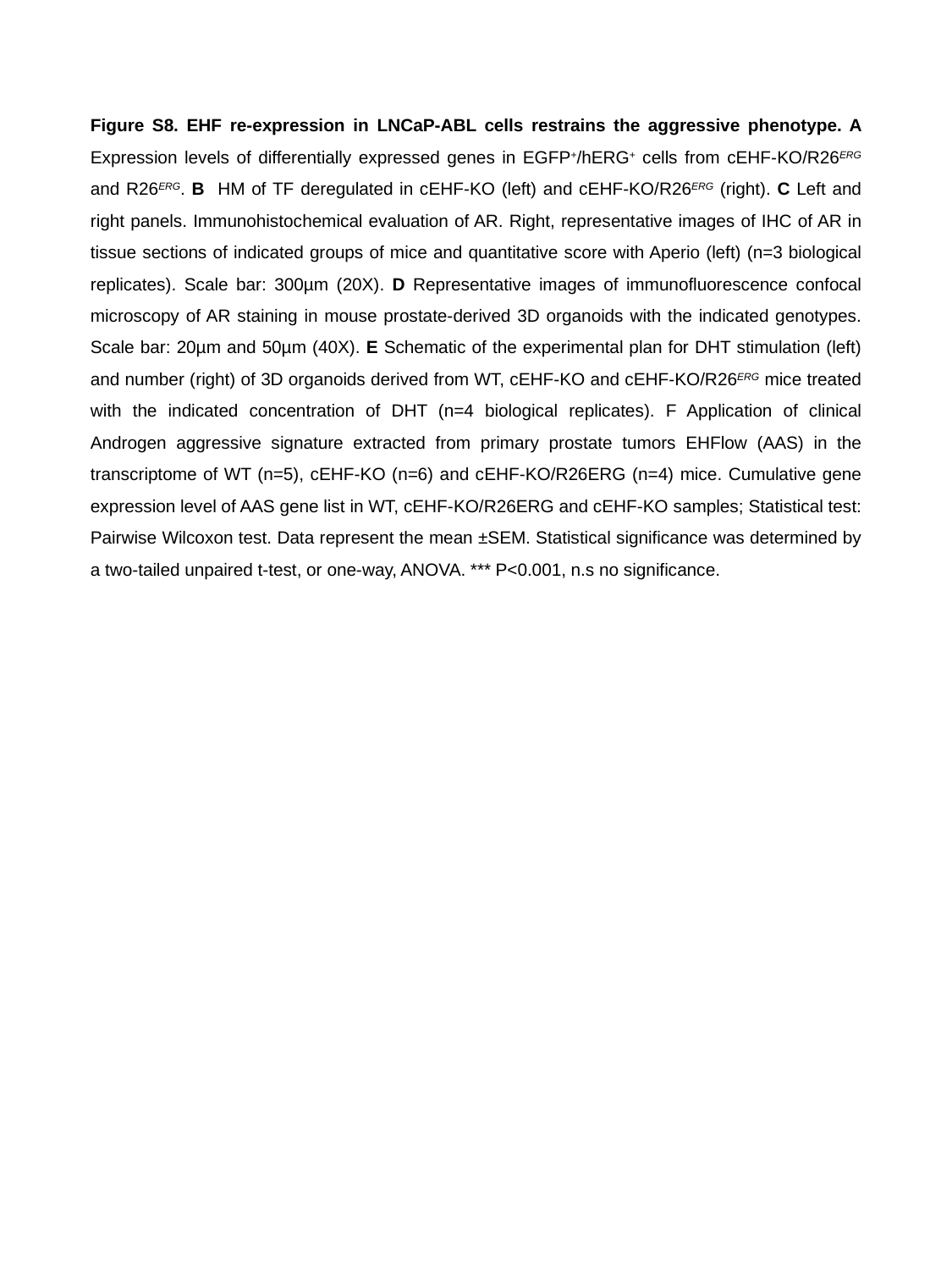

Figure S8. EHF re-expression in LNCaP-ABL cells restrains the aggressive phenotype. A Expression levels of differentially expressed genes in EGFP^+^/hERG^+^ cells from cEHF-KO/R26^ERG^ and R26^ERG^. B HM of TF deregulated in cEHF-KO (left) and cEHF-KO/R26^ERG^ (right). C Left and right panels. Immunohistochemical evaluation of AR. Right, representative images of IHC of AR in tissue sections of indicated groups of mice and quantitative score with Aperio (left) (n=3 biological replicates). Scale bar: 300µm (20X). D Representative images of immunofluorescence confocal microscopy of AR staining in mouse prostate-derived 3D organoids with the indicated genotypes. Scale bar: 20µm and 50µm (40X). E Schematic of the experimental plan for DHT stimulation (left) and number (right) of 3D organoids derived from WT, cEHF-KO and cEHF-KO/R26^ERG^ mice treated with the indicated concentration of DHT (n=4 biological replicates). F Application of clinical Androgen aggressive signature extracted from primary prostate tumors EHFlow (AAS) in the transcriptome of WT (n=5), cEHF-KO (n=6) and cEHF-KO/R26ERG (n=4) mice. Cumulative gene expression level of AAS gene list in WT, cEHF-KO/R26ERG and cEHF-KO samples; Statistical test: Pairwise Wilcoxon test. Data represent the mean ±SEM. Statistical significance was determined by a two-tailed unpaired t-test, or one-way, ANOVA. *** P<0.001, n.s no significance.

### Slide 18
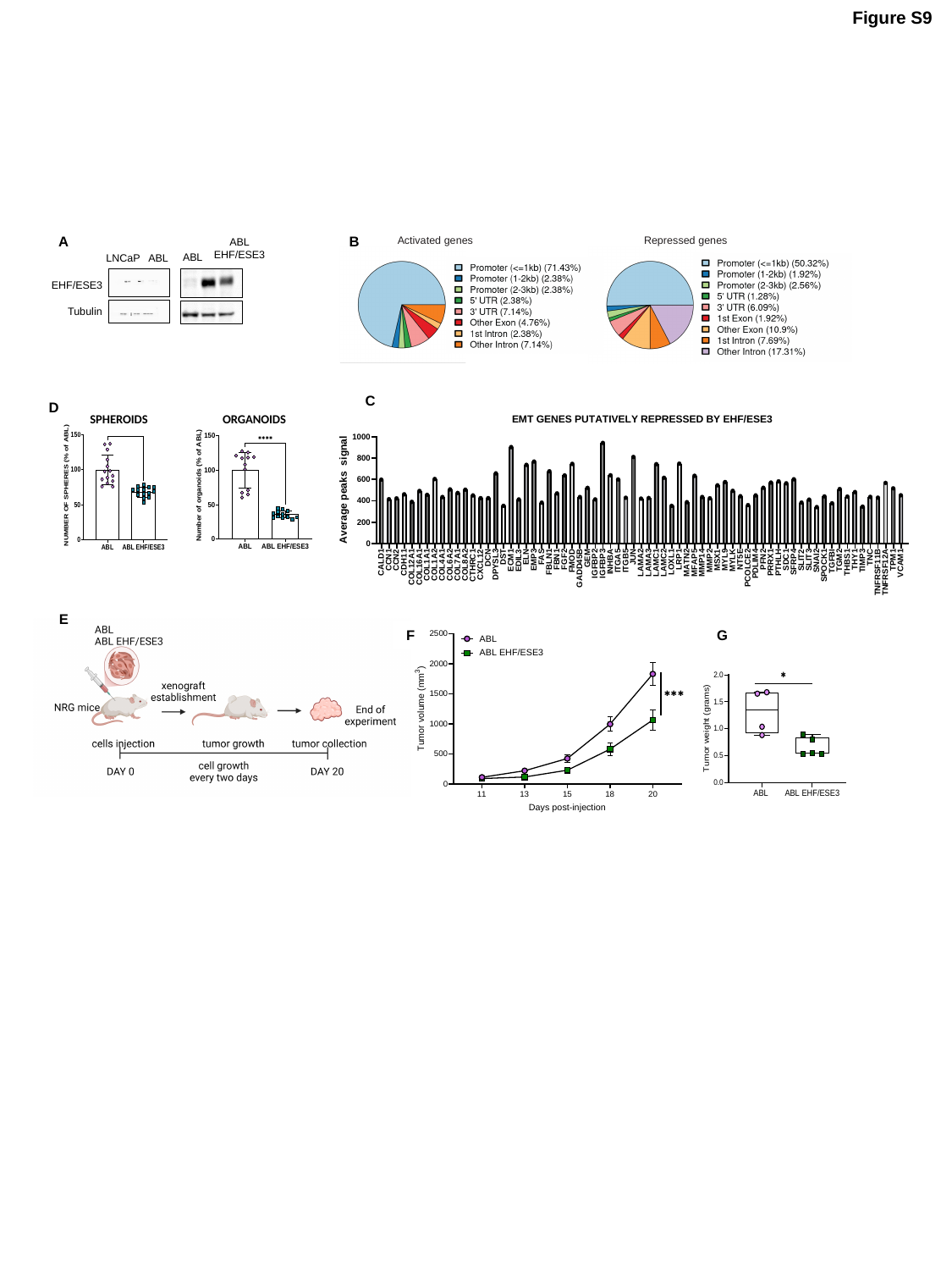

Figure S9
A
B
Repressed genes
Activated genes
ABL EHF/ESE3
ABL
LNCaP
 ABL
EHF/ESE3
Tubulin
C
D
SPHEROIDS
ORGANOIDS
E
F
G

### Slide 19
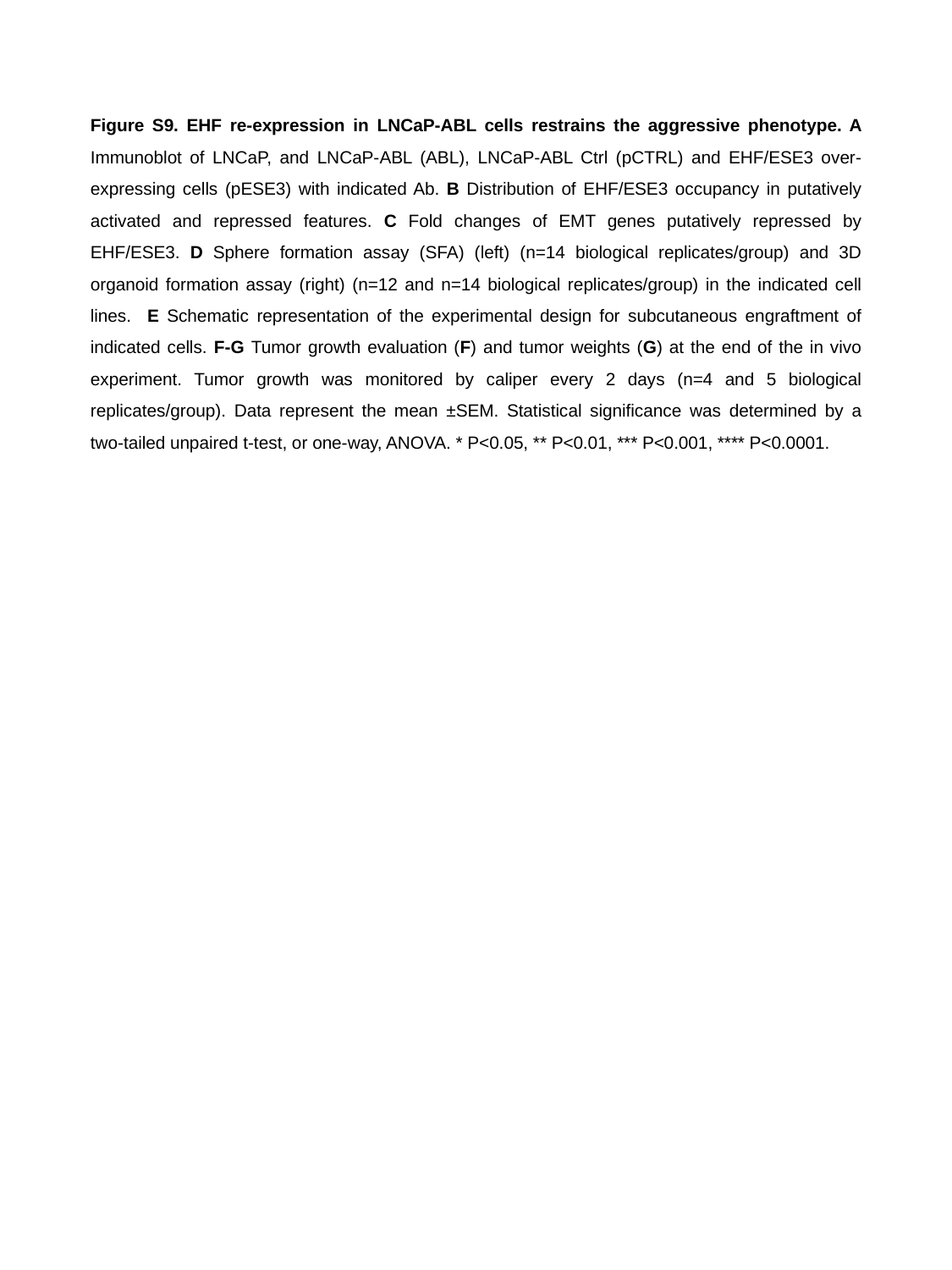

Figure S9. EHF re-expression in LNCaP-ABL cells restrains the aggressive phenotype. A Immunoblot of LNCaP, and LNCaP-ABL (ABL), LNCaP-ABL Ctrl (pCTRL) and EHF/ESE3 over-expressing cells (pESE3) with indicated Ab. B Distribution of EHF/ESE3 occupancy in putatively activated and repressed features. C Fold changes of EMT genes putatively repressed by EHF/ESE3. D Sphere formation assay (SFA) (left) (n=14 biological replicates/group) and 3D organoid formation assay (right) (n=12 and n=14 biological replicates/group) in the indicated cell lines. E Schematic representation of the experimental design for subcutaneous engraftment of indicated cells. F-G Tumor growth evaluation (F) and tumor weights (G) at the end of the in vivo experiment. Tumor growth was monitored by caliper every 2 days (n=4 and 5 biological replicates/group). Data represent the mean ±SEM. Statistical significance was determined by a two-tailed unpaired t-test, or one-way, ANOVA. * P<0.05, ** P<0.01, *** P<0.001, **** P<0.0001.

### Slide 20
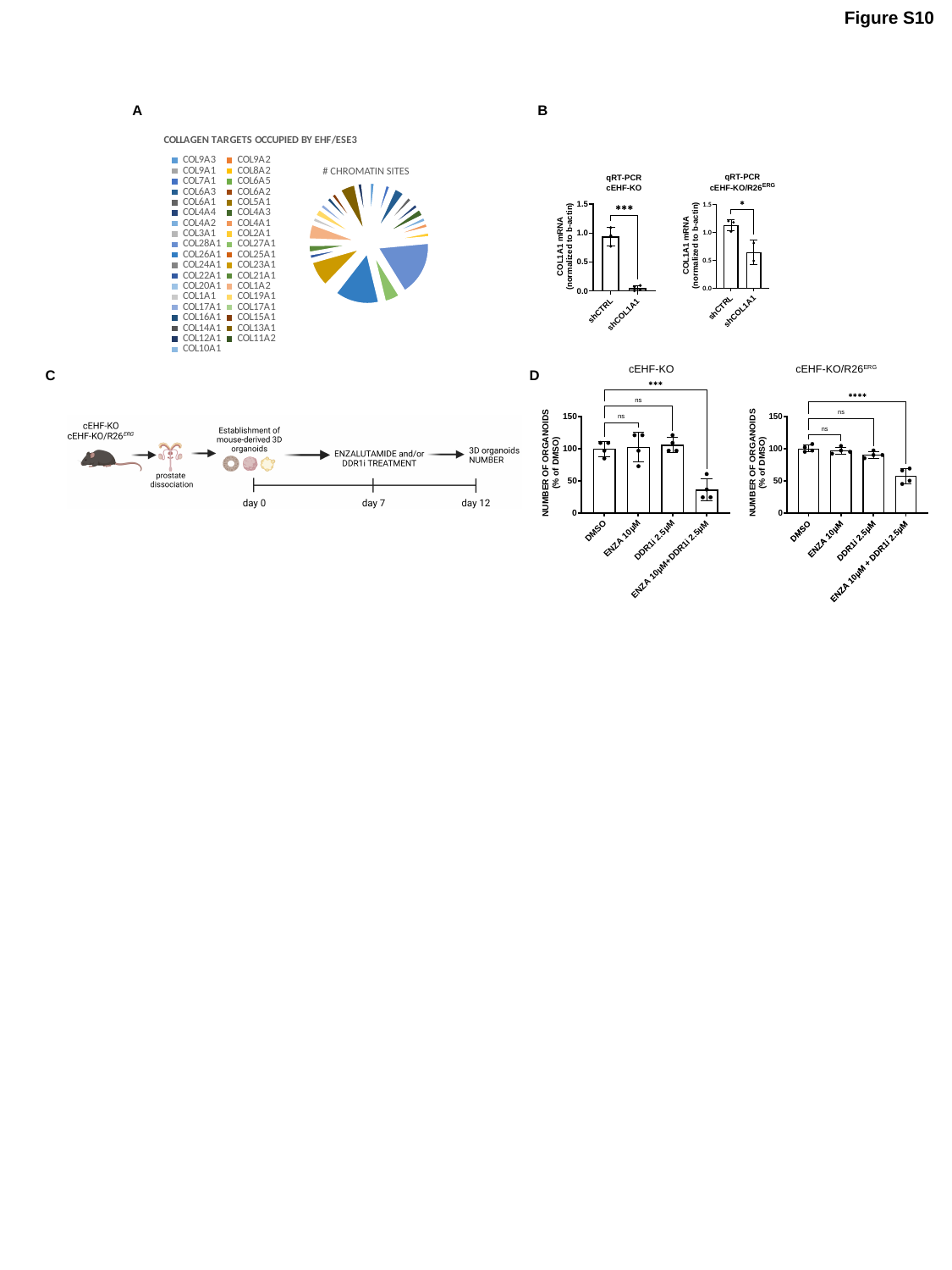

Figure S10
A
B
#### Chart: COLLAGEN TARGETS OCCUPIED BY EHF/ESE3
| Category | # ESE3 OCCUPIED SITES |
|---|---|
| COL9A3 | 2.0 |
| COL9A2 | 1.0 |
| COL9A1 | 1.0 |
| COL8A2 | 1.0 |
| COL7A1 | 2.0 |
| COL6A5 | 1.0 |
| COL6A3 | 4.0 |
| COL6A2 | 1.0 |
| COL6A1 | 2.0 |
| COL5A1 | 1.0 |
| COL4A4 | 2.0 |
| COL4A3 | 3.0 |
| COL4A2 | 2.0 |
| COL4A1 | 2.0 |
| COL3A1 | 1.0 |
| COL2A1 | 2.0 |
| COL28A1 | 21.0 |
| COL27A1 | 6.0 |
| COL26A1 | 17.0 |
| COL25A1 | 1.0 |
| COL24A1 | 1.0 |
| COL23A1 | 10.0 |
| COL22A1 | 2.0 |
| COL21A1 | 3.0 |
| COL20A1 | 1.0 |
| COL1A2 | 6.0 |
| COL1A1 | 2.0 |
| COL19A1 | 3.0 |
| COL17A1 | 2.0 |
| COL17A1 | 1.0 |
| COL16A1 | 2.0 |
| COL15A1 | 2.0 |
| COL14A1 | 1.0 |
| COL13A1 | 6.0 |
| COL12A1 | 2.0 |
| COL11A2 | 1.0 |
| COL10A1 | 1.0 |# CHROMATIN SITES
cEHF-KO/R26^ERG^
cEHF-KO
C
D

### Slide 21
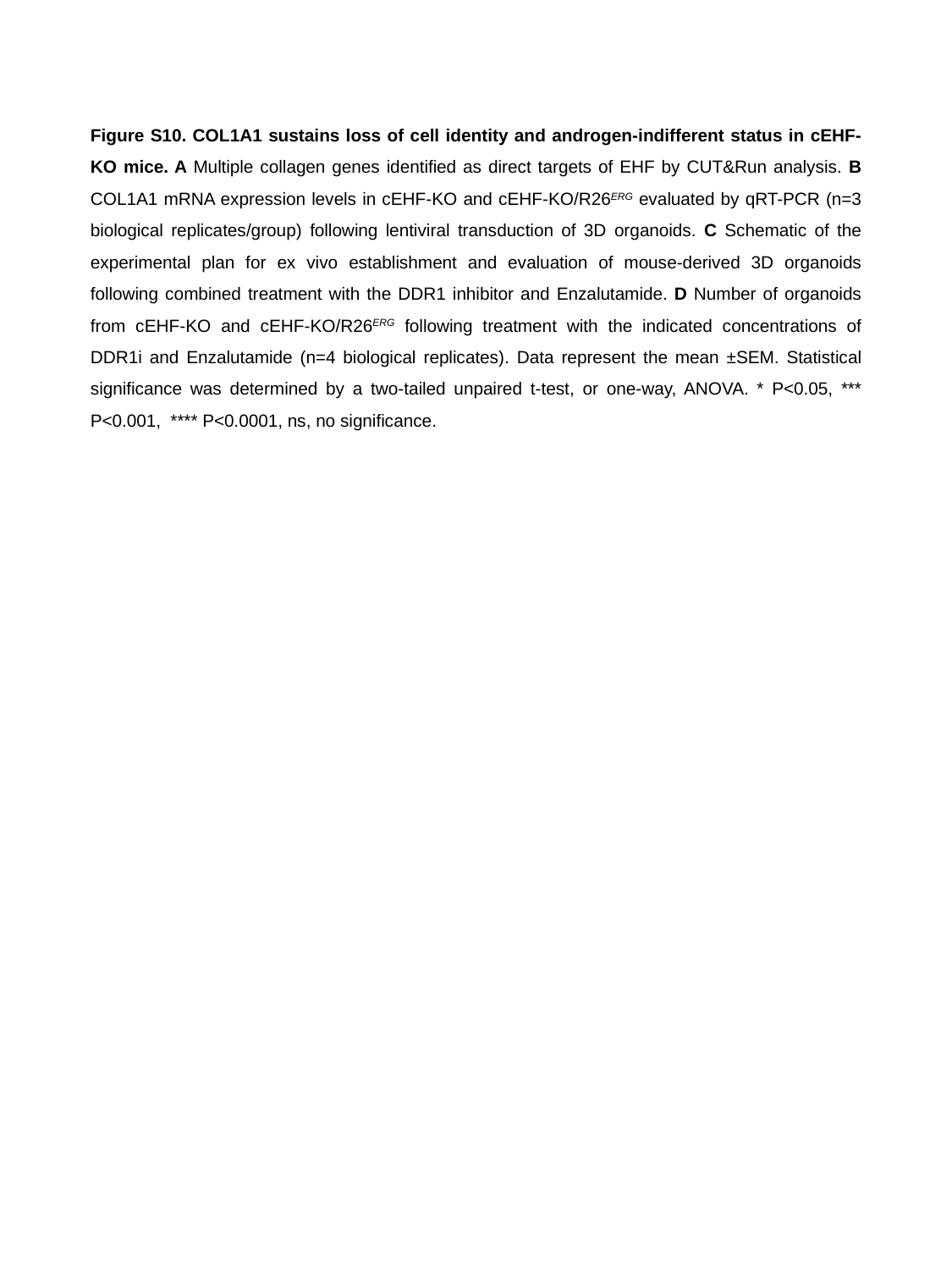

Figure S10. COL1A1 sustains loss of cell identity and androgen-indifferent status in cEHF-KO mice. A Multiple collagen genes identified as direct targets of EHF by CUT&Run analysis. B COL1A1 mRNA expression levels in cEHF-KO and cEHF-KO/R26^ERG^ evaluated by qRT-PCR (n=3 biological replicates/group) following lentiviral transduction of 3D organoids. C Schematic of the experimental plan for ex vivo establishment and evaluation of mouse-derived 3D organoids following combined treatment with the DDR1 inhibitor and Enzalutamide. D Number of organoids from cEHF-KO and cEHF-KO/R26^ERG^ following treatment with the indicated concentrations of DDR1i and Enzalutamide (n=4 biological replicates). Data represent the mean ±SEM. Statistical significance was determined by a two-tailed unpaired t-test, or one-way, ANOVA. * P<0.05, *** P<0.001, **** P<0.0001, ns, no significance.

### Slide 22
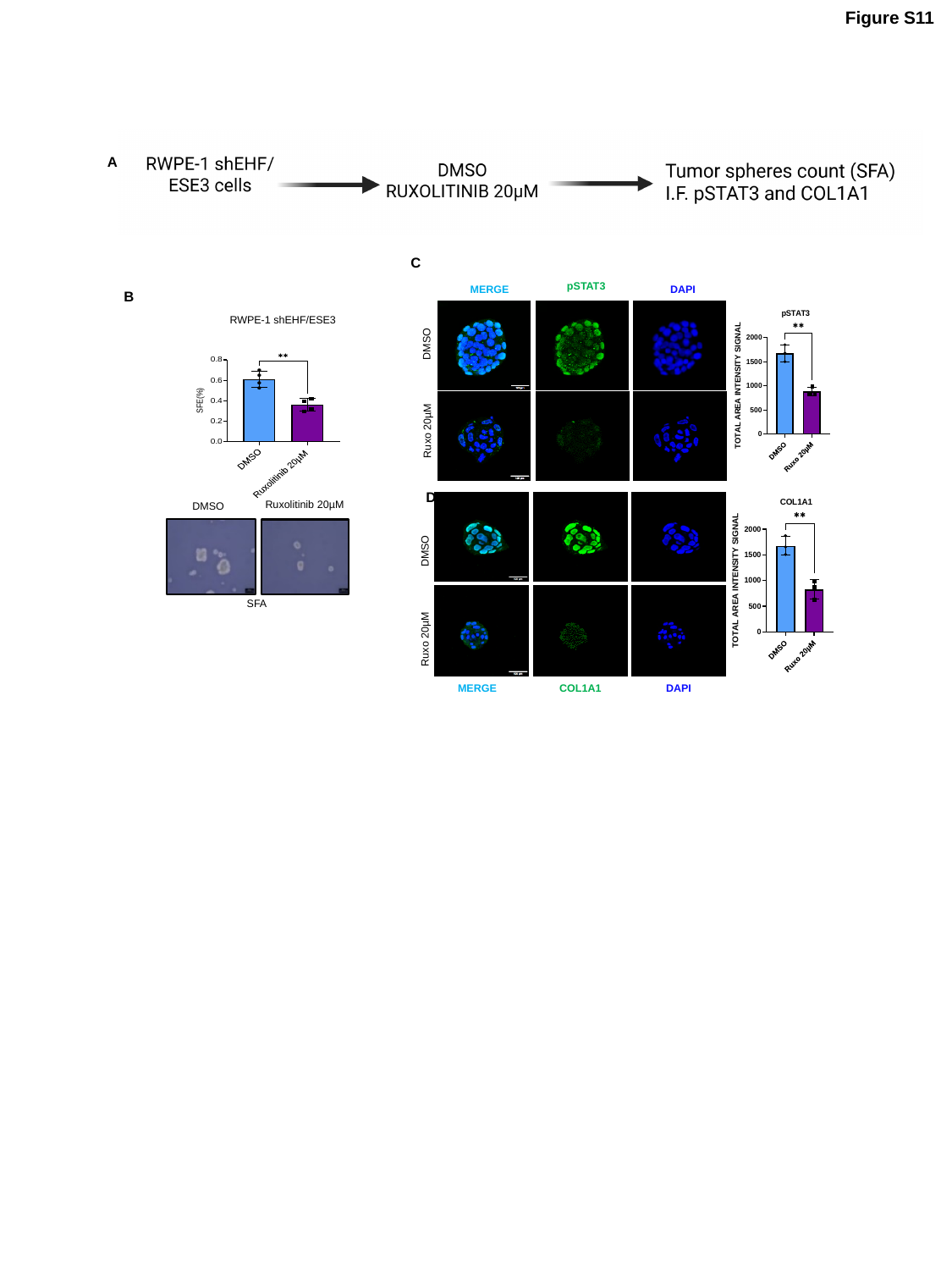

Figure S11
A
C
pSTAT3
DAPI
MERGE
DMSO
Ruxo 20µM
B
RWPE-1 shEHF/ESE3
Ruxolitinib 20µM
DMSO
SFA
D
MERGE
COL1A1
DAPI
DMSO
Ruxo 20µM

### Slide 23
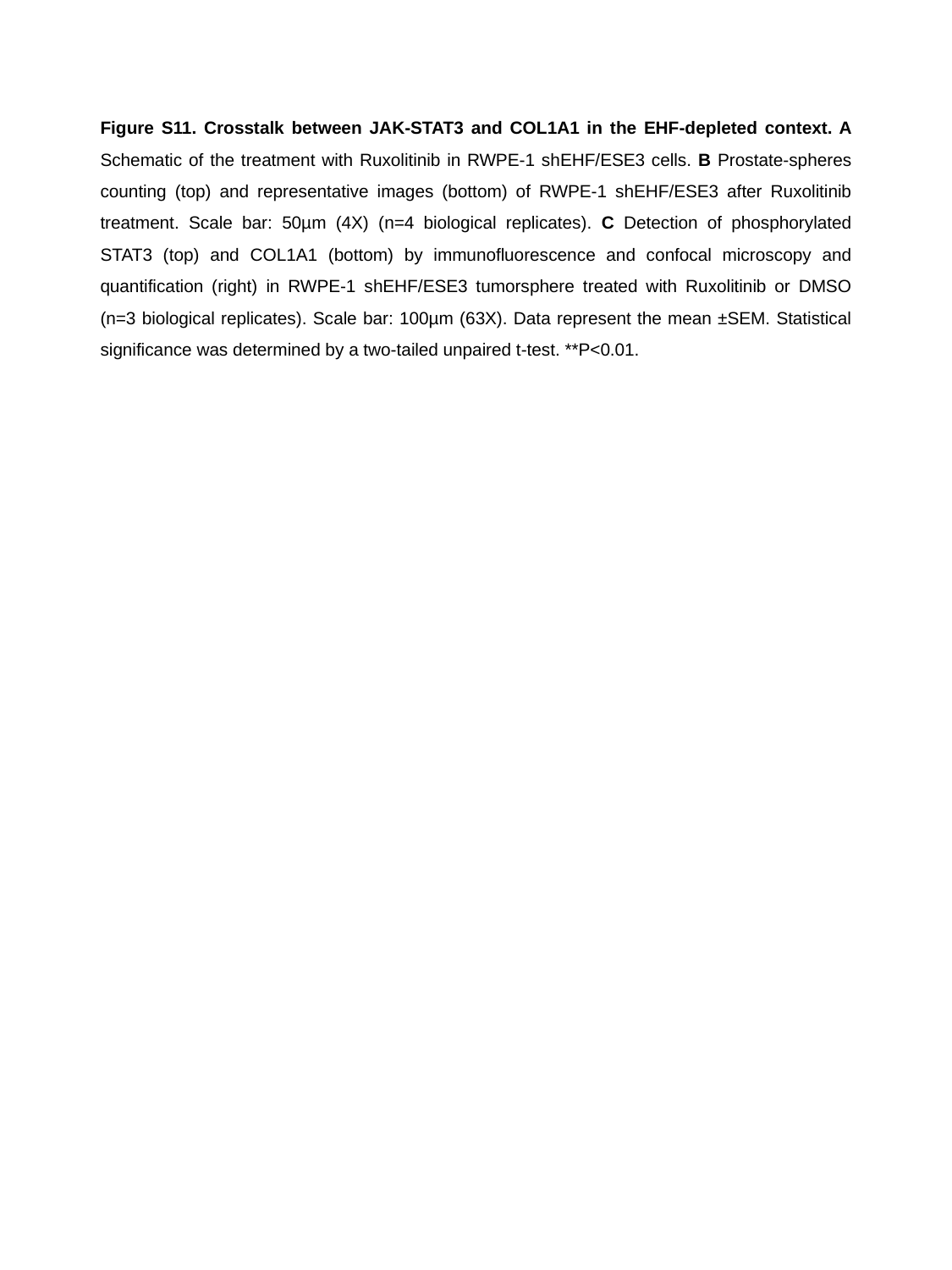

Figure S11. Crosstalk between JAK-STAT3 and COL1A1 in the EHF-depleted context. A Schematic of the treatment with Ruxolitinib in RWPE-1 shEHF/ESE3 cells. B Prostate-spheres counting (top) and representative images (bottom) of RWPE-1 shEHF/ESE3 after Ruxolitinib treatment. Scale bar: 50µm (4X) (n=4 biological replicates). C Detection of phosphorylated STAT3 (top) and COL1A1 (bottom) by immunofluorescence and confocal microscopy and quantification (right) in RWPE-1 shEHF/ESE3 tumorsphere treated with Ruxolitinib or DMSO (n=3 biological replicates). Scale bar: 100µm (63X). Data represent the mean ±SEM. Statistical significance was determined by a two-tailed unpaired t-test. **P<0.01.
